# Meiotic, genomic and evolutionary properties of crossover distribution in *Drosophila yakuba*

**DOI:** 10.1101/2022.02.10.479767

**Authors:** Nikale Pettie, Ana Llopart, Josep M. Comeron

## Abstract

The number of crossovers and their location across genomes are highly regulated during meiosis, yet the key components controlling them are fast evolving, hindering our understanding of the mechanistic causes and evolutionary consequences of changes in crossover rates. *Drosophila melanogaster* has been a model species to study meiosis for more than a century, with an available high-resolution crossover map that is, nonetheless, missing for closely related species, thus preventing evolutionary context. Here, we applied a novel and highly efficient approach to generate whole-genome high-resolution crossover maps in *D. yakuba* and tackle multiple questions that benefit from being addressed collectively within an appropriate phylogenetic framework, in our case the *D. melanogaster* species subgroup. The genotyping of more than 1,600 individual meiotic events allowed us to identify several key distinct properties relative to *D. melanogaster*. We show that together with higher crossover rates than *D. melanogaster*, *D. yakuba* has a stronger centromere effect and stronger crossover assurance than any *Drosophila* species analyzed to date. We also report the presence of an active crossover-associated meiotic drive mechanism for the *X* chromosome that results in the preferential inclusion in oocytes of chromatids with crossovers. Our evolutionary and genomic analyses suggest that the genome-wide landscape of crossover rates in *D. yakuba* has been fairly stable and captures a significant signal of the ancestral crossover landscape for the whole *D. melanogaster* subgroup, even informative for the *D. melanogaster* lineage. Contemporary crossover rates in *D. melanogaster*, on the other hand, do not recapitulate ancestral crossovers landscapes. As a result, the temporal stability of crossover landscapes observed in *D. yakuba* makes this species an ideal system for applying population genetic models of selection and linkage, given that these models assume temporal constancy in linkage effects. Our studies emphasize the importance of generating multiple high-resolution crossover rate maps within a coherent phylogenetic context to broaden our understanding of crossover control during meiosis and to improve studies on the evolutionary consequences of variable crossover rates across genomes and time.

**AUTHOR SUMMARY:** Meiotic recombination is a fundamental cellular process required for the formation of gametes in most eukaryotic organisms. Recombination also plays a fundamental role in evolution, increasing rates of adaptation. Paradoxically for such an essential process, key components evolve fast, and model organisms differ vastly in number of recombination events and distribution across chromosomes. This variation has limited our understanding of how recombination is controlled or why it can vary according to environmental cues. *D. melanogaster* has been a model species to study meiosis and recombination for more than a century and, in this study, we applied a new and highly efficient whole-genome genotyping scheme to identify recombination properties for a closely species, *D. yakuba*, thus providing an informative counterpoint and needed phylogenetic context. Our results describe important differences relative to *D. melanogaster,* including the first description of a mechanism favoring the segregation of recombinant chromatids to the functional egg pole under benign conditions. We argue that such a mechanism may also be active in other species. Moreover, our studies suggest that *D. yakuba* has maintained its recombination landscape for a longer time than *D. melanogaster,* indicating that the former species may be more adequate for evolutionary analyses. Combined, we show the importance of tackling the current siloed approach that focuses on model species with a more comprehensive analysis that requires the inclusion of closely related species when studying the causes and consequences of recombination variation.

## INTRODUCTION

Meiotic recombination is an essential cellular and evolutionary process. During meiosis, double strand breaks (DSBs) are typically repaired by reciprocal exchange between homologous chromosomes that result in either crossovers or non-crossover gene conversion events [1-6]. Crossovers and the cross-connections they form between non-sister chromatids (chiasmata) are essential to signal proper chromosome segregation during the first meiotic division in many species [7, 8]. Meiotic recombination and crossovers also play critical roles in evolution, directly generating new combinations of alleles and, indirectly, increasing the level of genetic variation within species and the rate of adaptations [9-15]. Given the important dual role of meiotic recombination and crossing-over, it is not surprising that both processes are present in the vast majority of eukaryotic organisms.

Crossover number and distribution are critically regulated, with three phenomena playing key and likely interrelated roles: *centromere effect*, *crossover interference* and *crossover assurance*. The centromere effect and crossover interference influence crossover patterning along chromosomes while crossover assurance plays a role in the distribution of crossovers among tetrads or bivalents (reviewed in [16, 17]). The centromere effect describes the classic observation in *Drosophila* that crossovers do not form in regions near the centromeres [18-24], a phenomenon that has also been observed in many other organisms such as *Saccharomyces cerevisiae*, *Arabidopsis thaliana*, and humans [25-29]. Although the causes of the centromere effect are not yet fully understood, it seems to be influenced by the distance from the centromere and highly repetitive heterochromatin [17, 20, 23, 24, 30]. A similar but weaker pattern is also observed near the telomeres in *D. melanogaster* [17, 18, 24]. The extent of the centromere effect varies between species, even among *Drosophila*. For instance, it is weaker in *D. simulans* than in *D. melanogaster*, and even weaker—if at all present—in *D. mauritiana* based on crossover data using P-elements as markers [31]. Studies in the sister species *D. pseudoobscura* and *D. persimilis* also show a tendency towards detectable but weaker centromere effect than in *D. melanogaster* [32-34]. Notably, all these *Drosophila* species with weaker centromere effect than *D. melanogaster* have higher crossover rates and longer genetic maps.

Crossover interference occurs when one crossover inhibits the formation of a second crossover in neighboring regions and therefore results in fewer and more distant double crossovers than expected if the formation of each crossover along a chromosome arm was independent [35, 36]. This phenomenon was first identified in *D. melanogaster* [35-41] and has been now observed in many other, but not all, organisms [28, 42-49]. Interestingly, two species with no apparent crossover interference (fission yeast and *Aspergillus nidulans*) also lack a synaptonemal complex [50-52]. In support for a direct role of the synaptonemal complex in crossover interference, mutants for the synaptonemal complex protein ZIP1 in *S. cerevisiae* lack interference [53]. On the other hand, the complete synaptonemal complex is not required for interference in *D. melanogaster* [40]. Worth noting, the centromere effect and crossover interference could not be detected in *D. melanogaster Blm* (Bloom syndrome helicase) mutants, suggesting a likely unifying pathway for crossover patterning across chromosomes [37].

Crossover assurance or “obligate chiasma” is the observation that each tetrad or pair of homologous chromosomes undergoes at least one crossover, irrespective of chromosome length or total number of crossovers genome-wide [16, 40, 54]. Because crossover formation is important for proper chromosome segregation at meiosis I, crossover assurance is expected to be under critical and tight regulatory control. As expected, most species with limited number of crossovers per meiosis show some degree of crossover assurance thus reducing the chance of tetrads with zero crossovers. *C. elegans* is an extreme case of absolute crossover assurance, with a percentage of tetrads with no crossovers (*E*_0_ tetrads) representing less than 1% [55]. Nevertheless, crossover formation is not always required for faithful segregation of homologs. In *Drosophila*, as in most diptera, males show faithful chromosome segregation despite no crossing over during meiosis [56-58]. Females also show proper achiasmate chromosome segregation of the small (dot) chromosome (also known as Muller F element and chromosome 4 in the *D. melanogaster* subgroup) through alternative mechanisms now known to involve centromeres and heterochromatic DNA [59-63]. A deep understanding of the different mechanisms controlling meiosis, therefore, benefits from quantifying *E*_0_ and more generally the full series of *E_r_* tetrad classes (where *r* indicates the number of crossovers in a tetrad).

The direct quantification of *E*_0_ tetrads is, however, only possible in systems where the four products of meiosis are recovered. In species where only one of the four chromatids resulting from meiosis can be analyzed (as in most species including *Drosophila*), *E_r_* can be estimated based on the fraction of products with different number of crossovers under specific meiotic models. The first and most frequently used model to estimate *E_r_* was proposed by Weinstein (1936) under the assumptions of no crossover interference, no chromatid interference, random segregation of chromatids into haploid gametes and equal viability among all possible meiotic products ([64, 65]; see Methods for details).

In *D. melanogaster*, estimates of *E*_0_ (based on Weinsten’s model unless noted) vary between studies, genotypes and conditions but the general trend suggests that crossover assurance in female meiosis is incomplete (*E*_0_ > 0; see [17] and references therein). Using Whole-Genome Sequencing (WGS), Miller *et al*. (2016) [41] calculated an overall *E*_0_ of 0.112, with similar values for the *X* chromosome and autosomes whereas studies of visible markers (observed fly phenotypes) and much larger sample sizes often show smaller estimates of *E*_0_ for the *X* chromosome than for autosomes [16, 30, 41, 66]. In all, complete or absolute crossover assurance has not been reported for any chromosome of *D. melanogaster* wild-type flies. Interestingly, studies in *D. virilis* show smaller *E*_0_ compared to *D. melanogaster* [67, 68], a result that has been proposed to be consequence of the higher rate of recombination and longer genetic map [67].

Insights into the distribution of crossover rates across genomes also provides an excellent opportunity for testing hypotheses concerning evolutionary processes. A variety of models of selection predict that variation in recombination rates has consequences on rates of evolution and levels of variation within species (diversity). These models rely on the notion that selection acting at a locus or nucleotide site causes population dynamics at neighboring genomic regions that can be understood as a reduction in local *N_e_* relative to a case without selection [69-77]. Notably, this phenomenon results from either positive or negative selection [69, 70, 74, 77-83]. Although the precise effects of selection and linkage on *N_e_* depend on numerous factors, all these models predict a positive association between intragenomic variation in crossover rates and both the efficacy of selection (which is influenced by the population-scaled selection coefficient; *N_e_* × *s*) and the level of neutral intraspecific variation or diversity (which is influenced by the population-scaled mutation rate; *N_e_* × µ) (reviewed in [72, 73, 84]). Whereas the predicted effect of variation in crossover rates on neutral diversity has been observed in many species ([72, 73, 84, 85]; although see [86]), studies in humans, *Drosophila* or yeast have provided mixed support for the prediction that variation in crossover rates across genomes modulates the efficacy of selection as estimated from either codon usage bias or rates of protein evolution (see [87, 88] and references therein).

In many species, synonymous codons are not used in equal frequencies. This observation has been proposed to be the result—at least partially—of weak selection favoring a subset of codons that increase translational speed and accuracy, particularly in highly expressed genes (see [89-92] and references therein). This model directly predicts a positive association between crossover rates (through increased efficacy of selection) and the degree of bias towards preferred codons (CUB) across genomes. In *D. melanogaster,* initial studies of variation in crossover rates and CUB provided some support for models of weak selection and linkage [93, 94] but later studies using larger data sets reported different trends for the *X* chromosome and autosomes, and instances of a negative association between recombination rates and CUB [95-98]. These unexpected results for CUB in *D. melanogaster* are difficult to reconcile with models of selection and linkage, given that either demographic effects or the inclusion of a fraction of mutations under strong selection [99, 100] would predict a weakly positive or no association between crossover rates and CUB, but not a significantly negative one. At the same time, analyses of the rate of protein evolution that include data from the *D. melanogaster* lineage show rates that are not (or only very weakly negatively) associated with crosover rates after excluding genes putatively under positive selection. Analyses focusing on genes likely under positive selection for amino acid changes show either no significant or a weakly significant positive association between crossover rates and rates of protein evolution [98, 101-105]. Studies in the *D. pseudoobscura* subgroup show no correlation between variation in crossover rates and rates of protein evolution [32].

The causes for the apparent weak support for models of selection and linkage when applied to the efficacy of selection can be multiple, including limitations when capturing potential demographic events [73, 106-108]. Most of these previous studies have also used low-resolution crossover maps, which is significant because the effect of variation in crossover rates on *N_e_* across genomes has been shown to be best captured when using fine-scale crossover data [32, 34, 80, 83, 109]. Additionally, there are potential biases associated with maps based on visible markers or P-elements with reporter genes, resulting from combinations of markers with different viability or from the presence of TEs, such as P-elements, that can directly affect recombination frequencies near insertion sites [110-112]. The use of WGS-based high-resolution crossover maps, however, eliminates such limitations [113-119].

Another caveat to most of these previous studies is the frequent assumption that crossover maps observed today capture, to a significant degree, ancestral crossover rates despite evidence that the rate and genomic distribution of crossovers are fast-evolving, with differences between distant as well as closely related species [32, 113, 120-130].Temporal changes in recombination landscapes would decouple our contemporary measures of recombination rates (and contemporary *N_e_*) from the long-term *N_e_* influencing efficacy of selection, thus reducing the genomic correlations predicted by models ([73, 86, 107] and refs therein). These temporal changes in recombination landscapes and local *N_e_* can be understood as similar to those proposed after demographic events but with the additional challenge that different genomic regions will follow different non-equilibrium conditions and dynamics whereas demographic changes act genome-wide.

In this study, we apply a novel, highly-efficient, WGS approach to generate a high-resolution crossover map in *D. yakuba* and tackle multiple questions that benefit from being addressed collectively and within a phylogenetic framework. These questions range from the patterns of crossover distribution across genomes and between chromatids, to the relationship between crossover rates and repetitive elements and short motifs, and the evolutionary consequences of crossover rate variation across genomes and time. Our data in *D. yakuba* can be directly compared with those of the related species *D. melanogaster*, a model system to study crossover and meiosis for a century for which high-resolution crossover maps already exist [41, 113]. Importantly, our new crossover landscape in *D. yakuba* also allows an initial study of the evolutionary effects of variation in recombination landscapes within the framework of the *D. melanogaster* subgroup, and specifically test the hypothesis that the lack of general support for models of selection and linkage in the *D. melanogaster* lineage may be the consequence of recent changes in the distribution of crossover rates across the *D. melanogaster* genome [98, 131].

## RESULTS

### A high resolution crossover map in *D. yakuba* using dual SNP-barcode genotyping

To generate a whole-genome high-resolution crossover map and genotype a high number of F_2_ flies with Illumina WGS, we used a new highy efficient dual-barcoding genotyping scheme (**Figure 1**). This method involves the analysis of multiple crosses between different parental lines and allows combining multiple F_2_ individuals (one per cross) per individual library barcode-sequence to later ‘demultiplex’ them bioinformatically based on genetic barcodes (diagnostic SNPs). These diagnostic SNPs are singletons, variants unique to a single line used in the study, and every Illumina read containing one can be assigned to a single parental genome and cross. Furthermore, before genotyping F_2_ individuals we improved the *D. yakuba* reference genome using PacBio long-read sequencing and then obtained high-quality sequences of all parental lines used in the crosses, with an average coverage across all chromosomes of 125× (see Materials and Methods and Supplemental Materials and Methods). Each parental line had more than 90,000 diagnostic SNPs, about one every 1.3 kb across the genome, and were used for the dual-barcoding method as well as to identify the genomic location of crossovers.

**Figure 1.**
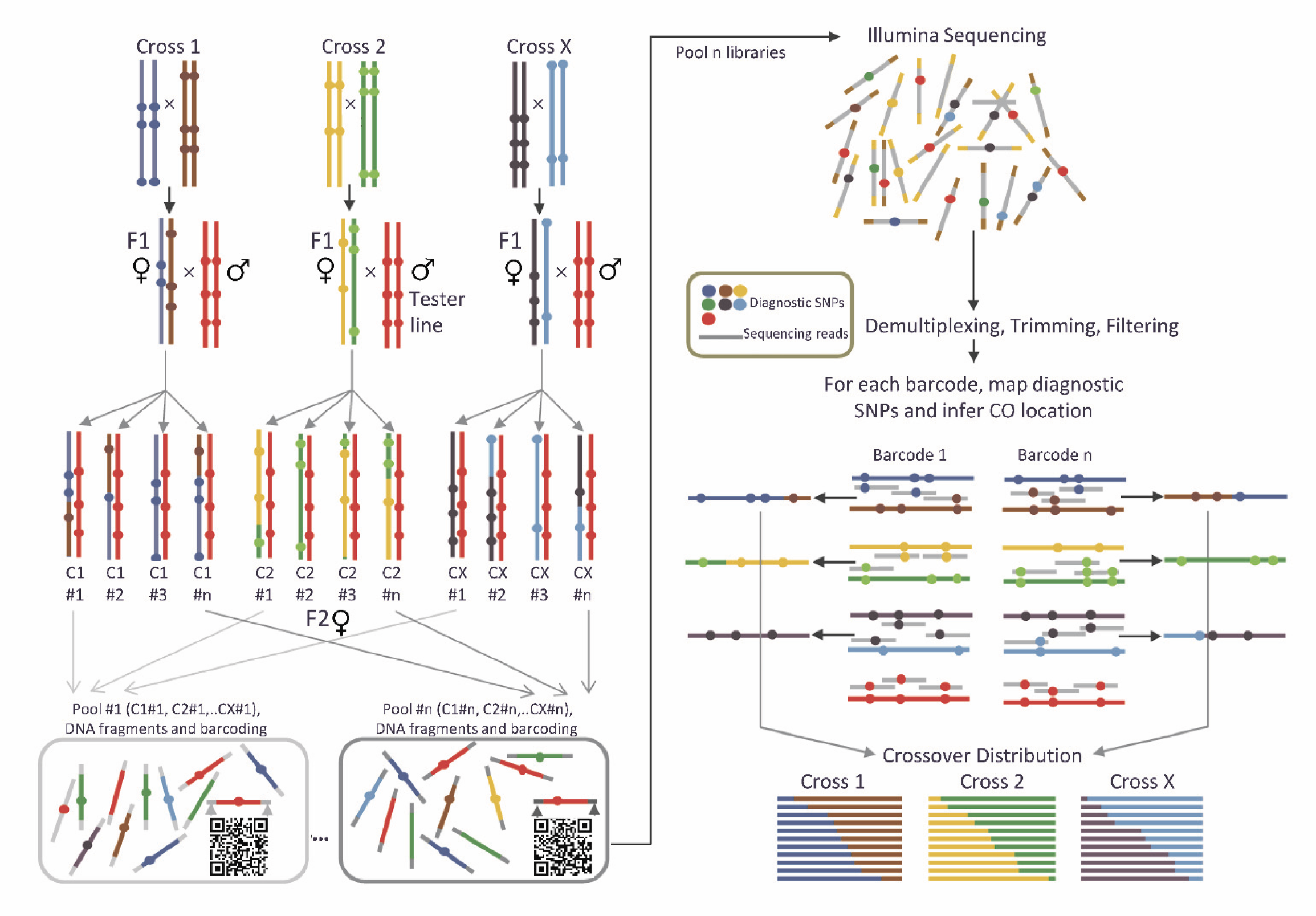
Dual-barcoding genotyping method used to obtain crossover rates. Diagnostic SNPs are used as genetic barcodes and allow for the pooling of several F_2_ individuals from different crosses for a given sequence barcode (left panel). The combination of genetic and sequence barcodes ensures efficient genotyping of multiple individuals and accurate crossover localization along chromosome arms (right panel). Diagnostic, or strain-specific SNPs, are singletons for the complete set of genotypes used in the study, including the tester line.

More than 1,600 F_2_ individuals from three crosses of wild-type lines were genotyped to generate this first high-resolution genetic map in *D. yakuba* **(Table 1).** One cross (*Sn20* x *Sn17*) showed a heterozygous chromosomal inversion on *2R* (see Supplemental Materials and Methods) and, therefore, no crosover data from that chromosome arm were used in the analyses. In all, we recovered a total of 5,273 crossover events distributed across all chromosome arms except the dot chromosome(**Figure 2A**, **Table 1** and **Supplemental Table 1**). The absence of crossovers on this small chromosome 4 despite the presence of diagnostic SNPs is consistent with results in other *Drosophila* species [37, 41, 113, 132, 133].

**Figure 2.**
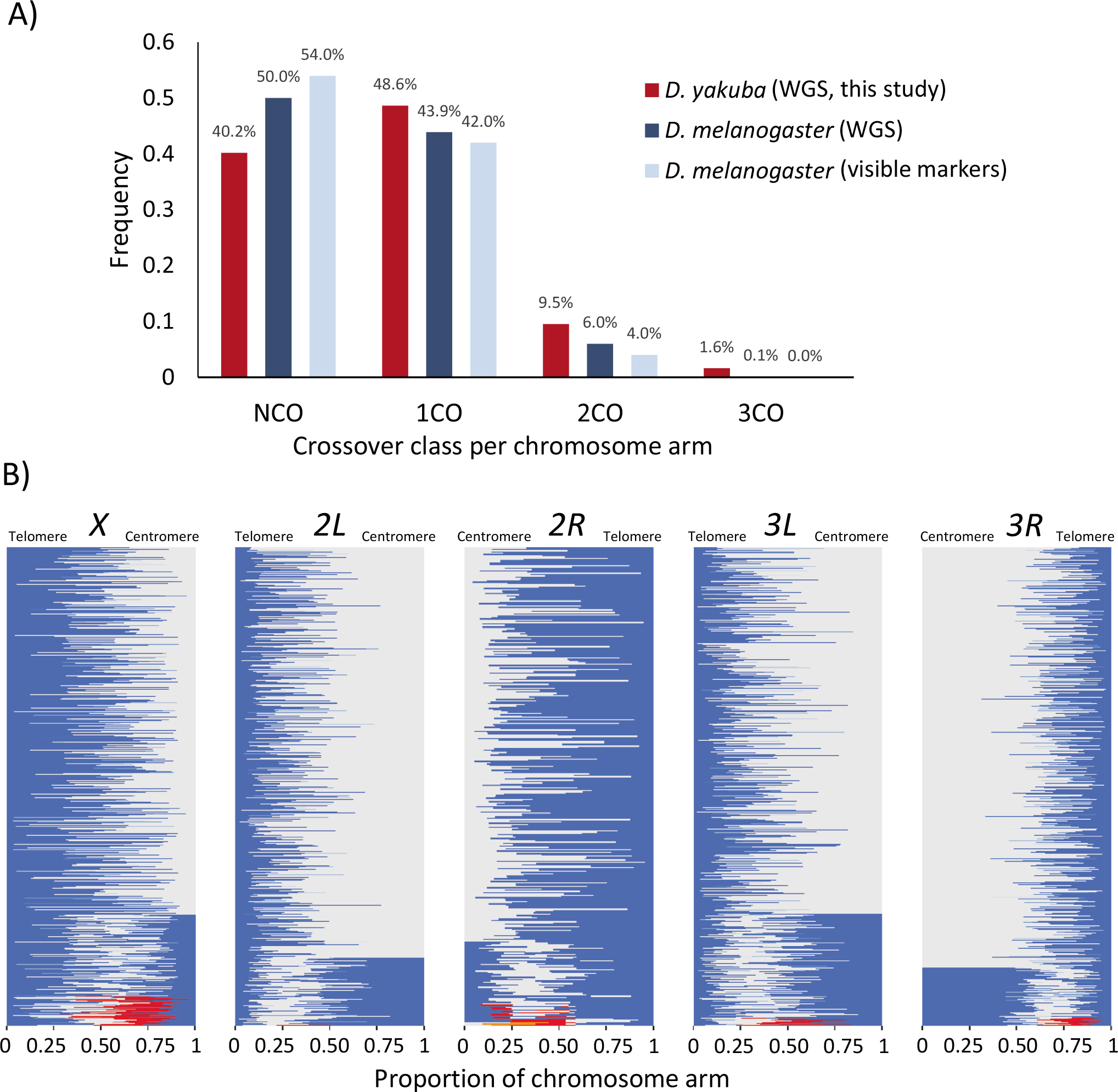
Number and distribution of crossovers. **A)** Observed crossovers per chromatid in *D. yakuba* and *D. melanogaster*. *D. melanogaster* frequencies from whole genome sequencing (WGS) from Miller *et al.* (2016) [41], and those from visible markers adapted from Hatkevich *et al*. (2017), Baker and Carpenter (1972), and Parry (1973) [37, 212-213]. Note that studies based on visible markers are based on much larger sample sizes but, at the same time, may underestimate crossover events due to limited density of markers along chromosome arms. NCO: non-crossover, 1CO: single crossover, 2CO: double crossover, 3CO: triple crossover, 4CO: quadruple crossovers per chromosome arm. **B)** Relative location of crossovers along chromosome arms in *D. yakuba*. Each horizontal line represents a chromosome with one or more crossovers. A change in color represents a crossover event, starting from the telomere (blue). Chromosome arms have been ordered based on crossover class for better visualization, from 1CO (top) to 4 CO (bottom). Number of chromatids analyzed (*n): X*: *n* = 1,622; *2L*: *n* = 1,661; *2R*: *n* = 841; *3L*: *n* = 1,701; *3R*: *n* = 1,635.

**Table 1.** Observed number of meiotic events in *D. yakuba*.

| CO Class<sup>1</sup> | Chromosome |  |  |  |  | Total |
| --- | --- | --- | --- | --- | --- | --- |
|  | *X* | *2L* | *2R*<sup>2</sup> | *3L* | *3R* |  |
| NCO | 447 | 835 | 463 | 694 | 683 | 3122 |
| 1CO | 901 | 708 | 312 | 781 | 829 | 3531 |
| 2CO | 200 | 116 | 50 | 209 | 107 | 682 |
| 3CO | 74 | 2 | 14 | 17 | 15 | 122 |
| 4CO | 0 | 0 | 2 | 0 | 1 | 3 |
| Chromatids | 1622 | 1661 | 841 | 1701 | 1635 | 7460 |

<sup>1</sup>NCO: zero crossover, 1CO: single crossover, 2CO: double crossover, 3CO: triple crossover, 4CO: 4 crossovers in a single chromatid. <sup>2</sup>The Sn20 x Sn17 cross is heterozygous for an inversion on *2R* (Supplemental Materials and Methods) and, therefore, no crossovers were analyzed for this chromosome arm. For comparison, meiotic events in *D. melanogaster* analyzed in this study are shown in Supplemental Table 11.

There are fewer non-crossovers (NCOs), more single crossovers (1COs) and, especially, more double crossovers (2COs) in *D. yakuba* than in *D*. *melanogaster* (**Figure 2A**). This trend is observed across all chromosomes examined, with the *X* chromosome of *D. yakuba* showing the largest increase (64%) in crossovers relative to *D. melanogaster*. Notably, we observed multiple chromatids with triple crossovers (3CO) and 0.05% (three) cases with four crossovers (4CO).

We observe some variation in the average number of crossovers per gamete among the three crosses analyzed (from 3.25 to 3.76). Nevertheless, there is a much larger degree of variability in the distribution of crossovers per gamete, and an ANOVA test indictes that cross plays a minor role in the number of crossovers per gamete (ANOVA test, partial η^2^ = 0.015; **Supplemental Figure 1**). Additionally, there is no evidence of an interchromosomal effect [39, 134, 135] caused by the presence of a heterozygous inversion on chromosome *2R* in the *Sn20* x *Sn17* cross given that the number of crossovers in freely recombining chromosome arms for this cross is not higher than in the other two crosses (**Supplemental Table 1).** In agreement, an ANOVA test shows that the presence of the heterozygous *2R* inversion has no detectable effect on the number of crossovers per gamete on freely recombining chromosome arms (partial η^2^ = 0.001; *P* > 0.05).

We, therefore, performed all analyses combining the results of the three crosses unless noted. In *D. yakuba*, the average number of crossovers per chromosome arm is 0.94, 0.57, 0.55, 0.73 and 0.67, for the *X* chromosome and chromosome arms *2L*, *2R*, *3L* and *3R*, respectively. Overall, our study shows an average of 3.46 crossovers per viable meiotic product relative to 2.76 in *D. melanogaster*. The total genetic map length for *D. yakuba* obtained in our crosses is 339.31 cM, about 25% longer than that for *D. melanogaster* [41, 113, 136]. Average crossover rates are highest for chromosome *X* (4.07 cM/Mb) and range between 2.00 and 2.76 cM/Mb for autosomal arms (2.00, 2.47, 2.51 and 2.76 cM/Mb for chromosome arms *3R*, 2*L*, 2*R* and *3L*, respectively). Although recombination maps generated for the different crosses show some differences, the main trends of intrachromosomal variation are consistent among crosses for all chromosome arms (**Figure 3**).

**Figure 3.**
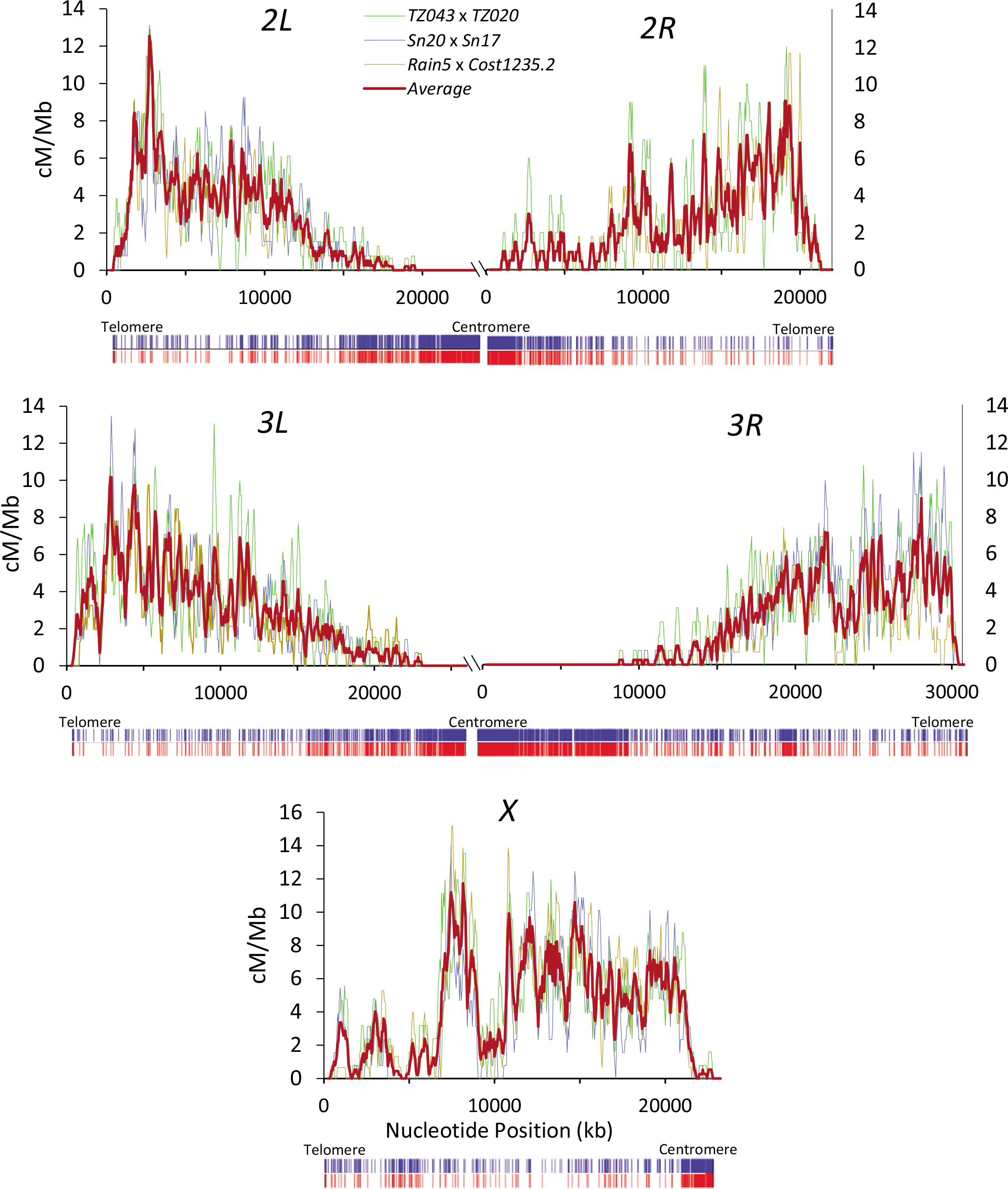
High-resolution crossover maps for *D. yakuba*. Crossover rates (cM/Mb) are shown along chromosome arms from three different crosses as well as the average (thick red line). Maps are shown for overlapping 250-kb windows with increments of 50kb. Below each chromosome, vertical blue and red bars indicate the presence of transposable and INE-1 elements, respectively. The scale for the different chromosome arms is equivalent and differences in figure size capture differences in chromosome arm length.

### Centromere and telomere effects

Similar to other *Drosophila* species, *D. yakuba* shows a noticeable centromere effect (**Figure 2B** and **Figure 3**). Using the standard approach of comparing observed and expected crossover numbers within a centromere-proximal region of a predetermined size (e.g., one-third of the chromosome arm [41]), we identified significant centromere effects on the autosomes but not on the *X* chromosome (**Figure 3** and **Supplemental Table 2**).

To compare the magnitude of centromere effects among chromosome arms and species, we used a more quantitative method that allows estimating the actual size of centromere-proximal regions with a significant deficit in crossovers relative to expectations (see Material and Methods). Using this method, we identified a centromere effect in all chromosomes of *D. yakuba* (**Figure 4** and **Supplemental Table 3)**. On autosomes, the proportion of the chromosome that experiences the centromere effect is much larger in *D. yakuba* than in *D. melanogaster*, with chromosome arms *2L* and *3R* in *D. yakuba* showing an almost two-fold increase in the size of the affected genomic region relative to *D. melanogaster*. Chromosome arm *3R* in *D. yakuba*, in particular, stands out with 47% and 52% of the arm showing a detectable centromere effect at *P* = 1×10^-6^ and *P* = 0.01 significance levels, respectively. On the other hand, the centromere effect is weakest on the *X* chromosome of *D. yakuba*, as it was also reported for *D. melanogaster* [21, 41, 113, 137], and both species show a similar proportion of the chromosome influenced by the centromere effect (12% and 13% of the chromosome arm at *P* = 1×10^-6^ and *P* = 0.01 significance levels, respectively).

**Figure 4.**
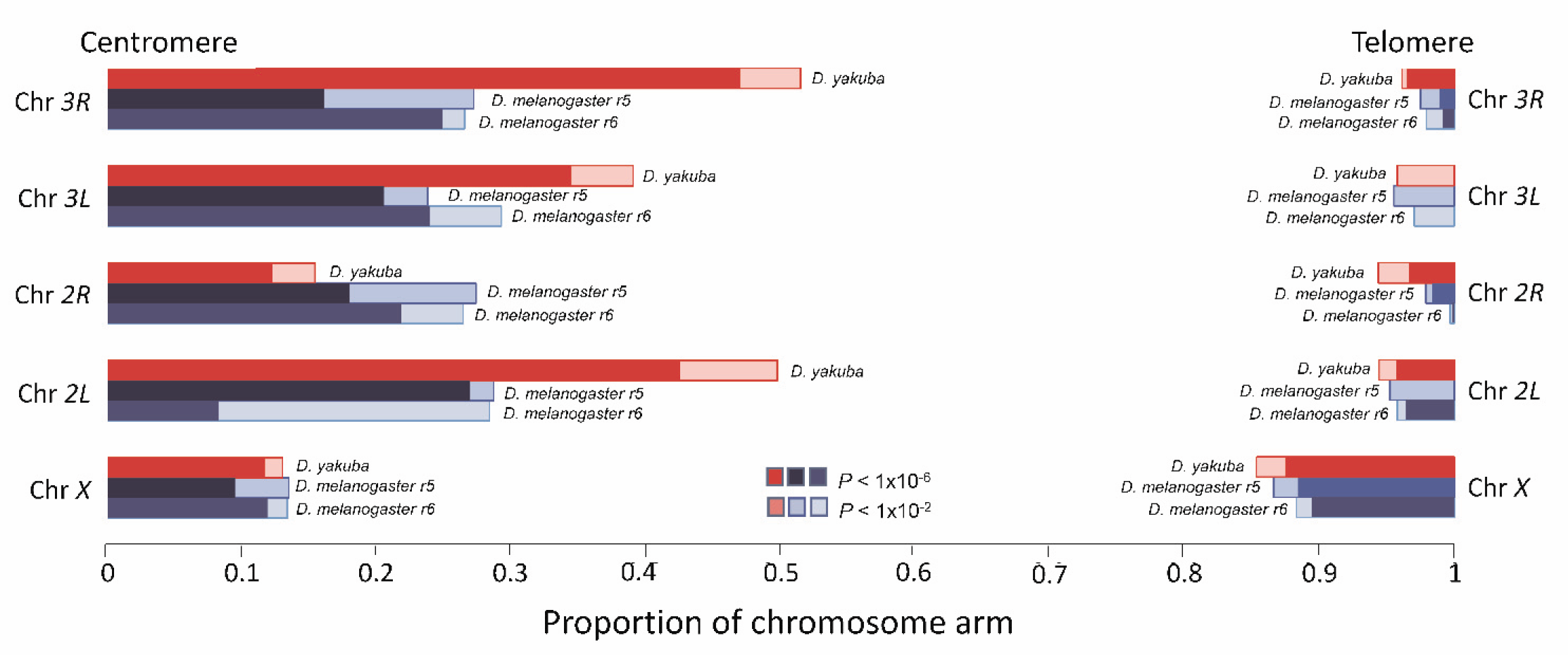
Centromere and telomere effects for different chromosome arms of *D. yakuba* and *D. melanogaster*. For *D. melanogaster* results are shown for genome r5.3 and r6 assemblies. The proportion of the chromosome with significant reduction in crossovers is based on the study of a 1Mb- sliding window with increments of 100kb. Two levels of significance are shown.

To examine the telomere effect, we first used the standard methodology applied to the telomere-proximal one-third of the chromosome arm, and identified a significant deficit of crossovers on the *D. yakuba X* chromosome but not on any of the autosomal arms, reversing the observation for the centromere effect (**Supplemental Table 2**). The use of the quantitative method to identify the proportion of the chromosome affected by the telomere effect in *D. yakuba* confirms a much smaller effect in all chromosome arms compared to the centromere effect (**Figure 4** and **Supplemental Table 3)**, a trend that is also observed in *D. melanogaster* [21, 31, 41, 113]. Our results for *D. yakuba* are also consistent with the observation in *D. melanogaster* that the *X* chromosome has the strongest telomere effect (12% and 11% of the chromosome for *D. yakuba* and *D. melanogaster*, respectively, at *P* = 1×10^-6^ significance level). Chromosome arm *3L*, on the other hand, shows the weakest telomere effect in *D. yakuba* (as it is also the case in *D. melanogaster*), with <3% and 4% of the chromosome arm affected at *P* = 1×10^-6^ and *P* = 0.01 significance levels, respectively.

### Satellite repeats and the centromere effect

Although the mechanisms controlling the magnitude of the centromere effect are not fully understood, analyses in *D. melanogaster* suggest that the distance to the centromere plays a role, with increasing suppression of crossovers in peri-centromeric regions when centromeric heterochromatin is removed [137, 138]. At the same time, centromeres are heavly enriched in satellite DNA sequences [137, 139-141]. We, therefore, investigated whether the greater centromere effect in *D. yakuba* relative to *D. melanogaster* could be related to reduced centromeric heterochromtin and centromere size (see Supplemental Materials and Methods). The complete sequence of centromeres in *D. yakuba* is yet unknown and we, therefore, estimated the amount of satellite DNA in heterochromatic regions as an indirect measure of centromere size. We used PacBio reads that do and do not align to our updated euchromatic *D. yakuba* assembly as proxy for euchromatic and heterochromatic sequences, respectively, and characterized satellite DNA for simple repeats (k-mers) using k-Seek [142]. For *D. melanogaster*, PacBio reads were aligned to both the r5.3 and r6 releases [143], with similar conclusions and, therefore, only the r6 results will be discussed here. In agreement with previous studies indicating a high turnover rate in satellite repeats between closely related *Drosophila* species [139-141], we find that satellites that are most enriched in the centromeres of *D. melanogaster* are mostly absent in the heterochromatic reads of *D. yakuba* (**Supplemental Table 4**). Overall, heterochromatic reads in *D. yakuba* show a lower abundance (more than 20-fold) of satellite repeats than those in *D. melanogaster* and this tendency remains when considering the difference observed in euchromatic reads, which is only 4-fold (**Supplemental Table 5**). Tentatively, these results support the notion that centromeres in *D. yakuba* are likely shorter than those in *D. melanogaster*, and shorter centromeres in *D. yakuba* may play a role in the stronger centromere effect relative to *D. melanogaster*.

### Crossover interference in *D. yakuba*

In this study we recovered a total of 673 chromatids with 2 crossovers from a single meiosis event, thus allowing for a detailed study of crossover interference. Genome-wide, the average inter-crossover distance (ICD) in 2CO chromatids is shorter than that expected if crossovers were randomly distributed across the genome (7.5 vs 8.4Mb, respectively), with a minimum ICD of 820 Kb. The average ICD in 3CO chromatids is also shorter than random expectations (5.9 vs 6.3 Mb, respectively). Indeed, the standard approach of studying crossover interference by randomly positioning crossover events across the genome would suggest that crossover interference is completely absent in *D. yakuba*.

However, given the strong centromere/telomere effects, we quantified crossover interference based on expectations from the distribution of crossovers in 1CO chromatids (see Material and Methods for details). The ratio of observed to expected ICD using this approach is 1.36 genome-wide, with the highest values observed for chromosome arms *2L* and *3R* (1.54 and 1.51, respectively) and the lowest values observed for the *X* chromosome and chromosome arm *2R* (1.17 and 1.08, respectively). **(Table 2)**. These analyses reveal a highly significant tendency for crossovers in 2CO chromatids to be more distant than expected (positive interference) for all chromosome arms except *2R,* with a weak, albeit significant, departure from genome-wide expectations (*P* = 0.046) that becomes highly significant when removing *2R* (*P* < 1×10^-6^).

**Table 2.**
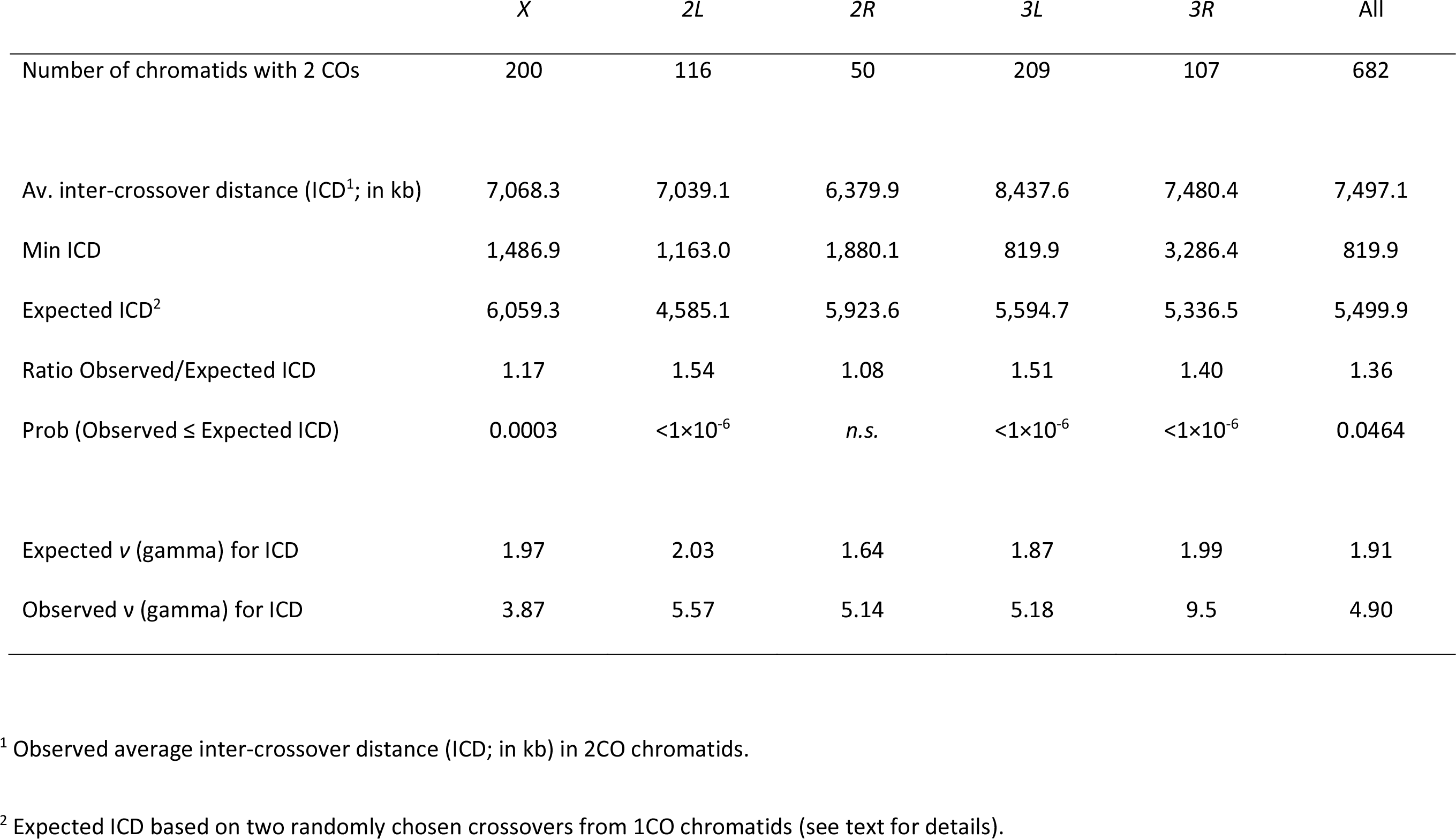
Crossover interference in *D. yakuba*

|  | *X* | *2L* | *2R* | *3L* | *3R* | All |
| --- | --- | --- | --- | --- | --- | --- |
| Number of chromatids with 2 COs | 200 | 116 | 50 | 209 | 107 | 682 |
| Av. inter-crossover distance (ICD<sup>1</sup>; in kb) | 7,068.3 | 7,039.1 | 6,379.9 | 8,437.6 | 7,480.4 | 7,497.1 |
| Min ICD | 1,486.9 | 1,163.0 | 1,880.1 | 819.9 | 3,286.4 | 819.9 |
| Expected ICD<sup>2</sup> | 6,059.3 | 4,585.1 | 5,923.6 | 5,594.7 | 5,336.5 | 5,499.9 |
| Ratio Observed/Expected ICD | 1.17 | 1.54 | 1.08 | 1.51 | 1.40 | 1.36 |
| Prob (Observed ≤ Expected ICD) | 0.0003 | <1×10<sup>-6</sup> | *n.s.* | <1×10<sup>-6</sup> | <1×10<sup>-6</sup> | 0.0464 |
| Expected $\nu$ (gamma) for ICD | 1.97 | 2.03 | 1.64 | 1.87 | 1.99 | 1.91 |
| Observed $\nu$ (gamma) for ICD | 3.87 | 5.57 | 5.14 | 5.18 | 9.5 | 4.90 |

<sup>1</sup> Observed average inter-crossover distance (ICD; in kb) in 2CO chromatids.
<sup>2</sup> Expected ICD based on two randomly chosen crossovers from 1CO chromatids (see text for details).

We also studied crossover interference based on the full distribution of ICD in 2CO chromatids and estimated the shape parameter (ν) of a gamma distribution (see **Table 2** and Materials and Methods). Estimates of ν greater than those expected under no crossover interference (which is ν = 1 under ideal conditions; see Materials and Methods) are an indication of positive interference. In *D. melanogaster*, a large-scale study of visible markers along the *X* chromosome indicates ν ∼ 5 [144]. In *D. yakuba*, estimates of ν range between 3.85 and 9.5 (for the *X* chromosome and chromosome arm *3R*, respectively) and provide additional support for a significant degree of positive crossover interference in this species.

### Tetrad analysis shows stronger crossover assurance in *D. yakuba* than in *D. melanogaster*

Tetrad analysis suggests that there are fewer tetrads with zero crossovers (*E*_0_) in *D. yakuba* than in *D. melanogaster* (**Figure 5A** and **Supplemental Table 6**). The direct application of Weinstein’s equations [64] generates a *D. yakuba* genome-wide estimate of *E_0_* very close to zero (*E_0_* = -0.005), which is compatible with absolute crossover assurance (P[*E_0_* = 0] = 0.99; see Materials and Methods for details). Autosomes show a weak but positive estimate of *E*_0_ (E*_0_* = 0.059) suggesting a stronger degree of crossover assurance than that reported for *D. melanogaster* autosomes. The *D. yakuba X* chromosome data, however, produces a negative estimate of *E_0_* ( *E_0_* = -0.202).

**Figure 5.**
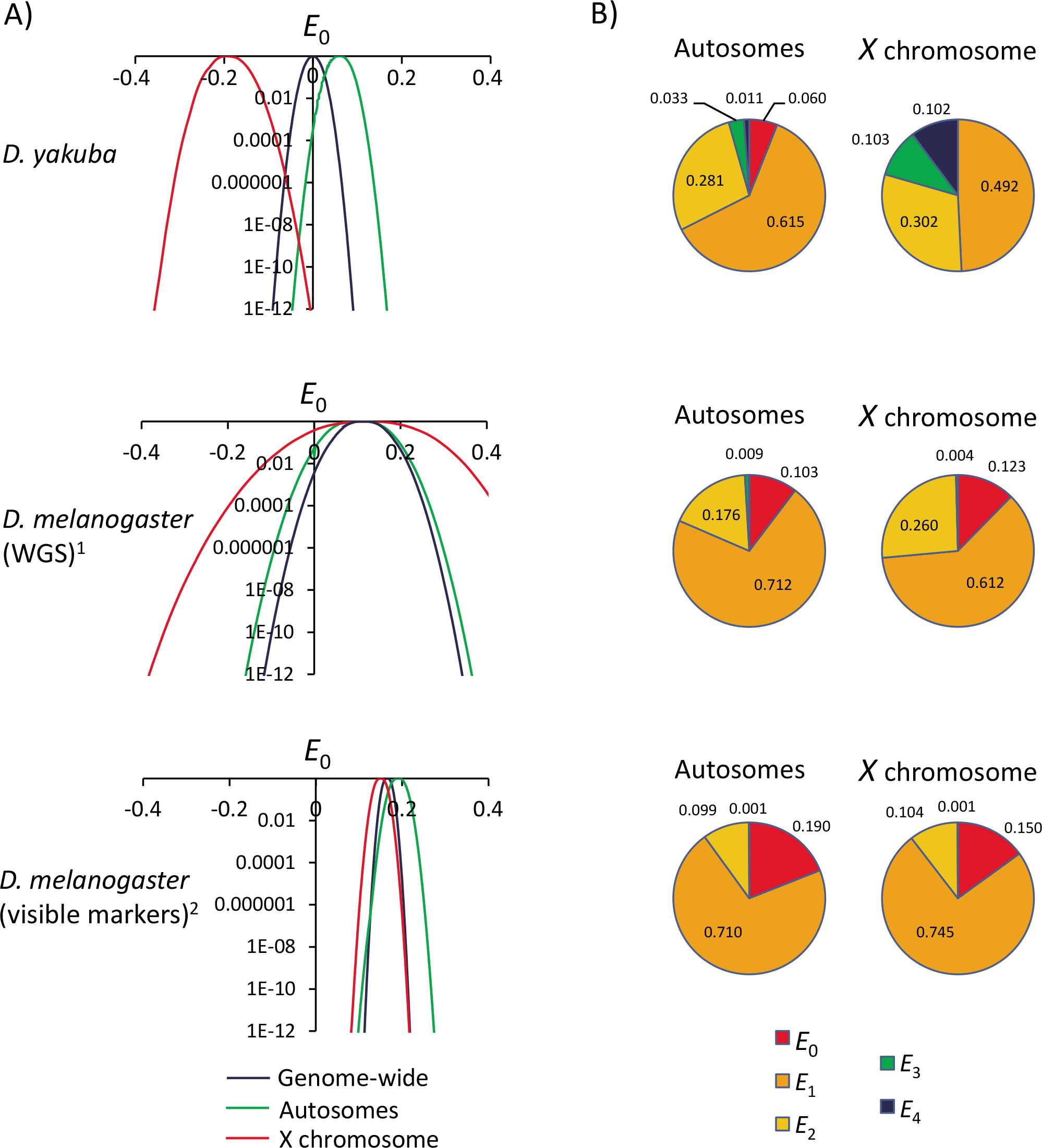
Maximum Likelihood (ML) estimates of tetrad frequencies. **A)** Estimates of *E_0_* under a ML model with unrestricted *E_r_*. Probability (y-axis) of models with variable *E*_0_ (x-axis), with *E*_r>0_ allowed to vary to provide the best fit for a given *E*_0_. ^1^ Estimates of tetrad frequency in *D. melanogaster* based on the distribution of 196 meiotic products analyzed by Miller *et al*. (2016) [41] using whole genome sequencing (WGS). ^2^ Estimates of tetrad frequency in *D. melanogaster* based on visible markers (see text). **B)** Estimates of tetrad frequencies based on ML models that constrain biologically relevant ranges for *E_r_* (*E_r_* ≥ 0). Note that the best model for the *D. yakuba X* chromosome (*E_0_* = 0) is, nonetheless, incompatible with the observed data [*P*(*E_0_* = 0) = 1.6×10^-13^] whereas all other models show a good fit (*P* > 0.99). *E*_0_: tetrads that do not undergo crossing over, *E*_1_: tetrads with 1 CO, *E*_2_: tetrads with 2 COs, *E*_3_: tetrads with 3 COs, and *E*_4_: tetrads with 4 COs.

Negative *E*_0_ can be caused by sampling errors and small sample sizes. Whereas limited sampling is hardly a limitation when using visible markers, it can become a serious drawback in studies based on WGS that are often limited to a few hundred meiotic products. For instance, a typical distribution of crossover events for *D. melanogaster* of 50% NCO, 45% 1CO and 5% 2CO chromatids predicts *E_0_* = 0.10 but random sampling could generate negative *E_0_* in 10 and 22% of the studies when the number of analyzed chromosomes is 250 and 100, respectively. Given our very large sample size of 1,622 chromatids analyzed for the *D. yakuba X* chromosome, the likelihood of obtaining our negative *E_0_* given a true *E_0_* = 0 is remarkably small (*P* < 1×10^-14^). We used a ML approach that allows adding diverse sets of rules (see Materials and Methods and [66, 145]), and explored a model that constrains all tetrad classes to biologically relevant frequencies (*E_r_* ≥ 0) (**Figure 5B**). Although the best model for the *D. yakuba X* chromosome under these conditions is *E_0_* = 0, we can, nonetheless, rule out *E_0_* ≥ 0 (P[*E_0_* ≥0] < 1.2×10^-14^).

### Analysis of viability effects

Besides statistical considerations, estimates of *E*_0_ under Weinstein’s model can generate negative values if at least one of the assumptions is violated. The most commonly argued violation is viability effects associated with specific combinations of alleles. This is often an issue when using visible markers to characterize crossovers, and has been shown to play a significant role in tetrad analyses in *D. melanogaster* [66]. Moreover, fitness effects associated with visible mutations can also alter recombination frequencies in other chromosome arms [39, 41, 134, 135, 146]. WGS genotyping methods such as ours and the use of wild type flies, on the other hand, can ameliorate such potential biases but viability effects should be still considered especially if inbred lines are being used in the crossing schemes. This is because inbreeding could cause the fixation of recessive deleterious alleles, as evidenced by the fitness decline in inbred lines documented in many species, including *Drosophila* [147-151]. In such cases, the crossing of two inbred lines has the potential to bias (either upward or downward) the relative frequency of recombinant haplotypes observed in adults (e.g., non-epistatic deleterious alleles in repulsion phase would bias upward the frequency of recombinant haplotypes whereas the fixation of compensatory mutations along a chromosome could generate opposite effects).

Our crossing and genotyping schemes are designed to reduce the likelihood of viability effects. In our crossing scheme, heterozygous F_1_ females are crossed to males from a ‘third’ (tester) line and we genotyped F_2_ females (**Figure 1**). Thus, potential deleterious alleles with egg-to-adult viability effects that became fixed due to inbreeding will likely be heterozygous in F_2_ females, limiting viability defects across the whole genome. Moreover, the inbred lines used in our study were generated by in mass brother-sister mating, which exposes mutations to both drift and selection and increases the likelihood of purging severely deleterious and non-recessive alleles relative to more extreme inbreeding methods [152, 153]. Finally, we used three different crosses involving a total of six inbred parental lines from different natural populations, making unlikely that multiple lines share similar combinations of deleterious alleles. All three crosses show the same trend of a negative unconstrained estimate of *E*_0_ for the *X* chromosome [*E_0_* of -0.163, -0.185 and -0.263]. Combined, our results and crossing schemes would predict limited viability effects altering haplotype frequencies.

To rule out viability effects as a cause of the unsual *E*_0_ values for the *X* chromosome more directly,we estimated the frequency of the four possible haplotypes (pairwise combinations of the two parental genotypes) recovered in F_2_ individuals as a function of the genetic distance across the entire chromosome. Viability effects would generate one parental haplotype to be more frequent than the other and/or one recombinant haplotype to be more frequent than the other. Our study (**Supplemental Figure 2**) shows that the two parental haplotypes are observed at equal frequencies and that the two recombinant haplotypes are also observed at equal frequencies for any given genetic distance. Moreover, the frequency of recombinant haplotypes increases with genetic distance, as predicted if all (or the great majority) of recombinants are observed in adult females (note that the frequency of the two recombinant haplotypes stays lower than 25% for ∼ 50 cM due to the presence of multiple COs in a fraction of gametes). Additionally, we obtain equivalent results for each of the three crosses analyzed. These results allow us to rule out viability effects in our recombination data and strongly hint at the presence of mechanisms that alter the segregation or transmission of chromatids for the *D. yakuba X* chromosome.

### Crossover-associated meiotic drive in *D. yakuba X* chromosome

A biological explanation for an observed deficit in chromatids with zero crossovers would be a form of meiotic drive [154] during meiosis II. Specifically, our data could be explained by meiotic drive if the sister chromatid that experienced a crossover was preferentially segregated into the oocyte nucleus relative to its nonrecombinant sister chromatid, which would be preferentially extruded to the second polar body (**Figure 6A**). As a consequence of this form of crossover-associated meiotic drive (MD­), the number of crossovers detected in the offspring would be above that expected under Mendelian segregation for any given number of crossovers during meiosis I. Note that MD_CO_ differs from the standard concept of meiotic drive because only the latter is associated with allelic genetic variants in heterozygotes [154-164].

**Figure 6.**
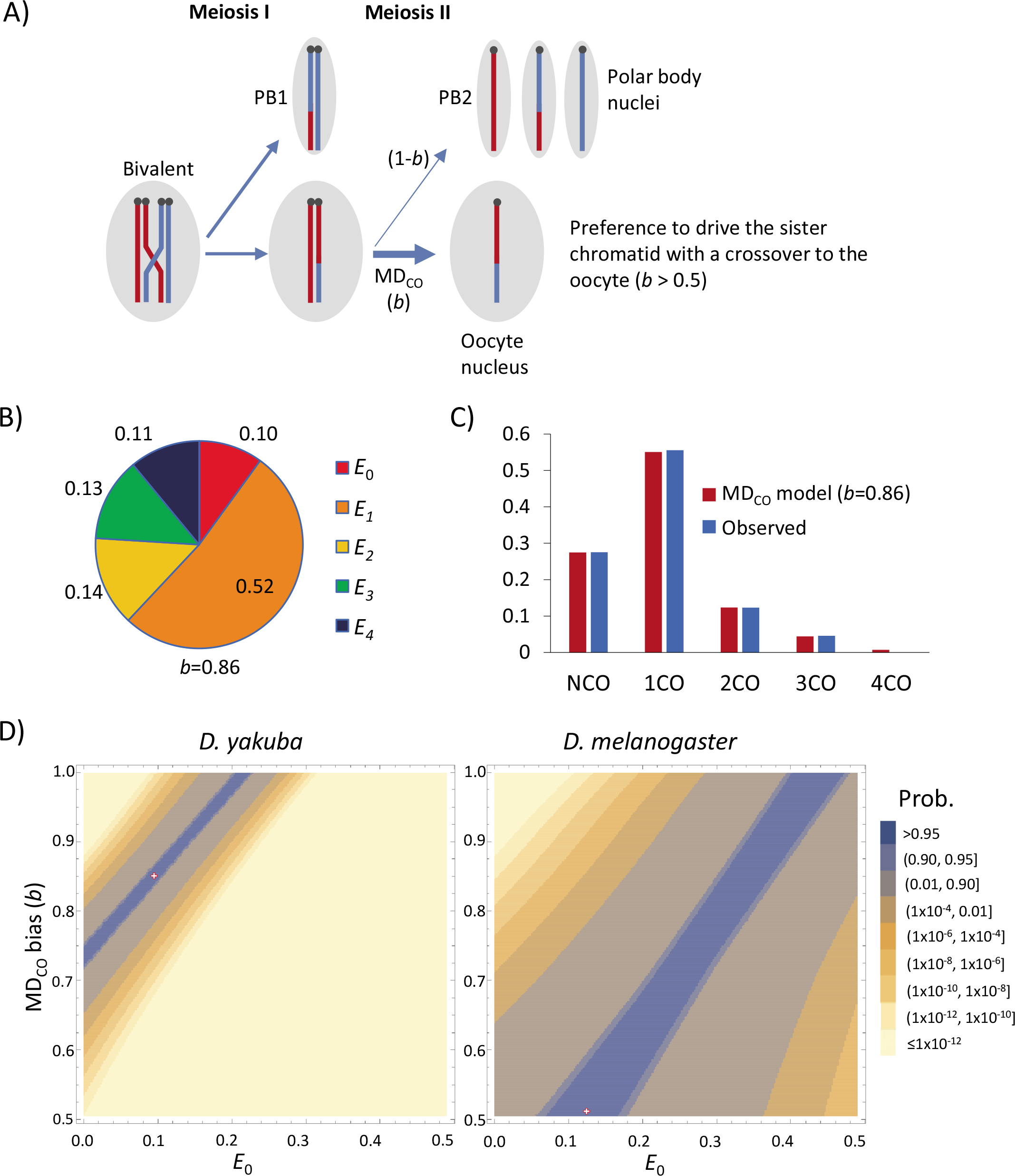
Model of tetrad frequencies with crossover associated meiotic drive (MD_CO_). **A)** MD_CO_ model where chromatids with crossovers are preferentially transmitted to the oocyte with a bias *b* (*b* > 0.5) when the sister chromatid has no crossovers. **B)** ML estimates of tetrad frequencies for the *D. yakuba X* chromosome under a MD_CO_ model, with a best fit to the *X* chromosome of *D. yakuba* when *b* = 0.86. **C)** Observed and predicted frequencies of crossover classes for chromosome *X* under this model. **D)** Joint maximum-likelihood estimates (MLE) of *E*_0_ and *b* for the *X* chromosome of *D. yakuba* (left) and *D. melanogaster* (WGS; right). The red dot represents point estimates with maximum fit to the data.

We explored a ML model including the possibility of a bias in meiosis II favoring the recovery (oocyte) of chromatids with crossovers when the sister chromatid has no crossovers. This modelling shows that allowing MD_CO_ not only improves the fit to the *X* chromosome crossover data significantly relative to a model without segregation bias (Likelihood-ratio test, LRT, *P* < 1×10^-13^) but also explains the overall distribution of crossovers accurately (*P* = 0.997) (**Figure 6B,C**). Specifically, point estimates for our *D. yakuba X* chromosome data suggest a model with *E_0_* = 0.10 and a probability of chromatids with crossovers to ultimately be present in oocyte nuclei of *b* = 0.86 instead of 0.5 when paired with sister chromatids with no crossovers. More generally, however, a MD_CO_ model fits the *D. yakuba X* chromosome data with *P* > 0.99 for a wide range of *E*_0_ and *b* values, from *E*_0_=0 (with *b*=0.73) to *E*_0_=0.22 (with *b*=1) (**Figure 6D, left**). Given these results for *D. yakuba*, we investigated the possibility of MD_CO_ for the *D. melanogaster X* chromosome for comparison (**Figure 6D, right**). This study shows that although a significant effect of MD_CO_ is not necessary to explain crossover information in this species, it is nonetheless compatible with it (*P* > 0.99) for a wide range of plausible *E*_0_ values (from 0.10 to 0.46).

We also examined the possibility of *chromatid interference* [64, 145]. Unlike MD_CO_, the average number of crossovers per offspring chromatid does not change under chromatid interference, only the distribution of crossovers among chromatids when *E_≥2_*. In this regard, *positive chromatid interference* or PCI (making the nonsister pair of chromatids involved in one crossover less likely to be involved in additional crossovers) would cause a reduction in the fraction of chromatids with extreme values of crossovers, including those with zero crossovers. For instance, an extreme case of PCI for *E_2_* would generate only chromatids with 1 crossover instead of the expected 25, 50 and 25% of chromatids with 0, 1 and 2 crossovers, respectively. Thus, although chromatid interference is rare in diploids [165], PCI could—in principle—generate the observed deficit of chromatids with zero crossovers for the *D. yakuba X* chromosome. To evaluate this possibility, we incorporated PCI into our ML model with constrained tetrad frequencies (*E_r_* ≥ 0), now allowing for a biased distribution of crossovers (*b*) favoring the use of the sister chromatid with the fewest number of previous crossovers (*b* ≥ 0.5 between sister chromatids instead of *b* = 0.5 for the random case). When applied to the *D. yakuba X* chromosome data, a PCI model is better than a model without PCI but, nonetheless, incompatible with our observations (best fit when *b* = 0.59, *P* = 1.1×10^-9^) and worse than the MD_CO_ model described above (LRT = 41.2, *P* = 1.4×10^-10^).

### Transposable elements (TEs) and crossover rates

Given the frequent deleterious effects of TE insertions, models concerning crossover rates and efficacy of selection predict that TEs would passively accumulate in regions with reduced crossover rates (reviewed in [166, 167]). In agreement, TEs tend to be more frequent in genomic regions with limited or absent crossovers in many species, including *D. melanogaster* [67, 110-112, 166, 168-170]. There is, however, an alternative explanation to this same observation which does not rely on an association between contemporary crossover rates and long-term efficacy of selection; TEs can actively reduce crossover rates via epigenetic mechanisms related to TE silencing around insertions thus creating a similar negative intra-genomic association between crossover rates and TE presence—even if crossover landscapes have changed very recently [171-177].

Consistent with previous studies [178, 179], our analysis shows that there are more TE copies (about 30% more) in the *D. yakuba* genome assembly than in *D. melanogaster* (see Supplemental Materials and Methods). Our study also shows that the overall difference in TE presence is driven by differences in copy number of INE-1 (*Drosophila* INterspersed Element), with a 1.7-fold increase relative to *D. melanogaster* (*P*_adj_ = 1.26×10^-78^; **Supplemental Table 7**; see also [180-184]). The survey of TE distribution across the genome of *D. yakuba* reveals an accumulation in regions with reduced crossover rates, clustered near the centromeres and, to a lesser degree, telomeres (**Figure 3**). Across the *D. yakuba* genome, TE presence is negatively correlated with crossover rates (Spearman’s ρ = -0.543, *P* = 5.2×10^-41^) (**Supplemental Table 8**). Although TEs in *D. melanogaster* also show a significantly negative association with crossover rates (ρ = -0.354, *P* = 1.6×10^-15^), the correlation is significantly more negative in *D. yakuba* than in *D. melanogaster* (Fisher’s Z transformation, *P* < 8×10^-15^), a difference that is not expected if epigenetic factors were the only cause of the negative association. A similar pattern is observed when only INE-1 elements are analyzed (**Supplemental Table 8**). More generally, the study of TE classes (**Supplemental Table 9**) or each TE independently indicates that the distribution of SINE elements and Class I retrotransposons, which includes INE-1, show stronger correlations between abundance and crossover rates in *D. yakuba* than in *D. melanogaster*. Combined, these results suggest that *D. yakuba* shows a distribution of TEs and crossover rates across the genome that fits better population genetic expectations rather than direct epigenetic effects of TE presence on crossover distribution, and is a first indication that *D. yakuba* may have a more stable landscape of crossover rates than *D. melanogaster*.

To gain insight into the temporal dynamics of TE accumulation, we focused on the large regions showing a clear difference in crossover rates between *D. yakuba* and *D. melanogaster*. Regions with centromere effect in *D. yakuba* but not in *D. melanogaster* show significantly more TEs in *D. yakuba* than in *D. melanogaster* (Mann–Whitney U test, *z* = 6.04, *P* < 1 × 10^-6^). On the other hand, regions affected by the centromere effect in both *D. yakuba* and *D. melanogaster* show similar TE presence in both species (*z* = - 0.01297, *n.s.*). These results are in agreement with predictions of models of selection and linkage if the stronger centromere effect has been fairly stable in *D. yakuba*.

### Short DNA Motifs and crossover localization

We investigated whether short DNA motifs associated with intragenomic variation in crossover rates in *D. melanogaster* [185] and *D. simulans* [186] are also significantly enriched near crossovers in *D. yakuba* (see Supplemental Materials and Methods). These motifs contain [A]_n_ , [CA]_n_, [TA]_n_, [GCA]_n_ and a more variable [CYCYYY] _n_ (**Supplemental Figure 3**). All nine individual motifs identified in *D. melanogaster* are significantly enriched near crossovers also in *D. yakuba* when studying crossovers flanked by diagnostic SNPs separated by a maximum of 3 kb (or 5kb). All except a variable form of [CA]_n_ are also significantly enriched near crossovers with a resolution of 1-kb (see **Table 3** and **Supplemental Table 10**). Because several of the motifs have overlapping characteristics within the [A]_n_ and [CA]_n_ classes, we combined them and observed that both [A]_n_ and [CA]_n_ classes of motifs are very significantly enriched near crossovers in *D. yakuba* at all resolutions analyzed (**Table 3**).

**Table 3.**
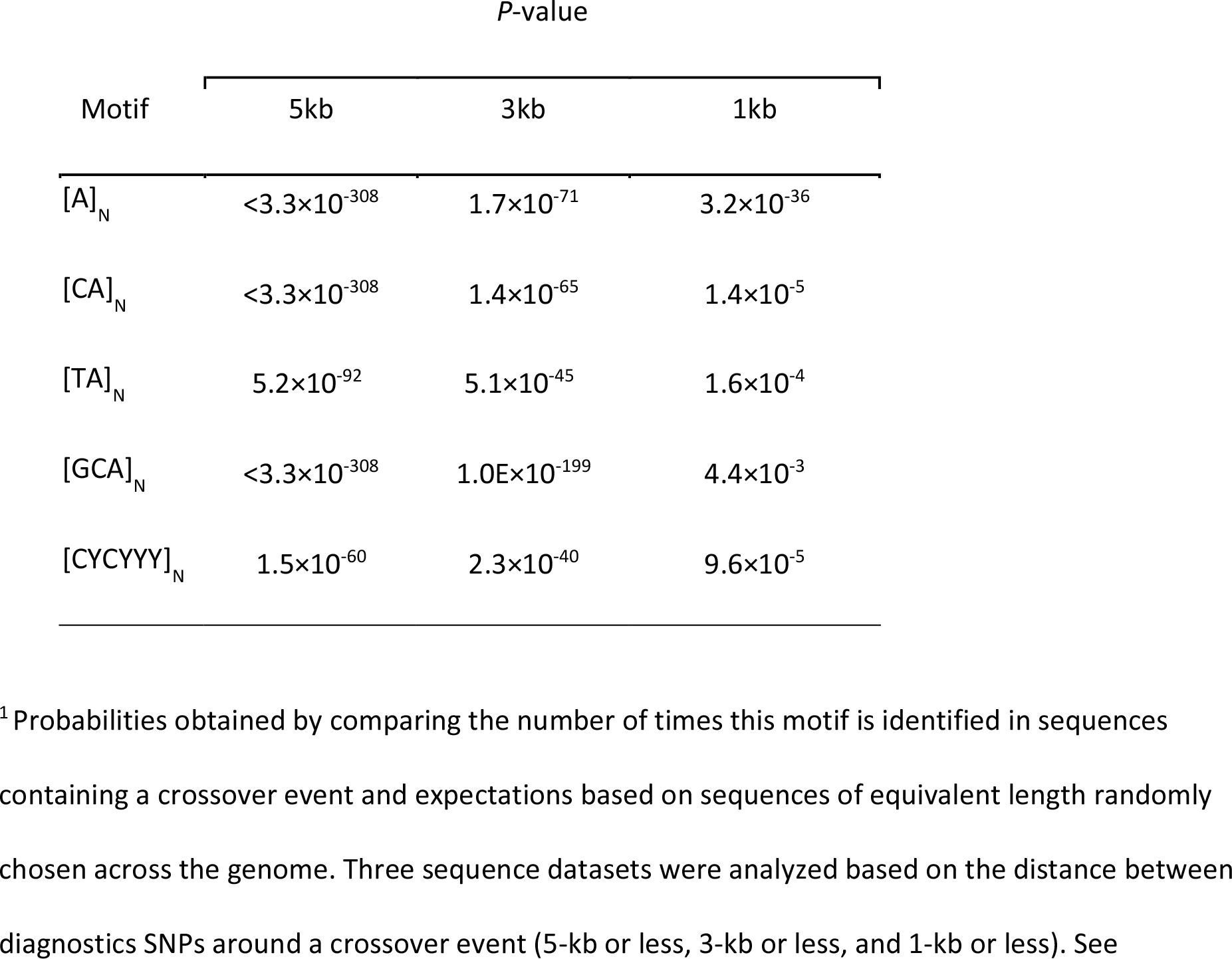
Enrichment analysis of motifs near crossover events in *D. yakuba*^1^.

| Motif | P-value |  |  |
| --- | --- | --- | --- |
|  | 5kb | 3kb | 1kb |
| [A]<sub>N</sub> | $<3.3 \times 10^{-308}$ | $1.7 \times 10^{-71}$ | $3.2 \times 10^{-36}$ |
| [CA]<sub>N</sub> | $<3.3 \times 10^{-308}$ | $1.4 \times 10^{-65}$ | $1.4 \times 10^{-5}$ |
| [TA]<sub>N</sub> | $5.2 \times 10^{-92}$ | $5.1 \times 10^{-45}$ | $1.6 \times 10^{-4}$ |
| [GCA]<sub>N</sub> | $<3.3 \times 10^{-308}$ | $1.0 \times 10^{-199}$ | $4.4 \times 10^{-3}$ |
| [CYCYYY]<sub>N</sub> | $1.5 \times 10^{-60}$ | $2.3 \times 10^{-40}$ | $9.6 \times 10^{-5}$ |

<sup>1</sup> Probabilities obtained by comparing the number of times this motif is identified in sequences containing a crossover event and expectations based on sequences of equivalent length randomly chosen across the genome. Three sequence datasets were analyzed based on the distance between diagnostics SNPs around a crossover event (5-kb or less, 3-kb or less, and 1-kb or less). See

### Evolutionary consequences and dynamics of intragenomic variation in crossover rates

#### Crossover rates and the efficacy of selection on synonymous codons

In good agreement with predictions of population models of linkage and selection, estimates of the strength of CUB (-ENC) in *D. yakuba* genes show a very significant and positive correlation with our *D. yakuba* high-resolution crossover rates (*Rec_yak_*): *R* = +0.228 (*P* = 6.0×10^-13^) for *X*-linked genes and *R* = +0.133 (*P* = 1.3×10^-22^) for autosomal genes. We also re-analyzed CUB in *D. melanogaster* for the same set of genes now with high-resolution crossover rates (*Rec_mel_*) and confirmed the previous results: CUB increases with crossover rates along the *X* chromosome (*R* = +0.170, *P* = 8.9×10^-8^) whereas autosomal genes show a significant and negative correlation (*R* = -0.075, *P* = 3.8×10^-8^). These results suggest that previous incongruent results in *D. melanogaster* were not caused by the use of inadequate selection models (given that these models fit remarkably well *D. yakuba* data) but rather by departures from model assumptions when using *D. melanogaster* data. More specifically, the fit of *D. yakuba* data to models that assume constancy in linkage effects suggests that the landscape of crossover rates might have been fairly stable along the *D. yakuba* lineage whereas a change in the recent past of *D. melanogaster* may have likely occurred, as previously proposed by [98, 131].

To explore this possibility, we first investigated how well *Rec_yak_* would predict CUB in *D. melanogaster* and how well *Rec_mel_* would predict CUB in *D. yakuba*. We show that crossover rates in *D. yakuba* are better predictors of intragenomic variation in CUB across the *D. melanogaster* genome than contemporary crossover rates from *D. melanogaster*, and that the results fit models of selection and linkage along both the *X* chromosome and autosomes (**Figure 7**). *D. melanogaster* CUB in *X*-linked genes shows a stronger positive association with crossover rates in *D. yakuba* (*Rec_yak_*; *R* = +0.201, *P* = 2.2×10^-10^) than with crossover rates in *D. melanogaster* (*Rec_mel_*; *R* = +0.170; *P* = 8.9×10^-8^). CUB in autosomal genes in *D. melanogaster* also shows a significantly positive correlation with *Rec_yak_* (*R* = +0.063, *P* = 4.1×10^-6^), even though the use of *Rec_mel_* instead generated a negative association (see above). At the same time, *Rec_mel_* is a poor predictor of CUB along the *X* chromosome of *D. yakuba* (*R* = +0.115, *P* = 0.0003) and shows a negative association with CUB in *D. yakuba* autosomes (*R* = - 0.076, *P* = 2.7×10^-8^), again contrary to predictions. These results, therefore, open the possibility that contemporary crossover maps in *D. yakuba* may recapitulate to some extent the ancestral recombination environment of the *D. melanogaster* subgroup.

**Figure 7.**
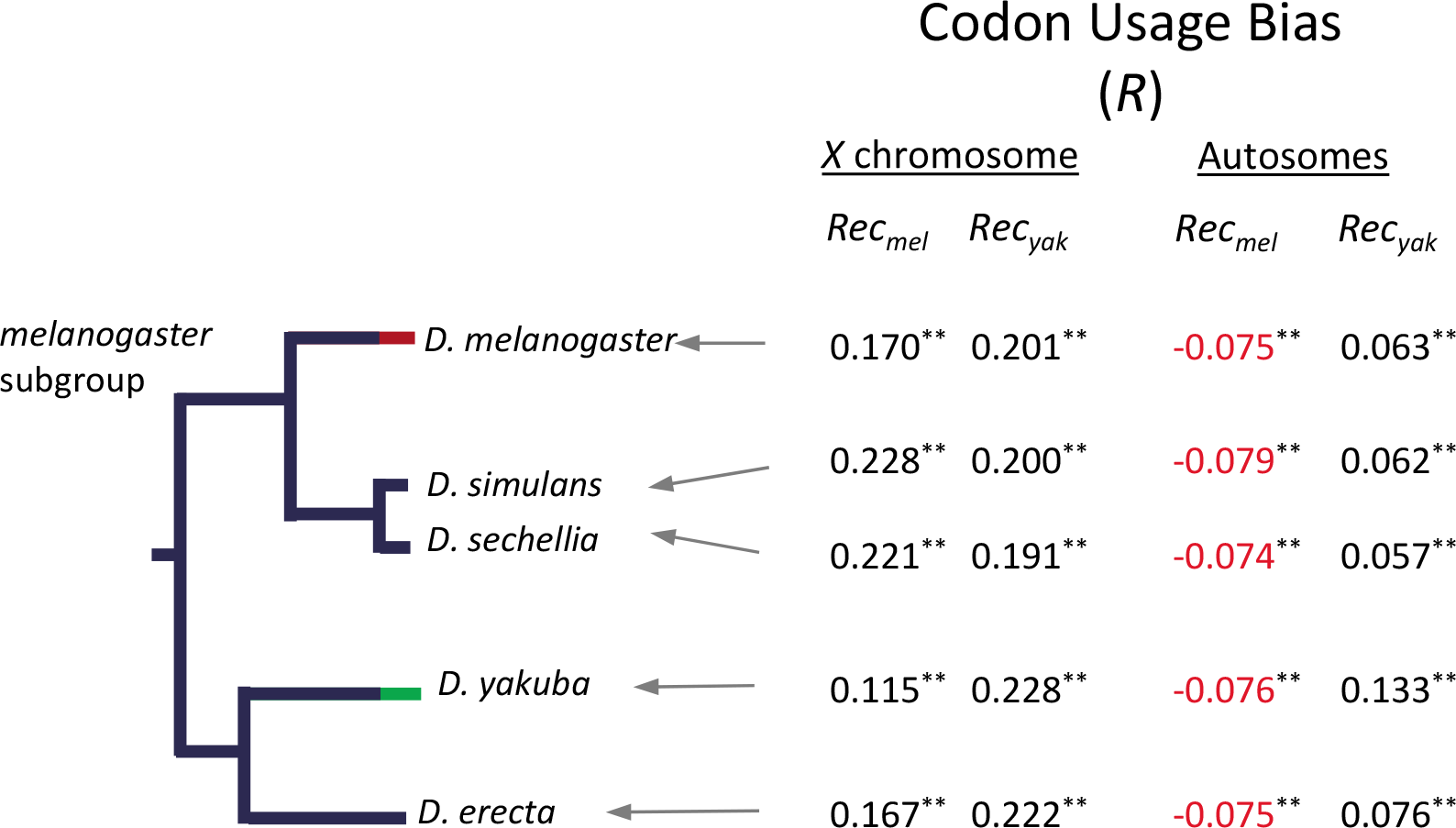
Correlation between CUB and crossover rates in *D. yakuba* (*Rec_yak_*) or *D. melanogaster* (*Rec_mel_*). A generalized linear model (GLM) was used to estimate correlation coefficient *R* between CUB for each gene and log_10_ of crossover rates, either *Rec_yak_* or *Rec_mel_*, for these same genes. Numbers in black indicate significant estimates of *R* in the direction predicted by models of selection and linkage. Red numbers indicate a significant association in the opposite direction than that predicted by models. *, P < 0.05; **, *P* < 0.01); n.s., non-significant associations (*P* > 0.05).

To delve into the prospect that *Rec_yak_* may capture ancestral crossover patterns of the whole *D. melanogaster* subgroup, we obtained regression estimates between *Rec_yak_* and CUB for three other species of the subgroup (*D. simulans*, *D. sechellia* and *D. erecta;* see **Figure 7**). For the *X* chromosome, we observe that *Rec_yak_* predicts almost equally well CUB in *D. yakuba* as CUB in *D. erecta* (*R* = +0.222, *P* = 2.5×10^-12^), a species that separated from *D. yakuba* ∼10.4 Mya [187], and only slightly less well (*R* ranging between 0.191 and 0.201; *P* < 3.0×10^-10^) for *D. simulans* and *D. sechellia,* which separated from *D. yakuba* ∼12.7 Mya [187]. Combined, these results strengthen the notion that *Rec_yak_* for the *X* chromosome captures to a significant degree the ancestral crossover rates of the whole subgroup and that *Rec_yak_* has been remarkably constant for more than 40 My of evolution. For autosomes, our results suggest a faster evolution of crossover landscapes relative to the *X* chromosome within the *D. melanogaster* subgroup. Nonetheless, *Rec_yak_* keeps being a good predictor of CUB across the entire phylogeny analyzed while *Rec_mel_* shows a negative association with CUB for all species, indicating that significant changes in crossover landscapes in the *D. melanogaster* lineage had to occur not only after the split from its common ancestor with *D. simulans/D. sechellia* 5.4 Mya [187] but also very recently in order to severely limit the linkage effects of the new crossover landscape on CUB.

Note that the conclusions about temporal stability of crossover rates based on the analysis of CUB would be also valid if the cause of biased codon frequency was GC-biased gene conversion (gcGBC) insted of, or in combination with, translational selection [188, 189]. This is because the resolution of heteroduplex DNA that arises during recombination is GC-biased and preferred synonymous codons are often G- and C-ending. gcBGC, therefore, directly predicts a positive relationship between crossover rates across genomes and the degree of GC-ending codons (and hence CUB) that mimics the predictions of weak selection [190].

#### Crossover rates and the efficacy of selection on protein evolution

To identify variation in the efficacy of selection at protein level, we estimated rates of nonsynonymous (*d*_N_) and synonymous (*d*_S_) evolution, and the *d*_N_/*d*_S_ ratio (*ω*) across the *D. melanogaster* subgroup using the branch-model in the *codeml* program as implemented in PAML [191, 192]. Moreover, to capture fine-scale intragenomic effects of variation in crossover rates, we analyzed rates of evolution using single-gene data, without grouping genes into crossover-groups. Additionally, we took into account gene-specific properties possibly influencing *ω* and used residuals as a measure of variable efficacy of selection on amino acid changes (*ω*_R_) for each gene and phylogenetic branch (see Materials and Methods).

We first investigated changes in efficacy of selection on weakly deleterious amino acid mutations by focusing on genes with no signal of rapid evolution (see Materials and Methods and [104]). Variation in contemporary crossover rate across the *D. yakuba* genome is negatively associated with gene-specific rates of protein evolution along the *D. yakuba* (*ω*_R-yak_) lineage genome-wide (*r* = -0.047, *P* = 0.0002), as well as for the *X* chromosome (*r* = -0.078, *P* = 0.02) and autosomes (*r* = -0.043, *P* = 0.002) (**Figure 8A**). Along the *D. melanogaster* lineage, *ω*_R-mel_ shows a weak negative association with *Rec_mel_* genome-wide (*r* = -0.026, *P* = 0.044) that is also observed for the *X* chromosome (*r* = -0.100, *P* = 0.002) but not for autosomes (*r* = -0.012, *P* > 0.05). Equivalent to our results with CUB, we observe that *Rec_yak_* is a better predictor of rates of evolution along the *D. melanogaster* lineage than *Recmel*; *Rec_yak_* shows a significantly negative association with *ω*_R-mel_ genome-wide (*r* = -0.053, *P* = 0.00004). In fact, contemporary crossover rates in *D. yakuba* show a weak but significantly negative association with rates of protein evolution across the whole phylogeny (*ω*_R-total_) (*r* = -0.036, *P* = 0.0044) while *Rec_mel_* shows no significantly association with *ω*_R-total_ (**Figure 8A**).

**Figure 8.**
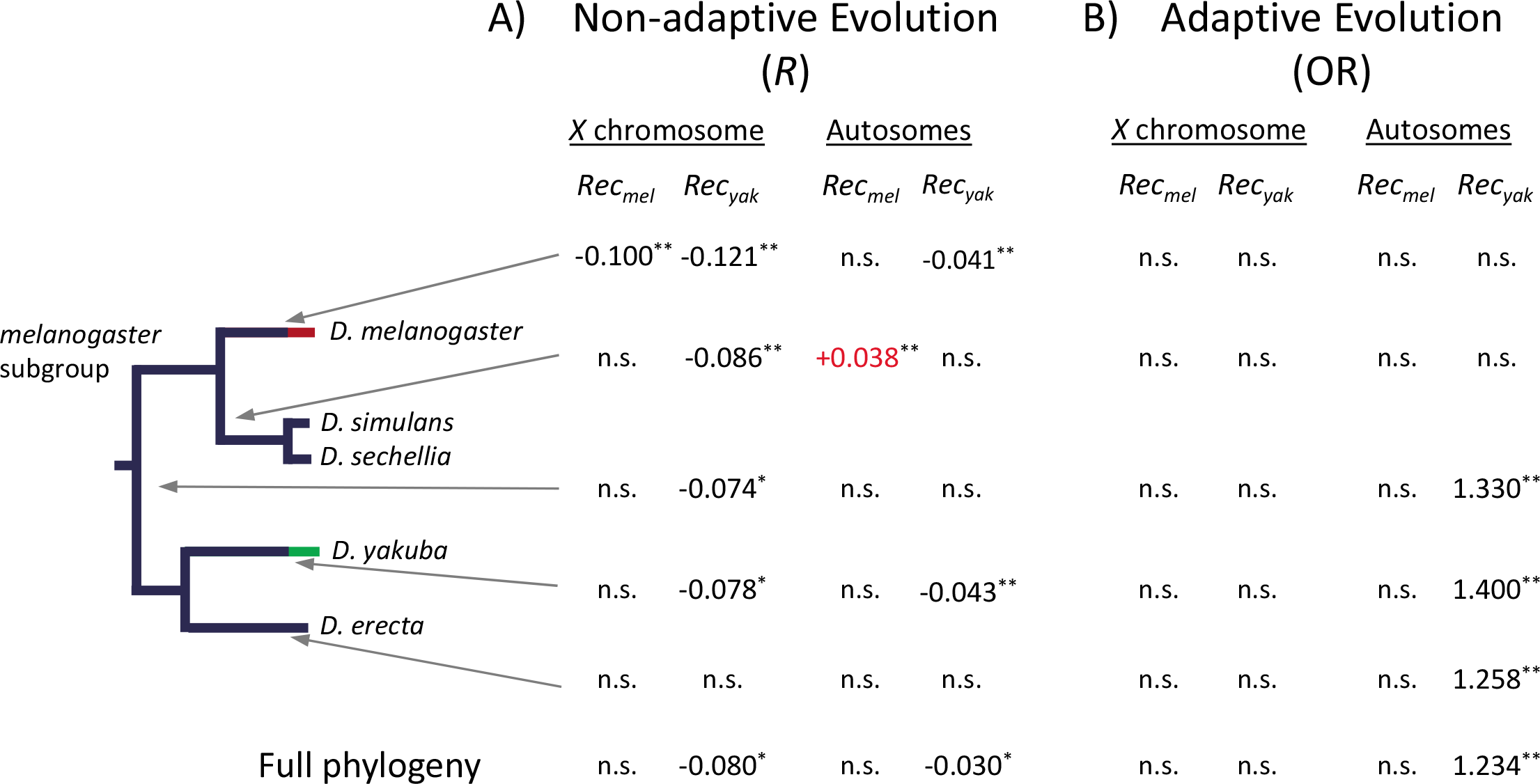
Correlation between rates of protein evolution and crossover rates in *D. yakuba* (*Rec_yak_*) or *D. melanogaster*(*Rec_mel_*). **A)** For each branch across the *D. melanogaster* subgroup, estimates of the efficacy of selection on amino acid changes (*ω*_R_) were compared to crossover rates, either *Rec_yak_* or *Rec_mel_*, for these same genes with a generalized regression model (GRM) to estimate correlation coefficient *R.* **B)** For each branch across the *D. melanogaster* subgroup, genes with and without signal of positive selection based on PAML (see text for details) were compared to crossover rates for these same genes. Odds Ratio (OR) from logistic regression analysis was applied to capture variable likelihood of positive selection with crossover rates (see text for details). Numbers in black indicate significant estimates of *R* in the direction predicted by models of selection and linkage. Red numbers indicate a significant *R* in the opposite direction than that predicted by models. *, P < 0.05; **, *P* < 0.01); n.s., non-significant associations (*P* > 0.05).

When studying the relationship between *Rec_yak_* and *ω*_R_ for different branches of the phylogeny, we confirm a consistently significant and negative association for the *X* chromosome for most branches (except for the *D. erecta* lineage), and a less conserved trend for autosomes (**Figure 8A**). *Rec_mel_*, on the other hand, shows no signal of being negatively associated with *ω*_R_ along any lineage outside *D. melanogaster*; it even shows a positive association (contrary to predictions) with rate of protein evolution along the *D. simulans* lineage. In agreement with the results based on CUB, these analyses support a very recent change in crossover landscape in *D. melanogaster* and, additionally, identify a likely change in crossover patterns in *D. simulans*/*D. sechellia* after the split from their common ancestor with *D. melanogaster*. Therefore, our studies of protein evolution also support the conclusions that contemporary crossover rates across the *D. yakuba* genome capture ancestral properties of crossover landscapes, with the *X* chromosome showing a stronger degree of conservation.

We also investigated whether variation in crossover rates is associated with the probability of adaptive protein evolution (based on PAML analyses; see Materials and Methods). To this end, we applied a logistic regression test (or logit) that allows the study of binary variables (a gene showing or not signals of positive selection along a lineage) as a response to a continuous variable (crossover rates), without grouping genes into arbitrary crossover classes. We observe that contemporary *Rec_yak_* is a good predictor of adaptive protein evolution along the *D. yakuba* lineage [Odds Ratio (OR) = 1.33, *P* = 0.0017]. Outside the *D. yakuba* lineage, *Rec_yak_* shows a similar association with adaptive protein evolution along the *D. erecta* lineage and along the lineages ancestral to both *D. yakuba*/*D. erecta* and *D. melanogaster*/*D. simulans*/*D. sechellia* (**Figure 8B**), again suggesting that *Rec_yak_* captures some of the ancestral crossover landscape of the *D. melanogaster* subgroup. Contemporary *Rec_mel_* shows no significant association with the likelihood of adaptive amino acid changes along any of the lineages analyzed (including the *D. melanogaster* lineage), in agreement with previous studies [104] and with the proposal of a recent change in the distribution of crossover rates across the genome of *D. melanogaster* [98, 131].

#### Crossover rates and variation in levels of neutral diversity

We estimated neutral diversity at four-fold synonymous sites (π_4f_) across the *D. yakuba* and *D. melanogaster* genomes as a proxy for recent *N_e_* (see Materials and Methods) and compared these estimates with our high-resolution crossover landscapes for non-overlapping 100 kb regions. In *D. yakuba,* we observe that variation in crossover rates is a very strong predictor of π_4f_ for both autosomes [*r* = 0.74, *P* = 7.6×10^-169^] and the *X* chromosome (*r* = 0.63, *P* = 5.6×10^-25^), as expected based on models (**Figure 9A**). In *D. melanogaster,* we estimate equivalent trends when using similar methodologies and crossover map resolutions: *r* = 0.64 (*P* = 5.4×10^-110^) and *r* = 0.568 (*P* = 3.3×10^-20^) for autosomes and the *X* chromosomes, respectively (**Figure 9B**). These results show that contemporary *Rec_mel_* does a good job capturing heterogeneity in recent *N_e_* across the *D. melanogaster* genome, and emphasizes the disparity between long- and short-tem *N_e_* along the *D. melanogaster* lineage when combined with the results from CUB and protein evolution. At the same time, we observe that the *D. yakuba* lineage may conform better than *D. melanogaster* to models of selection and linkage when applying methodologies that assume consistency between long-term and short-term *N_e_*.

**Figure 9.**
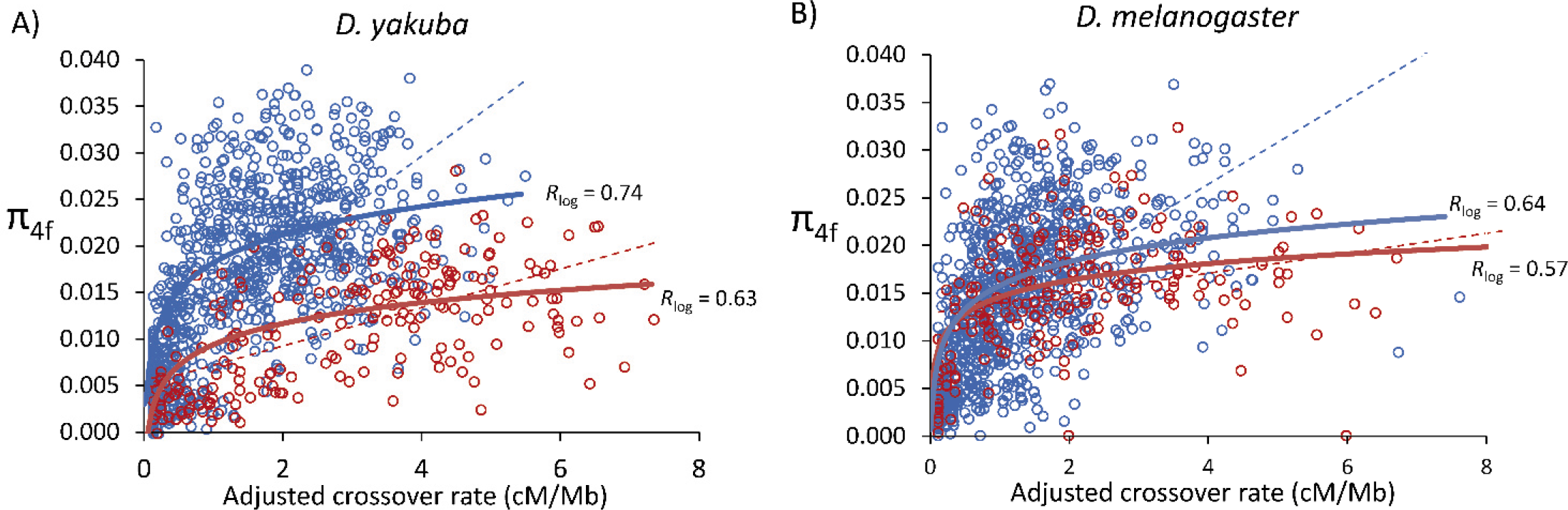
Relationship between crossover rate (cM/Mb) and levels of neutral nucleotide polymorphism (π_4f_) in *D. yakuba* and *D. melanogaster*. π_4f_ indicates pairwise nucleotide variation (/bp) at four-fold synonymous sites. Autosomal and X-linked regions are indicated as blue and red circles, respectively. Crossover rates are adjusted to allow a direct comparison between X-linked and autosomal regions. Dashed lines indicate linear regressions between π_4f_ and adjusted crossover rates; solid lines indicate regressions between π_4f_ and the log_10_ of crossover rates. Data shown for non-overlapping 100-kb windows.

## DISCUSSION

Here we report a genome-wide crossover map for *D. yakuba* using WGS and a new dual-barcoding methodology. Our method reduces library and sequencing costs significantly, allowing for the genotyping of a large number of individuals and, ultimately, the generation of genetic maps at very high resolution. The dual-barcoding approach involves crosses between multiple wild-type parental genotypes and, in turn, provides the opportunity to capture species-wide genetic maps that are more informative for genomic and evolutionary analyses than those generated using a single cross. Our crossing and genotyping schemes also reduce the potential effects of using inbred lines when studying meiotic outcomes. Nevertheless, the study of the frequency of genotypes in the chromatids analyzed is an important—if not required—step to asses directly the potential role of egg-to-adult viability deffects. In our case, the study of the frequency of parental and recombining genotypes allowed us to rule out detectable viability effects biasing the meiotic products genotyped.

We analyzed a large number of individual meiotic events (more than 1,600) and identified the precise localization of more than 5,300 crossover events. Amongst *Drosophila* species, *D. yakuba* shows an intermediate total map length (339 cM), longer than *D. melanogaster* (287 cM [41, 113, 136]) and shorter than *D. mauritiana* (about 500cM; [31]), *D. pseuddobscura* (>450 cM; [193, 194]) or *D. virilis* (732 cm [117]). This high degree of variation is consistent with the proposal that crossover rates and distribution have the potential to evolve very fast and, in particular, with recent studies suggesting that these changes can be the result of adaptive processes ([118, 126] see also [127, 129, 195]). Although our study in *D. yakuba* provides confirmatory evidence to some information garnered from the model species *D. melanogaster,* most of our analyses in *D. yakuba* reveal clear differences and a contrasting view of crossover distribution, control and evolution in *Drosophila*. As such, this study adds to the expanding notion that a deep understating of the causes and consequences of crossover rate variation across genomes benefits from the analysis of multiple species within an appropriate phylogenetic context.

Before this study, a general trend among *Drosophila* species appeared to support a link between the length of genetic maps and the magnitude of the centromere effect; *D. simulans*, *D. mauritiana*, *D. pseudoobscura* and *D. persimilis* all show higher number of crossovers per meiosis and weaker centromere effect than *D. melanogaster* [31-34, 126]. Such a trend suggested that the centromere effect is either a direct cause or a consequence of lower genome-wide crossover rates, or that the two processes coevolve. *D. yakuba* shows, however, a significantly higher crossover rate and overall longer genetic map than *D. melanogaster* while also showing a greater centromere effect, thus decoupling the two observations for the first time in *Drosophila*. Our results in *D. yakuba*, the species with the greatest centromere effect in the *Drosophila* genus to date, uncover a more complex relationship between crossover distribution and the centromere effect.

A strong centromere effect also limits substantially the genomic region naturally available for multiple crossover events along a chromosome arm, begging the question of whether this impacts crossover interference*. D. yakuba* shows significant crossover interference, as observed in all other *Drosophila* species analyzed. Notably, the chromosome arms with stronger centromere effect (*2L*, *3L* and *3R*) also show a higher degree of crossover interference than chromosome arms with weaker centromere effects (*2R* and *X*) (see **Table 2** and **Supplemental Table 3**). Combined, these results would support the concept of interconnected molecular mechanisms controlling the overall number of crossovers, centromere effects and crossover interference with outcomes that may vary among chromosome arms.

Our study in *D. yakuba* also differs from multiple reports in other *Drosophila* species showing that the suppression of crossovers in inversion heterozygotes increases the number of crossovers on freely recombining chromosome arms (the interchromosomal effect [39, 134, 135]). The presence of a heterozygous inversion on chromosome arm *2R* in *D. yakuba* plays no detectable role in increasing the number of crossovers in any of the freely recombining chromosome arms. A key difference between *D. yakuba* and most *Drosophila* species where the interchromosomal effect has been studied is the higher number of crossovers per female meiosis in the former species. Therefore, although it is not possible to fully disentangle the effects of the genetic backgrounds carrying inversions, the results of our *D. yakuba* study open the possibility that the higher number of crossovers per chromosome arm in this species limits the potential increase in crossovers and the overall magnitude of the interchromosomal effect.

### Crossover assurance and crossover-associated meiotic drive in *D. yakuba*

*D. yakuba* shows a large fraction of recovered chromatids with multiple crossovers, particularly for the *X* chromosome. As a consequence, and contrary to studies in *D. melanogaster*, tetrad analysis in *D. yakuba* requires the inclusion of E_≥3_ tetrads to capture the full observed distribution of meiotic products. Our tetrad analysis shows that *D. yakuba* has a stronger degree of crossover assurance than *D. melanogaster*, compatible with absolute crossover assurance under benign conditions for wild-type genotypes. We also show results that strongly support the presence of an active crossover-associated meiotic drive mechanism (MD_CO_) in *D. yakuba* that acts in addition to strong crossover assurance for the *X* chromosome and results in the preferential inclusion in oocytes of chromatids with crossovers (**Figure 6A**).

More specifically, we show that models without bias in transmission or with biases in distribution of crossovers between sister chromatids are incompatible with observed data for the *X* chromosome. A model allowing MD_CO_ not only fits the data far better than any of the other models investigated but also fits the data well. Moreover, our WGS approach together with the observation that this phenomenon is identified in all three crosses of wild-type strains make unlikely that the results are caused by viability effects. We, therefore, propose that *D. yakuba* has evolved increased effective crossover rates on the *X*, at least partly, by meiotic drive rather than by increasing the number of tetrads with two or more crossovers. In all, our analyses predict a total of 5.98 ─ 6.33 crossover events per female meiosis that would result in 3.46 detectable crossovers per viable meiotic product. The same quantitative study suggests that this MD_CO_ mechanism would be equivalent to a 15 to 34% increase in *X* chromosome homologous recombination events in offspring relative to a case with no drive but the same number of meiotic crossover events in prophase of meiosis I.

Our proposal of an active MD_CO_ mechanism is not completely new. Meiotic drive for recombinant chromatids at meiosis II has been proposed to be active in the human female germline based on one study that genotyped oocytes and polar bodies of women with average age of 37.3 years [196]. In *D. melanogaster*, Singh *et al*. (2015) [197] showed an increase in recombinant offspring (rather than a direct increase in crossovers in Meiosis I) as a plastic response to parasite pressures, suggesting an epigenetic activation of transmission distortion and assymetries during meiosis II, equivalent to the MD_CO_ mechanism. Our results suggest that an equivalent mechanism may be active under benign conditions for the *X* chromosome in *D. yakuba*. This said, and given the standard observation that *E*_0_ is smaller on the *X* chromosome than on autosomes also in *D. melanogaster*, it is possible that MD_CO_ is active under benign conditions in the *D. melanogaster X* chromosome as well, but it has not yet been identified due to the smaller number of crossovers. In fact, we show that MD_CO_ is compatible with *D. melanogaster X* chromosome data (**Figure 6D**). In this regard, the detailed genetic and cellular study of other species of the *D. melanogaster* subgroup can provide additional support to the possibility of a more widespread MD_CO_, at least for the *X* chromosome, that may be enhanced under stress conditions.

This leaves open the question of what might be the selective advantage, if any, to an active MD_CO_ mechanism on the *X* chromosome. Modifers of crossover rates are known to segregate in natural populations of many species [118, 120, 128, 198-201] and models of selection and recombination predict an overall benefit to increasing rates when finite populations are considered (see [202] and references therein). Increased crossover rates could be particularly frequent on the *X* chromosome because this chromosome would be more likely to take advantage of recesive beneficial modifiers that increase rates relative to autosomes, a possibility that would fit with *Drosophila* studies showing stronger positive selection acting on the *X* at both protein and gene expression levels [103, 203-211]. At the same time, there is a direct limitation to increasing the number of crossovers in meiosis I given the known increased probability of missegregation and higher rates of ectopic exchange and chromosomal aberrations in multi-chiasma tetrads [212, 213]. The proposed meiotic drive in meiosis II would allow taking advantage of the evolutionary benefits of higher recombination rates while limiting the number of crossovers per tetrad. The advantages of such mechanism, moreover, are predicted to be particularly relevant in species with high likelihood of ectopic recombination due to the abundance of TEs, as in the case of *D. yakuba*.

### Transposable elements, satellite repeats and crossover distribution in *D. yakuba*

The study of TE abundance across the genome of *D. yakuba* confirms trends observed in other *Drosophila* species including *D. melanogaster,* with TEs being more frequently observed in genomic regions with significantly reduced crossover rates. Notably, *D. yakuba* shows a stronger negative correlation between TE abundance and crossover rate across the genome than *D. melanogaster*. These results suggest that the proposed epigenetic effects of TEs reducing local crossover rates play a minor role in the observed crossover landscapes and, in turn, are more consistent with models of selection and linkage. Because these models assume long-term equilibrium, our results also suggest a more stable landscape of crossover rates in *D. yakuba* than in *D. melanogaster*.

The study of satellite DNA in *D. yakuba*, combined with previous results from *D. simulans,* confirms that *D. melanogaster* shows a highly evolved composition relative to other related species and provides outgroup information strengthening the idea that the changes in satellite composition in *D. melanogaster* arose after split from *D. simulans* [139, 214]. Our analysis of satellite DNA also suggests, albeit indirectly, shorter centromeres in *D. yakuba* than in *D. melanogaster*. Given the reports linking shorter centromeres to stronger centromere effects in *Drosophila* [138], this result is also congruent with the observed greater centromere effect in *D. yakuba* than in *D. melanogaster*.

### Short DNA motifs and crossover distribution

Short DNA repeat motifs enriched near crossover events in *D. melanogaster* [185] are also overrepresented in genomic regions with high crossover rates in the closely related species *D. simulans* [186]. Here, we showed that these same motifs are also enriched near crossover events in *D. yakuba* despite the greater evolutionary distance. Notably, many of these motifs share properties associated with high degree of open chromatin and DNA accessibility [185], and there is the additional observation that in *Drosophila*, as in other organisms, crossovers are enriched at or near transcriptionally active regions [215, 216]. In this regard, it is important to consider DNA:RNA hybrids (R-loops) during transcription and the proposed induction of DNA damage, instability and, ultimately, double-strand breaks [217-220]. Interestingly, poly A/T tracts, which is one of the crossover-associated motifs in *Drosophila* and in yeast [119, 221, 222], play a causal role in R-loop formation [223]. In all, therefore, crossover rate variation at the scale of several megabases seems best explained by centromere and telomere effects whereas variation at a finer or local scale would be associated with open chromatin, active transcription and specific DNA motifs. The inclusion of R-loops into the paradigm to describe crossover localization across *Drosophila* genomes is particularly appealing because it would provide a testable mechanistic explanation for the known influence on crossover rates and distribution of epigenetic changes due to environmental conditions and stress.

### Evolutionary consequences and dynamics of intragenomic variation in crossover rates

Because crossover rates vary across genomes, models of selection and linkage predict that *N_e_* will vary across genomes as well, as a function of crossover rates and gene distribution. Therefore, multi-locus analyses of diversity and selection are difficult and mostly unadvisable. To deal with this intragenomic variation in *N_e_*, the standard approach is to combine inter- (divergence) and intra-specific (diversity) data and assume that contemporary *N_e_* for a given locus is representative of past, long-term *N_e_* influencing efficacy of selection and rates of evolution [73, 80, 83, 84, 86, 88, 95, 106, 107, 224-231]. However, genomic landscapes of crossover rate vary between species evidencing that *N_e_* at a given locus changes with time (additional to demographic events) and, therefore, contemporary *N_e_* may differ from long- term *N_e_* in many species and genes. The reasons for not fully embracing temporal changes in crossover rates and *N_e_* are mostly practical ones. Population genomic data can be easily obtained today, providing detailed views of nucleotide variation within and between species. In contrast, genome-wide high-resolution crossover maps are much less common because they need both very high marker density (ideally WGS or equivalent) and many individuals genotyped; high marker density or high number of genotyped individuals alone is not sufficient to generate accurate high-resolution maps. In all, the assumption of temporally stable genomic landscapes of *N_e_* can produce inaccurate parameterization of linkage effects and selection, and thus hinders our understanding of the causes of variation within and between species.

Among *Drosophila* species, detailed, experimentally generated, crossover maps exist for *D. melanogaster* [41, 113] and *D. pseudoobscura* [32, 118] but equivalent data is still infrequent for closely related species. Within the *D. melanogaster* subgroup, sparse crossover rate maps for the sister species *D. simulans* and *D. mauritiana* showed higher crossover rates and a different crossover landscape than *D. melanogaster* [31]. Limited data for the *X* chromosome of *D. yakuba* also suggested higher crossover rates than *D. melanogaster* [232, 233]. Recent studies of meiosis genes comparing *D. melanogaster* and *D. mauritiana* further suggest that the change in crossover rates likely occurred in the *D. melanogaster* lineage [126], but there was no outgroup data to strengthen that conclusion. More generally, the lack of multispecies high-resolution crossover landscapes in the *D. melanogaster* subgroup has limited attempts to quantify the effects of linkage on the efficacy of selection in this species complex. Here, we used the new high-resolution contemporary crossover rate map for *D. yakuba* to add phylogenetic context for crossover landscapes within the *D. melanogaster* subgroup, and investigate the evolutionary influence of variation in crossover rates across the genome and time.

Our studies show that the weak support for an effect of crossover rate variation on the efficacy of selection when using *D. melanogaster* as focal species is, to a large degree, due to a widespread change in crossover rates in the very recent history of *D. melanogaster,* after the split from its common ancestor with *D. simulans.* Indeed, contemporary crossover rates in *D. melanogaster* (Rec_mel_) are a good predictor of diversity and contemporary *N_e_* within this species, but they are very poorly associated with estimates of long-term efficacy of selection (CUB, efficacy of selection against deleterious amino acid changes and the incidence of adaptive amino acid changes). As a consequence, analyses in *D. melanogaster* that assume temporally stable crossover landscapes will be inaccurate.

In contrast, the *D. yakuba* lineage seems to have had a more stable crossover landscape that is well captured by our new contemporary crossover maps. Studies of neutral diversity across the genome of *D. yakuba* fit well patterns predicted by models of selection and linkage. That is, contemporary Rec_yak_ captures well recent intragenomic contemporary *N_e_* as well as long-term *N_e_*. The conclusion of high stability in crossover rates along the *D. yakuba* lineage is confirmed when considering the whole *D. melanogaster* subgroup, with the contemporary crossover landscape of *D. yakuba* capturing information of ancestral linkage effects across the whole *D. melanogaster* subgroup, even for the *D. melanogaster* lineage. These results suggesting a fairly stable crossover landscape in *D. yakuba* are also congruent with the study of TE distribution, which indicates long-term co-evolutionary dynamics of TE abundance and crossover rates. Combined, these results strongly support the idea that *D. yakuba* may be an adequate species to study population genetics predictions influenced by crossover rates.

Our study represents only one step towards a better understanding of temporal variation in crossover rates and their consequences in *Drosophila*. More generally, our analyses underscore the importance of generating high-resolution crossover rate maps from closely-related species, as a necessary counterpart for every species with population genomic data and within a coherent phylogenetic context. Minimally, this information will allow testing assumptions of temporal stability of crossover landscapes. Moving forward, the additional information that these crossover maps provide will refine our understanding of crossover control during meiosis and activate research towards theoretical and modelling frameworks to study and parameterize evolutionary parameters amid changes in crossover rates.

## MATERIALS AND METHODS

### Dual-barcoding genotyping method

We used a genotyping approach with two layers of barcoding to reduce sequencing and labor costs (see **Figure 1**). In typical Illumina sequencing, each sample receives one adapter (one library sequence barcode), which allows assigning reads to different samples when sequenced together. In our method, we use multiple parental lines (line 1, line 2, line 3, line 4, etc.), and each line is involved in a different cross (cross 1: line 1 x line 2; cross 2: line 3 x line 4, etc.). F_1_ virgin females are crossed to males of a ‘tester’ line to generate F_2_ individuals and we combine multiple F_2_ individuals, one from each cross, per library (per sequence barcode). After sequencing, reads are separated based on sequence barcode and, for each set of reads sharing a sequence barcode, SNPs specific to a single parental line (singleton SNPs) are used as diagnostic SNPs or genetic barcodes. Reads containing diagnostic SNPs can be then assigned to a specific parental genome and cross without ambiguity, and sequence reads not containing any diagnostic SNP or with diagnostic SNPs for the tester line are discarded.

### Crossing scheme and sequencing of the parental genomes

Each of the seven *D. yakuba* lines used in this study was derived from the offspring of a single female (isofemale lines). Lines *TZ043* and *TZ020* were established from females collected on July 2001 in the Upanga District of Dar es Salaam (Tanzania; kindly provided by Bill Ballard), lines *SN20* and *SN17*, from females collected on January 2004 in the São Nicolau waterfall (São Tomé Island; collected by Ana Llopart), lines *Cost* 1235.2 and *Obat* 1200.5, from females collected on March 2001 in the Obo Natural Reserve (São Tomé Island; collected by Daniel Lachaise), and line *Rain* 5 from a female collected on July 2009 in the Obo Natural Reserve (São Tomé Island; collected by Ana Llopart). The three crosses we used to obtain recombination maps were: *TZ043* × *TZ020, Sn20* × *Sn17* and *Rain 5* × *Cost* 1235.2. One-day old F_1_ female offspring from these crosses were crossed with males from the tester line *Obat* 1200.5. This last cross to the tester line was important to allow the generation of haploid sequences from the parental lines bioinformatically when sequencing F_2_ individuals (see [113]).

We generated an improved *D. yakuba* genome reference sequence of Tai18E2 using PacBio and Illumina sequencing (Supplemental Materials and Methods) that was then used to generate high-quality genome sequences for all the parental lines used in the crosses, including the tester line (see Supplemental Materials and Methods). In short, to generate genome sequences of parental lines (and to correct PacBio sequences) we used two rounds of mapping, each one including Bowtie2 [234] and Stampy [235] mapping. Variants were called using samtools mpileup version 0.1.18 with minimum base quality of 30 and minimum mapping quality of 35 followed by BCFtools view to filter variants with a minimum read depth 3 and a fraction of reads supporting the variant greater than 0.8. After filtering, vcfutlis vcf2fq was used to convert the VCF files to fastq format followed by seqtk fq2fa to convert to fasta format [236]. Our approach generated final sequences for parental lines that only show non-N positions when high quality information is present.

### Diagnostic SNPs

Diagnostic SNPs were defined as sites where all parental (including the tester) genomes contain a high quality nucleotide call and only one genome had a different nucleotide variant (singleton SNP). Sites with low quality or ambiguity in one or more genomes, or with 3 or 4 nucleotide variants were removed from further consideration. Because the *D. yakuba* lines used in the crosses were inbred over several generations, they each contain few heterozygous sites and not used unless including a high quality and unambiguously informative variant.

### Sequencing of F_2_ flies

We combined one F_2_ female from each of the three crosses to prepare Illumina libraries with a given custom-designed barcode, following the same experimental protocols described in ’Illumina library preparation’ (Supplemental Materials and Methods). We multiplexed our libraries (112 libraries/lane) based on custom-designed adapters with ≥8-nucleotides that are at least 4 nucleotide changes away from any other barcode, and sequenced them in an Illumina HiSeq 4000 instrument housed at the IIHG (University of Iowa).

Fastx_barcode_splitter and fastx_trimmer from FASTX-Toolkit version 0.014 (http://hannonlab.cshl.edu/fastx_toolkit/) were used to split sequence reads and to remove barcode sequences, respectively. Sequence reads were also filtered with Trimmomatic and parameters as described in ’Illumina alignment pipeline’ (Supplemental Materials and Methods). A total of six Illumina HiSeq 4000 lanes were used in this project, allowing us to genotype up to 1,701 F_2_ individuals.

### Mapping to diagnostic SNPs

To identify reads mapping to a diagnostic SNP of a specific parental line, we first, conservatively, mapped reads to all other parental lines using Bowtie2 [234] with parameters that ensure perfect alignment for the whole read. Reads that did not align to any of these genomes were then mapped to the sequence of the parental genome of interest with the same restrictive parameters. After mapping, samtools mpileup was used to obtain all nucleotides with mapped reads followed by BCFtools view to generate VCF files [236]. Sites not previously assigned as a diagnostic SNP and sites assigned as as diagnostic SNPs for the tester line were filtered out. This process was repeated for every parental line used in the crosses, meaning that every F_2_ individual we sequenced had only one (if all diagnostic SNPs corresponded to a single parental genome) or two (if one or more crossovers occured) VCF files with mapped diagnostic SNPs per chromosome arm, identifying the original cross. Note that the filtering process removed a higher number of chromatids sequenced for chromosome arm *2R* relative to other chromosome arms (Supplemental Table 1) due to a lower fraction of diagnostic (singeton) SNPs. As a consequence, analyses for *2R* are based on fewer chromatids. For each F_2_ individual, we used the ordered distribution of diagnostic SNPs along chromosomes in the VCF files (after merging when needed using BCFtools merge) for the purpose of downstream analysis and crossover localization.

### Identification of crossovers

We used diagnostic SNPs as a method for dual multiplex sequencing but also to identify crossovers and their genomic location. After aligning reads from each F_2_ individual to parental genomes of a specific cross based on diagnostic SNPs, several filters were applied before determining crossover events. First, any F_2_ chromosome arm that contained less than 100 diagnostic SNPs was filtered out. Second, given that our diagnostic SNPs were not evenly distributed across the chromosomes, we removed regions with a significant bias of mapping to one parental genome over the other. We applied a sliding window approach with regions of 100kb and increments of 25kb, and removed from further consideration regions with a ratio of mapped reads to the parental genomes less than 0.1 or greater than 10. Our next filters aimed at decreasing events that could potentially be gene conversions without crossover. A block was defined as a region of continuous mapping to diagnostic SNPs from only one parental genome.

Blocks shorter than 100 kb were removed from analysis and flanking blocks were combined. Crossovers were identified as consecutive diagnostic SNPs of one parental genome switching to diagnostic SNPs from the other parental genome (boundaries between blocks). Crossovers were then assigned a random position between the two diagnostic SNPs defining the region between blocks of diagnostic SNPs. A final filter was applied to require crossover events to be separated by at least 250kb. Analyses using 500kb as threshold produced the same results, in agreement with the observation that the two closest crossovers we identified in our study were 771kb apart. To study crossovers in the dot chromosome, the filters used were relaxed to capture the paucity of SNPs among lines. In this case, the minimum number of diagnostic markers was set to 10, and the minimum number of diagnostic markers per block flanking potential crossovers was set to 4. Even with these more permissive filters, we did not detect any crossovers along the dot chromosome.

Crossover rates were studied as cM/Mb per female meiosis, for each cross separately and after combining the results from the three crosses. Unless noted, the distribution of crossover rates along chromosome arms in *D. yakuba* was based on non-overlapping 200 kb windows. For *D. melanogaster*, we analyzed equivalent genomic sizes based on high-resolution crossover maps for *r5.3* and r*6* genome releases [113, 237]. In all cases, we used log_10_ of crossover rates in analyses to estimate association between crossover rates and *N_e_* or efficacy of selection (CUB, protein evolution and neutral diversity).

### Analyses of viability effects

Differential egg-to-adult viability of parental and/or recombinant genotypes could bias the set of chromatids genotyped in adults. We, therefore, quantified the frequency of the four possible pairwise combinations of the parental genotypes for all possible genomic distances along the *X* chromosome. In particular, for each cross and chromatid analyzed, we identified the genotype at 1-kb resolution along the chromosome and obtained all pairwise 1-kb vs 1-kb (two-marker) haplotypes as a function of the genomic distance. For recombinant haplotypes, we followed the convention of ordering ‘left’ and ‘right’ genotypes from telomere-proximal to centromere-proximal. We then combined the results from all chromatids for each cross sepparately, obtaining the frequency of the four possible haplotypes as a function of genomic distance. Finally, we transformed genomic distances into genetic distances (cM) based on the genetic map generated for each cross.

### The centromere effect

Two different methods were used to examine the centromere effect. The first method was performed following previous studies where the number of crossovers observed near the centromere (i.e., the centromere-proximal region) is compared to the expected number of crossovers for a region of equivalent size under the assumption that crossovers are randomly distributed along a chromosome arm [18, 20, 21, 41]. This approach requires a predetermined, arbitrary, size for the centromere-proximal genomic region, and in this study we defined it as one third of the chromosome arm, as in Miller *et al*. (2016) [41]. For each chromosome arm we obtained the probability of detecting centromere effect after comparing the observed and expected number of crossovers within the centromere-proximal region for that chromosome arm based on 10 million random replicates. To obtain a random distribution of crossover locations on a chromosome arm, we took into account the genomic locations were the detection of a crossover would have been possible near centromeres based on our genotyping method.

To study the centromere effect quantitively and identify the genomic region influenced by it, we applied a second method that estimates the proportion of the chromosome showing a significant reduction in crossovers. Starting at the most centromere-proximal region, we calculated the likelihood of a 1 Mb genomic region (window) showing a significantly reduced number of crossovers relative to the rest of the chromosome. This process was repeated after moving the window into the chromosome arm with 100 kb increments. For each window, we used a binomial distribution to determine if the number of crossovers was significantly reduced relative to expectations under the assumption that crossovers are randomly distributed along a chromosome arm. We obtained probability values for each 1 Mb window based on *n* trials of the binomial distribution, where *n* is the total number of crossovers in a chromosome arm with a probability equal to the fraction that this 1 Mb region represents of the total length of the chromosome arm where crossovers could be detected. Using these parameters, we obtained *P* values for each 1-Mb window moving from the centromere towards the center of the chromosomes, and identified the threshold of the genomic regions affected by the centromere effect when five consecutive windows showed a non-significant reduction in crossovers. Two levels of significance were examined, *P* < 1×10^-6^ and *P* < 1× 10^-2^. In both models, the total number of crossovers from the three crosses analyzed were used to investigate centromere effects. Equivalent analyses were used to detect the telomere effect.

To enable a direct comparison between *D. yakuba* and *D. melanogaster*, we also examined the centromere and telomere effect in *D. melanogaster* with the same methodologies as in our study of *D. yakuba*. Furthermore, we analyzed the *D. melanogaster* genome assemblies *r5.3* and *r6* to capture different levels of inclusion of peri-centromeric (or telomeric) sequences and heterochromatin. Unless noted, the *D. melanogaster r6* release is used by default in the main text, and results comparing *D. yakuba* to both *D. melanogaster r5.3* and *r6* are shown in Figures and Tables.

### Crossover interference

Crossovers tend to avoid peri-centromeric and peri-telomeric regions, and even within the ‘central’ region of chromosomes (outside the influence of centromere and telomere effects), crossovers are not randomly distributed. Therefore, the direct comparison of observed inter-crossover distance (ICD) in chromatids with two crossovers (2CO) and the expected distance based on a random distribution of crossovers along a chromosome arm can produce biased conclusions on interference. Another approach to measure interference is to assume that the distance between two independent crossovers in 2CO chromatids (no interference) is exponentially distributed and, as such, follows a gamma distribution with shape parameter (*ν*) of 1 [45]. Fitting the observed series of ICD in 2CO chromatids to a gamma distribution, therefore, would provide information about the existence of crossover interference when *ν* is greater than 1, evidencing crossovers more evenly spaced (positive interference) than expected under a Poisson process [45, 144, 238]. This model, however, also assumes that all genomic sites along a chromosome are equally likely to become a crossover, which is not correct in many species and may cause expectations for *ν* to be different than 1 under no interference.

To study crossover interference in *D. yakuba*, we considered that crossovers are not randomly distributed along chromosome arms. Expectations of ICD for 2CO chromatids under no interference can be obtained by using data from single crossover chromatids (1CO), randomly choosing two crossover locations along a chromosome and estimating the distance [39]. Our large number of genotyped meioses and 1CO chromatids (a total of 3,531) allows us to use this approach to obtain ICD expectations for 2CO chromatids after 1 million independent replicates per chromosome arm. Genome-wide analyses of crossover interference were generated by considering the differences between chromosome arms (observed number of 1CO and 2CO chromatids, and the distribution of crossovers along each arm in 1CO chromatids).

For the sake of allowing comparisons with other studies, we also show the estimated *ν* for each chromosome and expectations of *ν* under assumption of no interference based on distance between two randomly chosen 1COs. Note that the random generation of ICDs based on the location of 1COs shows a best fit to gamma distributions with *ν* greater than 1 (as discussed above). Estimates of *ν* for ICD of 2CO chromatids from our *D. yakuba* datasets are consistently greater than 1 and, more relevant, also consistently greater than our expectations, also supporting the presence of positive interference.

### Chromatid and tetrad analysis

Crossover events were divided into classes depending on the number of observed crossovers per chromatid: zero or non-crossover (NCOs), single crossover (1COs), double crossovers (2COs), triple crossovers (3COs), and quadruple crossovers (4COs). This information was also used to estimate the frequency of bivalent (or tetrad) exchange classes, initially based on Weinstein’s algebraic method [64, 65]. The frequency of tetrad classes is estimated as *E_r_*, denoting tetrads with *r* crossovers (*E_0_* tetrads indicate the frequency of homologous chromosome pairs with no crossovers; *E*_1_ tetrads with a single crossover, *E*_2_ tetrads with two crossovers, etc.) Note that Weinstein’s method is based on a meiotic model that assumes random distribution of crossovers along chromatids with no crossover interference, random selection of pairs of nonsister chromatids (no chromatid interference), random distribution of chromatids into gametes, and equal viability among all possible meitotic products. Under this model, the frequency of tetrad classes can be estimated using the observed frequency of crossover classes (NCO, 1CO, 2CO, 3CO and 4CO) and by solving *E*_r_,

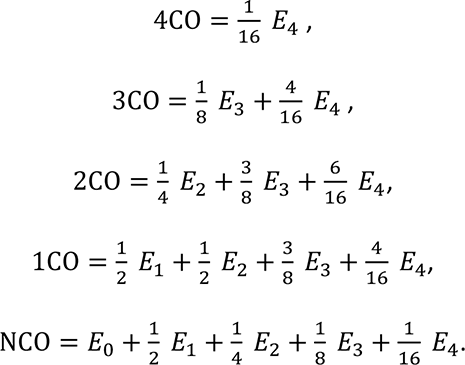

To obtain confidence intervals and compare models, a maximum likelihood (ML) approach was applied to identify by simulation the combination of tetrad frequencies (*E*_r_) that fits best to observed data (number of NCO, 1CO, 2CO, 3CO and 4CO chromatids) under several models. This allowed us to obtain point estimates for *E*_r_, confidence intervals and overall fit of a model to data. To compare models, we calculated the likelihood test statistic for each model and performed a likelihood-ratio test (LRT). Given our detection of rare but non zero chromosomes with 4 crossovers and a nonnegligible fraction of chromosomes with three crossovers, we modeled *E*_0_, *E*_1_, *E*_2_, *E*_3_ and *E*_4_. Models limiting *E*_≤2_ perform much worse than *E*_≤4_, at least for *D. yakuba*, and therefore are not used. Our simplest model assumed no restrictions in terms *E_r_* values, crossover distribution across chromatids, segregation or viability; it is equivalent to Weinstein’s expectations for a given maximum *r* (in our case *E*_≤4_).

We also used ML models to add a diverse set of rules [66, 145] to initially restrict expectations to biologically feasible values (*E*_r_ ≥ 0). We then expanded the rules of *E*_r_ ≥ 0 to study models including *meiotic drive* or *chromatid interference*. The model with *meiotic drive* assumes a non-random segregation of sister chromatids in meiosis II depending on whether sister chromatids had or not crossovers. We explicitly explored meiotic drive with a bias (*b*) favoring chromatids with crossovers preferentially segregating into the oocyte nucleus when the sister chromatid has no crossovers (MD_CO_ model). Under this model (see **Figure 6A**), chromatids with crossovers have a probability of being segregated into the oocyte nucleus of 0.5 ≤ *b* ≤ 1, instead of no bias (*b* = 0.5), which causes nonrecombinant sister chromatids to be preferentially extruded to the second polar body.

The model with *chromatid interference*, on the other hand, assumes that chromatids used for multiple exchanges in a tetrad with *E_≥2_* are not chosen independently and at random [64, 145]. This model allows for a biased distribution of crossovers (*b*) among chromatids when *E*_≥2_. In particular, we studied a model of *positive chromatid interference* (PCI) favoring (0.5 ≤ *b* ≤ 1) the sister chromatid with the fewest number of previous crossovers for each additional crossover, thus reducing the number of chromatids with no crossovers relative to a random case of *b* = 0.5.

To compare the observed frequency of different crossover and tetrad classes in *D. yakuba* (this study) and *D. melanogaster*, we used *D. melanogaster* data based on WGS [41] and visible markers [37, 239, 240]. We included results from visible markers in *D. melanogaster* because these studies are based on much larger sample sizes than that based on WGS for this species. On the other hand, it is worth noting that studies using visible markers are more likely to generate egg-to-adult viability defects than WGS and, because reduced density of markers along chromosomes, underestimate crossover events (hence overesting NCO chromatids and *E*_0_).

### Rates of protein evolution in the *D. melanogaster* subgroup

To identify variation in the efficacy of selection at protein level, we estimated *d*_N_ (the number of nonsynonymous substitutions per nonsynonymous site), *d*_S_ (the number of synonymous substitutions per synonymous site) and *ω* (the *d*_N_/*d*_S_ ratio) for each gene and across the entire phylogeny of the *D. melanogaster* subgroup (*D. melanogaster, D. simulans, D. sechellia, D. yakuba and D. erecta;* **Figure 7** and **Figure 8**). To this end, we applied the branch-model in the *codeml* program as implemented in PAML (v4.9j, February 2020) that allows different *ω* in all internal and external branches of the five-species tree [191, 192]. Furthermore, we took into account gene-specific properties influencing *ω* (CDS and transcript length, number of introns, gene expression, and CUB as an estimate of the degree of weak selection on synonymous mutations and therefore *d*_S_) and used residuals generated by Generalized Additive Models (GAM) as a measure of variable efficacy of selection on amino acid changes (*ω*_R_) for each gene and phylogenetic branch. We estimated rates of evolution for a curated set of 7,062 aligned orthologous CDS from flyDIVaS [241].

To identify genes under positive selection for amino acid changes in one or more branches, we compared Model M1a (nearly neutral evolution) against a model that allows the additional presence of positive selection at a fraction of sites (model M2a). For each branch, we compared maximum likelihood estimates (MLEs) under these two models and applied likelihood ratio tests (LRTs) to identify evidence of positive selection after correcting for multiple tests (FDR=0.05) [242]. To study the efficacy of selection against deleterious amino acid changes, we followed Larracuente *et al*. (2008) [104] and generated a set of genes with no evidence of positive selection for each branch. This set of genes was obtained by removing genes with the highest 10% *ω*_R_ as well as all those that showed significant signal of positive selection based on the LRT approach.

### Synonymous codon usage

We estimated the degree of bias in the use of different synonymous codons (Codon Usage Bias, CUB) by calculating ENC (Effective Number of Codons) [243] in genes with more than 200 codons aligned across species. ENC measures departures from random use among synonymous codons for each amino acid and has been shown to be minimally influenced by differences in CDS length and amino acid composition [244]. The values for ENC range between 20 (extreme CUB, with only one synonymous codon used per amino acid) and 59 (random use of synonymous codons) and, therefore, we used -ENC to evaluate the expected positive correlation between CUB and efficacy of selection.

### Neutral diversity in *D. yakuba* and *D. melanogaster*

We obtained estimates of neutral diversity based on pairwise sequence comparisons at four-fold synonymous sites (π_4f_). For *D. yakuba*, we analyzed the CY population (Cameroon; West Africa), which shows a high degree of long-term stability with median estimates of Tajima’s *D* [245] very close to equilibrium expectations (−0.08). For *D. melanogaster*, we analyzed the sub-Saharan African RG population from Rwanda (*Drosophila* Genome Nexus; http://johnpool.net/genomes.html [246, 247]), which combines a relatively large sample size (n=27), minimal signals of non-equilibrium dynamics, and low and well characterized levels of admixture [83, 247].

## Author Contributions

Conceived and designed the experiments: NP ALL JMC. Performed the experiments: NP ALL. Analyzed the data: NP ALL JMC. Contributed reagents and materials: ALL. Wrote the paper: NP ALL JMC.

## Data Availability

Sequence data have been deposited at the NCBI database (BioProject PRJNA751209).

## Supporting Information Captions

### Supplemental Materials and Methods

#### Supplemental tables

Meiotic, genomic and evolutionary properties of crossover distribution in *Drosophila yakuba* Nikale Pettie, Ana Llopart and Josep M. Comeron

**Supplemental Table 1.** Observed number of meiotic events for each of the three crosses of *D. yakuba* analyzed in this study

| *TZ043 x TZ020* |  |  |  |  |  |
| --- | --- | --- | --- | --- | --- |
|  | *X* | *2L* | *2R* | *3L* | *3R* |
| NCO<sup>1</sup> | 117 | 244 | 207 | 199 | 202 |
| 1CO | 290 | 232 | 147 | 237 | 273 |
| 2CO | 68 | 46 | 31 | 79 | 38 |
| 3CO | 27 | 1 | 11 | 7 | 5 |
| 4CO | 0 | 0 | 1 | 0 | 1 |
| Average CO / chromatid | 1.01 | 0.63 | 0.62 | 0.80 | 0.71 |

| *Sn20 x Sn17* |  |  |  |  |  |
| --- | --- | --- | --- | --- | --- |
|  | *X* | *2L* | *2R* | *3L* | *3R* |
| NCO<sup>1</sup> | 168 | 264 |  | 211 | 202 |
| 1CO | 281 | 220 |  | 278 | 278 |
| 2CO | 47 | 34 |  | 72 | 39 |
| 3CO | 18 | 0 |  | 3 | 3 |
| 4CO | 0 | 0 |  | 0 | 0 |
| Average CO / chromatid | 0.83 | 0.56 |  | 0.76 | 0.70 |

| *Rain5 x Cost1235.2* |  |  |  |  |  |
| --- | --- | --- | --- | --- | --- |
|  | *X* | *2L* | *2R* | *3L* | *3R* |
| NCO<sup>1</sup> | 162 | 327 | 256 | 284 | 279 |
| 1CO | 330 | 256 | 165 | 266 | 278 |
| 2CO | 85 | 36 | 19 | 58 | 30 |
| 3CO | 29 | 1 | 3 | 7 | 7 |
| 4CO | 0 | 0 | 1 | 0 | 0 |
| Average CO / chromatid | 0.97 | 0.53 | 0.49 | 0.66 | 0.60 |

<sup>1</sup> NCO: no crossovers, 1CO: single crossover, 2CO: double crossover, 3CO: triple crossover, 4CO: 4 crossovers in a single chromatid.

**Supplemental Table 2.**
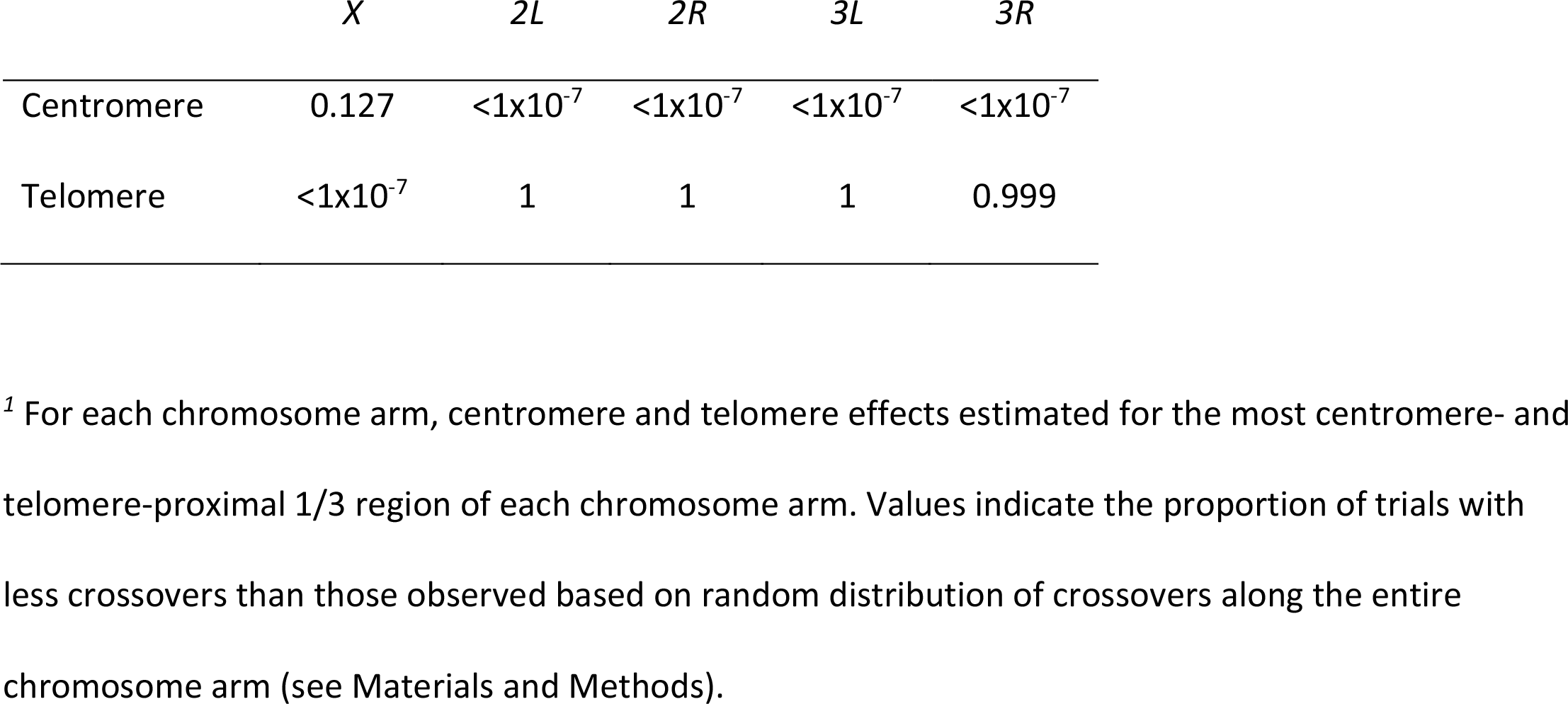
Centromere and telomere effect in *D. yakuba*

|  | *X* | *2L* | *2R* | *3L* | *3R* |
| --- | --- | --- | --- | --- | --- |
| Centromere | 0.127 | <1x10<sup>-7</sup> | <1x10<sup>-7</sup> | <1x10<sup>-7</sup> | <1x10<sup>-7</sup> |
| Telomere | <1x10<sup>-7</sup> | 1 | 1 | 1 | 0.999 |

<sup>1</sup> For each chromosome arm, centromere and telomere effects estimated for the most centromere- and telomere-proximal 1/3 region of each chromosome arm. Values indicate the proportion of trials with less crossovers than those observed based on random distribution of crossovers along the entire chromosome arm (see Materials and Methods).

**Supplemental Table 3.** Centromere and telomere effect in *D. yakuba* and *D. melanogaster*

| Centromere Effect |  |  |  |  |  |  |
| --- | --- | --- | --- | --- | --- | --- |
| Organism | Significance level | Proportion of Chromosome<sup>1</sup> |  |  |  |  |
|  |  | X | 2L | 2R | 3L | 3R |
| *D. yakuba* | $P < 1 \times 10^{-6}$ | 0.12 | 0.43 | 0.12 | 0.35 | 0.47 |
| | $P < 0.01$ | 0.13 | 0.50 | 0.15 | 0.39 | 0.52 |
| *D. melanogaster r6* | $P < 1 \times 10^{-6}$ | 0.12 | 0.08 | 0.22 | 0.24 | 0.25 |
| | $P < 0.01$ | 0.13 | 0.28 | 0.26 | 0.29 | 0.27 |
| *D. melanogaster r5.3* | $P < 1 \times 10^{-6}$ | 0.09 | 0.27 | 0.18 | 0.21 | 0.16 |
| | $P < 0.01$ | 0.13 | 0.29 | 0.27 | 0.24 | 0.27 |

| Telomere Effect |  |  |  |  |  |  |
| --- | --- | --- | --- | --- | --- | --- |
| Organism | Significance level | Proportion of Chromosome |  |  |  |  |
|  |  | X | 2L | 2R | 3L | 3R |
| *D. yakuba* | $P < 1 \times 10^{-6}$ | 0.12 | 0.04 | 0.03 | 0.00 | 0.04 |
| | $P < 0.01$ | 0.15 | 0.06 | 0.06 | 0.04 | 0.04 |
| *D. melanogaster r6* | $P < 1 \times 10^{-6}$ | 0.11 | 0.04 | 0.00 | 0.00 | 0.01 |
| | $P < 0.01$ | 0.12 | 0.04 | 0.00 | 0.03 | 0.02 |
| *D. melanogaster r5.3* | $P < 1 \times 10^{-6}$ | 0.12 | 0.00 | 0.02 | 0.00 | 0.01 |
| | $P < 0.01$ | 0.13 | 0.05 | 0.02 | 0.04 | 0.03 |

<sup>1</sup> Proportion of the chromosome with significant reduction in crossovers based on a sliding window study of a 1Mb with increments of 100kb (see Materials and Methods). For *D. melanogaster*, genome annotations r5.3 and r6 were analyzed. Two levels of significance are shown.

**Supplemental Table 4.** Satellite repeats in euchromatic and heterochromatic regions of *D. yakuba* and *D. melanogaster*

| Satellite | *D. yakuba* |  | *D. melanogaster* |  | *P* | Direction |
| --- | --- | --- | --- | --- | --- | --- |
|  | Euchrom. | Heterochrom. | Euchrom. | Heterochrom. |  |  |
| **AATAT** | 68091 | 8090 | 393911 | 92612 | $<1.8 \times 10^{-322}$ | M |
| **AAGAG** | 19 | 0 | 723191 | 108712 | $<1.8 \times 10^{-322}$ | M |
| **AACATAGAAT** | 0 | 0 | 44637 | 81886 | $<1.8 \times 10^{-322}$ | M |
| **AAGAT** | 0 | 0 | 95522 | 88427 | $<1.8 \times 10^{-322}$ | M |
| **ACCAGTACGGG** | 0 | 0 | 395 | 734 | $1.64 \times 10^{-247}$ | M |
| **AATAG** | 0 | 0 | 52896 | 39507 | $<1.8 \times 10^{-322}$ | M |
| C | 451494 | 20434 | 297575 | 25090 | $<1.8 \times 10^{-322}$ | Y |
| AAACAAT | 51037 | 5625 | 6849 | 58902 | $8.83 \times 10^{-149}$ | M |
| AAAATAT | 14189 | 1919 | 6868 | 6794 | $1.28 \times 10^{-45}$ | Y |
| AG | 5055 | 217 | 18402 | 7976 | $<1.8 \times 10^{-322}$ | M |
| AAT | 103831 | 7963 | 75269 | 2380 | $<1.8 \times 10^{-322}$ | Y |
| AT | 15149 | 509 | 56724 | 6425 | $<1.8 \times 10^{-322}$ | M |
| AGAT | 4717 | 61 | 81494 | 10024 | $<1.8 \times 10^{-322}$ | M |
| AC | 7824 | 29 | 5829 | 305 | $7.28 \times 10^{-48}$ | Y |
| AATATAT | 2983 | 0 | 75396 | 6956 | $<1.8 \times 10^{-322}$ | M |
| AAATTAT | 1791 | 55 | 7068 | 1254 | $<1.8 \times 10^{-322}$ | M |
| AATAATAT | 2076 | 0 | 25147 | 2323 | $<1.8 \times 10^{-322}$ | M |
| CG | 190 | 28 | 0 | 35 | $1.24 \times 10^{-30}$ | Y |
| AGAGGG | 154 | 0 | 39 | 59 | $4.19 \times 10^{-04}$ | Y |
| AAAACAATAACAAT | 110 | 8 | 9 | 60 | $3.39 \times 10^{-04}$ | Y |
| AAAACAAT | 119 | 14 | 9 | 60 | $6.70 \times 10^{-06}$ | Y |
| AAATAT | 1541 | 47 | 1766 | 256 | $5.07 \times 10^{-13}$ | M |
| AAG | 695 | 55 | 4715 | 1209 | $<1.8 \times 10^{-322}$ | M |
| AACAAT | 124 | 39 | 0 | 29 | $4.02 \times 10^{-22}$ | Y |
| AAATATAT | 704 | 0 | 4098 | 342 | $1.22 \times 10^{-187}$ | M |
| AAATAAT | 239 | 0 | 6620 | 1384 | $<1.8 \times 10^{-322}$ | M |
| AAAAG | 155 | 0 | 6649 | 1100 | $<1.8 \times 10^{-322}$ | M |
| AAC | 599 | 127 | 642 | 53 | $1.90 \times 10^{-17}$ | Y |
| AAAAATAT | 123 | 0 | 14 | 7 | $7.99 \times 10^{-25}$ | Y |
| AAAAAT | 1034 | 12 | 60 | 10 | $<1.8 \times 10^{-322}$ | Y |
| AATACC | 37398 | 64157 | 9 | 0 | $<1.8 \times 10^{-322}$ | Y |
| AATATATAT | 101 | 0 | 202 | 19 | $2.27 \times 10^{-11}$ | M |
| AATAGAATAT | 37279 | 4786 | 0 | 0 | $<1.8 \times 10^{-322}$ | Y |
| AAAGAAAT | 14740 | 7212 | 0 | 0 | $<1.8 \times 10^{-322}$ | Y |
| AATACAT | 13673 | 6907 | 0 | 0 | $<1.8 \times 10^{-322}$ | Y |
| ACTCTAT | 2087 | 0 | 0 | 0 | $<1.8 \times 10^{-322}$ | Y |

**Supplemental Table 4.** – continued
| Satellite | *D. yakuba* |  | *D. melanogaster* |  | *P* | Direction |
| --- | --- | --- | --- | --- | --- | --- |
|  | Euchrom. | Heterochrom. | Euchrom. | Heterochrom. |  |  |
| ACTGCT | 1961 | 0 | 0 | 0 | $<1.8 \times 10^{-322}$ | Y |
| ACATAT | 1403 | 0 | 0 | 0 | $4.68 \times 10^{-307}$ | Y |
| AATAGC | 1274 | 0 | 0 | 0 | $5.05 \times 10^{-279}$ | Y |
| ACCCAT | 1226 | 0 | 0 | 0 | $1.36 \times 10^{-268}$ | Y |
| AAAATAAAT | 1011 | 14 | 0 | 0 | $6.61 \times 10^{-225}$ | Y |
| AAGTAC | 773 | 0 | 0 | 0 | $4.00 \times 10^{-170}$ | Y |
| ACAGACGG | 680 | 0 | 0 | 0 | $6.68 \times 10^{-150}$ | Y |
| AAATAAATT | 617 | 34 | 0 | 0 | $1.35 \times 10^{-143}$ | Y |
| AAAAT | 245 | 304 | 0 | 0 | $2.08 \times 10^{-121}$ | Y |
| AACTAC | 547 | 0 | 0 | 0 | $5.66 \times 10^{-121}$ | Y |
| AAAAATT | 446 | 17 | 0 | 0 | $1.07 \times 10^{-102}$ | Y |
| AAATATG | 463 | 0 | 0 | 0 | $1.07 \times 10^{-102}$ | Y |
| AACAT | 385 | 0 | 0 | 0 | $1.01 \times 10^{-85}$ | Y |
| AACCT | 252 | 0 | 0 | 0 | $9.52 \times 10^{-57}$ | Y |
| AAACAG | 227 | 0 | 0 | 0 | $2.69 \times 10^{-51}$ | Y |
| ATCC | 229 | 0 | 0 | 0 | $9.85 \times 10^{-52}$ | Y |
| AAACATAC | 215 | 0 | 0 | 0 | $1.11 \times 10^{-48}$ | Y |
| AAGTAG | 196 | 0 | 0 | 0 | $1.56 \times 10^{-44}$ | Y |
| AAAAAAT | 54 | 141 | 0 | 0 | $2.58 \times 10^{-44}$ | Y |
| ACATCC | 192 | 0 | 0 | 0 | $1.16 \times 10^{-43}$ | Y |
| ACCACT | 182 | 0 | 0 | 0 | $1.77 \times 10^{-41}$ | Y |
| ACCTCC | 173 | 0 | 0 | 0 | $1.64 \times 10^{-39}$ | Y |
| ATATC | 158 | 0 | 0 | 0 | $3.09 \times 10^{-36}$ | Y |

<sup>1</sup> Number of repeats for euchromatic and heterochromatic regions in *D. yakuba* and *D. melanogaster*
based on the study of PacBio reads that do and do not map to the corresponding reference genomes,
respectively. Satellites in bold are centromeric and centromere-proximal satellite repeats in *D.*
*melanogaster* [1]. *P*, values of a $\chi^2$ test for the difference in the ratios of euchromatic and
heterochromatic satellite presence between the two species. Direction indicates which species has
more satellites: M, *D. melanogaster*; Y, *D. yakuba*.

**Supplemental Table 5.** Number of satellite repeats per Mb in heterochromatic and euchromatic PacBio reads of *D. yakuba* and *D. melanogaster*

|  | *D. yakuba* | *D. melanogaster* | Ratio<br>*D. yak./D. mel.* |
| --- | --- | --- | --- |
| Heterochromatin | 601.4 | 14867.9 | 24.7 |
| Euchromatin | 185.2 | 756.6 | 4.1 |
| Ratio Het./Eu. | 3.2 | 19.7 |  |

**Supplemental Table 6.**
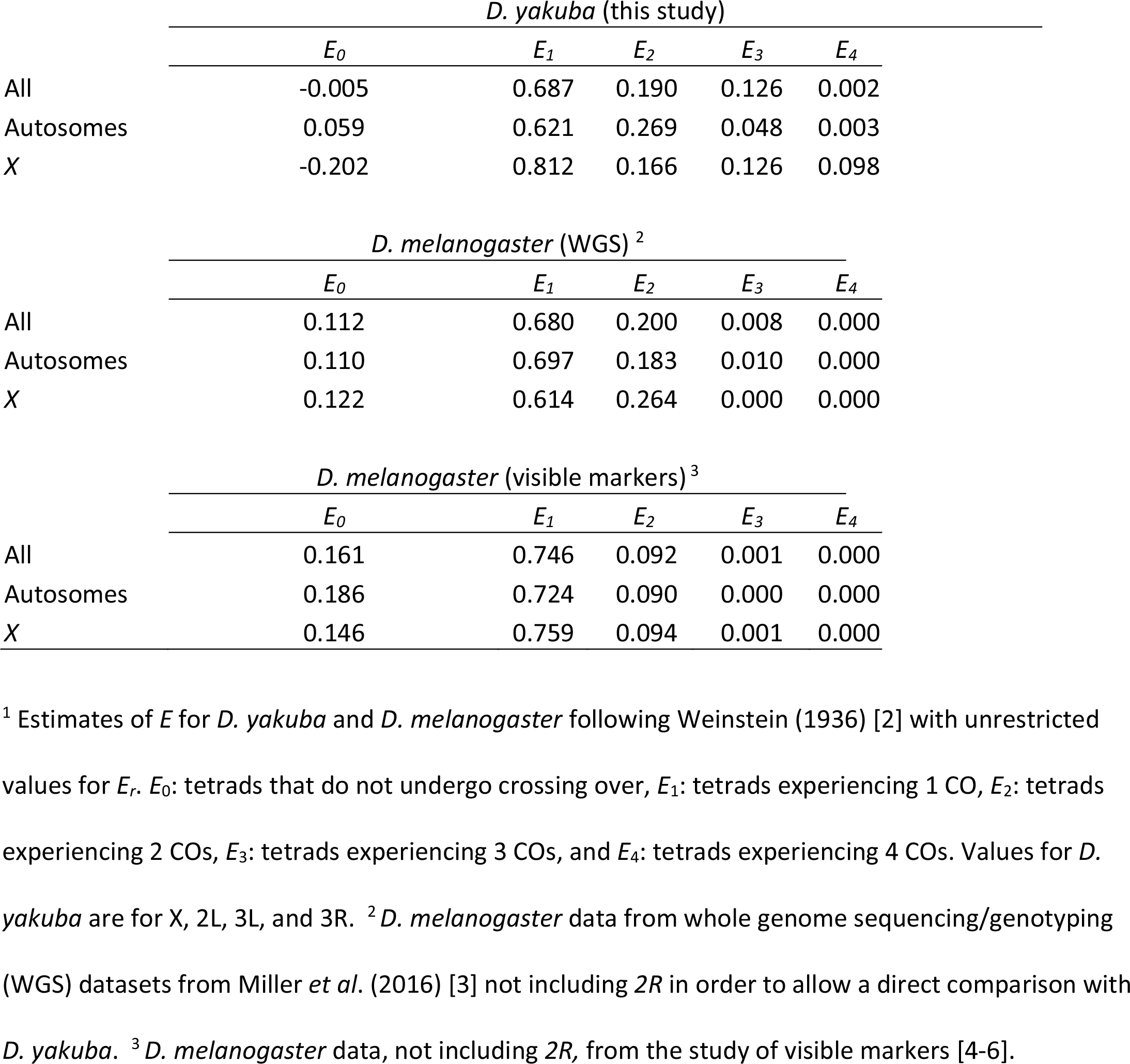
Tetrad analysis and estimates of *E* values for *D. yakuba* and *D. melanogaster*.

**Supplemental Table 7.**
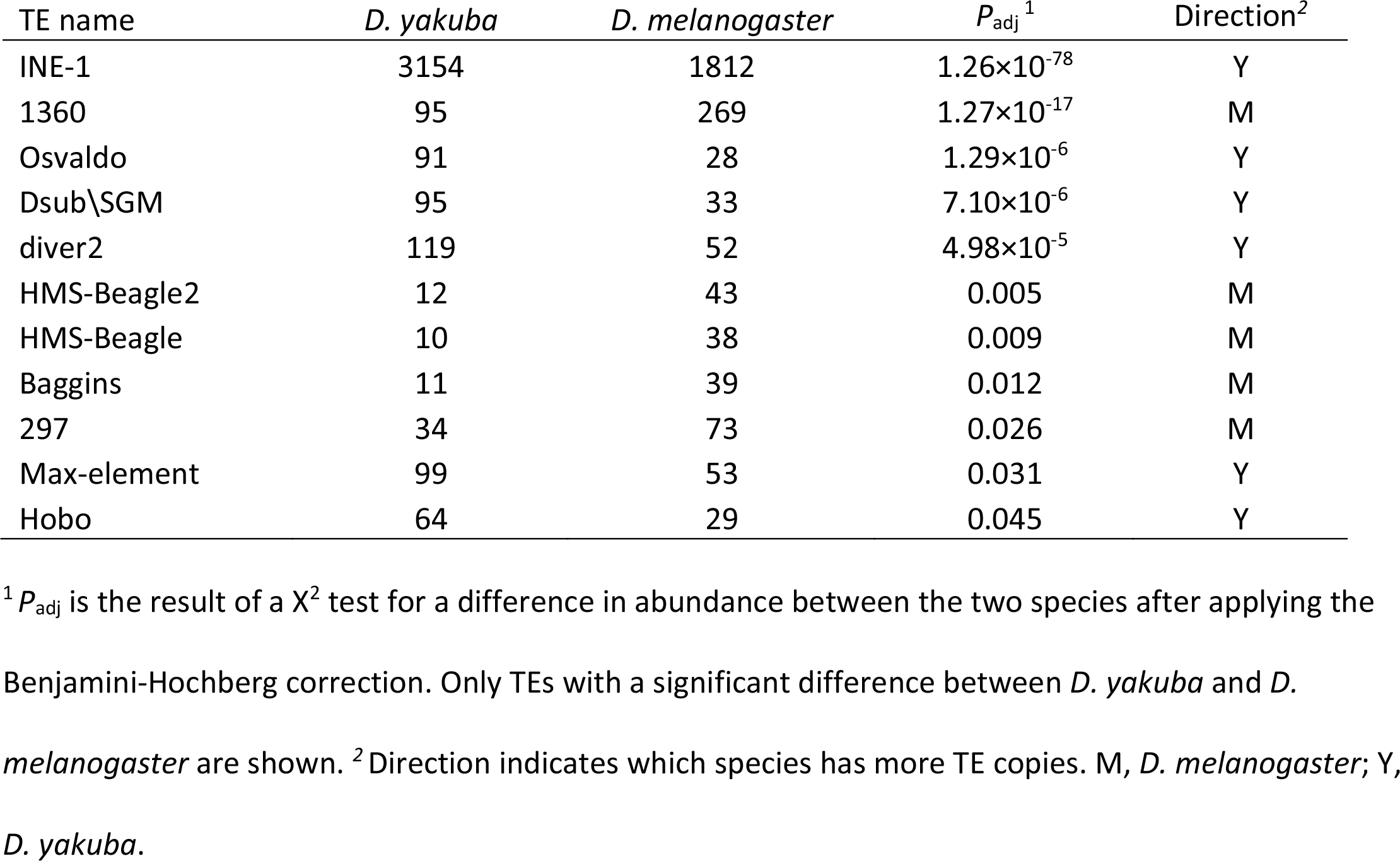
Transposable Elements (TEs) showing a significant difference in copy number between *D. yakuba* and *D. melanogaster*

| TE name | *D. yakuba* | *D. melanogaster* | $P_{\text{adj}}^1$ | Direction<sup>2</sup> |
| --- | --- | --- | --- | --- |
| INE-1 | 3154 | 1812 | $1.26 \times 10^{-78}$ | Y |
| 1360 | 95 | 269 | $1.27 \times 10^{-17}$ | M |
| Osvaldo | 91 | 28 | $1.29 \times 10^{-6}$ | Y |
| Dsub\SGM | 95 | 33 | $7.10 \times 10^{-6}$ | Y |
| diver2 | 119 | 52 | $4.98 \times 10^{-5}$ | Y |
| HMS-Beagle2 | 12 | 43 | 0.005 | M |
| HMS-Beagle | 10 | 38 | 0.009 | M |
| Baggins | 11 | 39 | 0.012 | M |
| 297 | 34 | 73 | 0.026 | M |
| Max-element | 99 | 53 | 0.031 | Y |
| Hobo | 64 | 29 | 0.045 | Y |

<sup>1</sup> $P_{\text{adj}}$ is the result of a $\chi^2$ test for a difference in abundance between the two species after applying the
Benjamini-Hochberg correction. Only TEs with a significant difference between *D. yakuba* and *D.*
*melanogaster* are shown. <sup>2</sup>Direction indicates which species has more TE copies. M, *D. melanogaster*; Y, *D. yakuba*.

**Supplemental Table 8.**
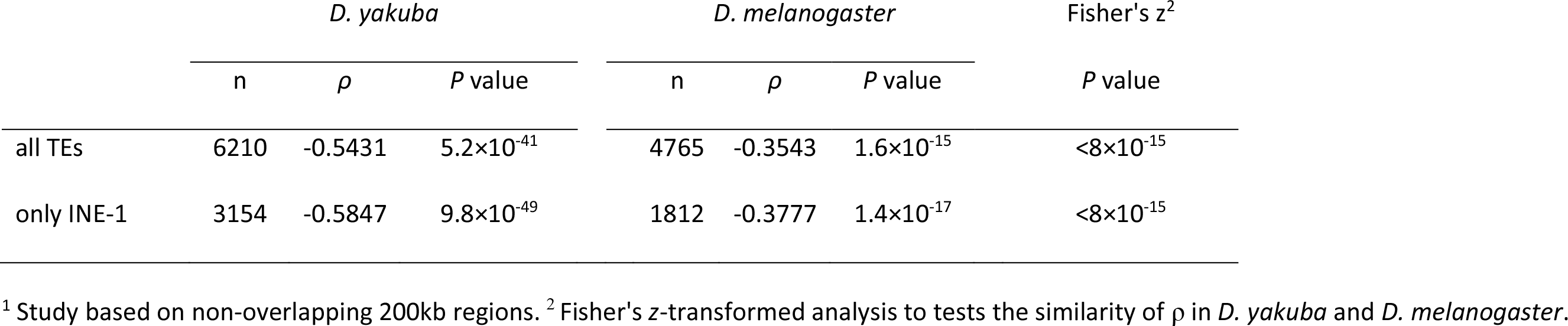
Spearman’s *ρ* correlation between TE abundance and crossover rate (cM/Mb) in *D. yakuba* and *D. melanogaster*

| | *D. yakuba* | | | *D. melanogaster* | | | Fisher's $z^2$ |
| --- | --- | --- | --- | --- | --- | --- | --- |
| | n | $\rho$ | *P* value | n | $\rho$ | *P* value | *P* value |
| all TEs | 6210 | -0.5431 | $5.2 \times 10^{-41}$ | 4765 | -0.3543 | $1.6 \times 10^{-15}$ | $< 8 \times 10^{-15}$ |
| only INE-1 | 3154 | -0.5847 | $9.8 \times 10^{-49}$ | 1812 | -0.3777 | $1.4 \times 10^{-17}$ | $< 8 \times 10^{-15}$ |

<sup>1</sup> Study based on non-overlapping 200kb regions. <sup>2</sup> Fisher's z-transformed analysis to tests the similarity of $\rho$ in *D. yakuba* and *D. melanogaster*.

**Supplemental Table 9.** Spearman’s *ρ* correlation between abundance of TE classes and crossover rate (cM/Mb) in *D. yakuba* and *D. melanogaster*

| TE Class | All Chromosomes |  |  |  |  |  |  |
| --- | --- | --- | --- | --- | --- | --- | --- |
|  | *D. yakuba* |  |  | *D. melanogaster* |  |  | Fisher's z |
| | N | P | P | N | $\rho$ | P | P |
| LTR | 1677 | -0.285 | $4.4 \times 10^{-11}$ | 1781 | -0.277 | $7.67 \times 10^{-10}$ | n.s. |
| TIR | 309 | -0.190 | $1.4 \times 10^{-05}$ | 177 | -0.329 | $1.66 \times 10^{-13}$ | n.s. |
| Line-like | 587 | -0.188 | $1.7 \times 10^{-05}$ | 538 | -0.263 | $5.59 \times 10^{-09}$ | n.s. |
| SINE | 3154 | -0.585 | $9.8 \times 10^{-49}$ | 1812 | -0.378 | $1.37 \times 10^{-17}$ | $< 8 \times 10^{-15}$ |
| non-LTR | 261 | -0.298 | $5.0 \times 10^{-12}$ | 295 | -0.276 | $8.76 \times 10^{-10}$ | n.s. |
| Foldback | 43 | -0.186 | $2.2 \times 10^{-5}$ | 69 | -0.124 | 0.0070 | n.s. |
| IR element | 82 | -0.311 | $4.4 \times 10^{-13}$ | 56 | -0.285 | $2.57 \times 10^{-10}$ | n.s. |
| MITE | 96 | -0.303 | $2.0 \times 10^{-12}$ | 30 | -0.323 | $5.01 \times 10^{-13}$ | n.s. |
| DNA | 1217 | -0.289 | $2.0 \times 10^{-11}$ | 178 | -0.326 | $3.13 \times 10^{-13}$ | n.s. |
| RNA | 4933 | -0.577 | $2.9 \times 10^{-47}$ | 4581 | -0.356 | $1.20 \times 10^{-15}$ | $< 8 \times 10^{-15}$ |

| TE Class | Autosomes |  |  |  |  |  |  |
| --- | --- | --- | --- | --- | --- | --- | --- |
|  | *D. yakuba* |  |  | *D. melanogaster* |  |  | Fisher's z |
| | N | P | P | N | $\rho$ | P | P |
| LTR | 1301 | -0.313 | $1.4 \times 10^{-10}$ | 1391 | -0.325 | $1.4 \times 10^{-10}$ | n.s. |
| TIR | 256 | -0.223 | $6.3 \times 10^{-06}$ | 157 | -0.347 | $6.4 \times 10^{-12}$ | n.s. |
| Line-like | 458 | -0.231 | $2.9 \times 10^{-06}$ | 489 | -0.324 | $1.6 \times 10^{-10}$ | n.s. |
| SINE | 2813 | -0.612 | $1.7 \times 10^{-42}$ | 1328 | -0.468 | $2.3 \times 10^{-21}$ | $9.4 \times 10^{-10}$ |
| non-LTR | 227 | -0.325 | $2.5 \times 10^{-11}$ | 204 | -0.282 | $3.1 \times 10^{-08}$ | n.s. |
| Foldback | 38 | -0.174 | $4.7 \times 10^{-04}$ | 40 | -0.138 | 0.0078 | n.s. |
| IR element | 77 | -0.309 | $2.5 \times 10^{-10}$ | 42 | -0.354 | $2.1 \times 10^{-12}$ | n.s. |
| MITE | 84 | -0.295 | $1.8 \times 10^{-09}$ | 15 | -0.342 | $1.3 \times 10^{-11}$ | n.s. |
| DNA | 998 | -0.330 | $1.3 \times 10^{-11}$ | 198 | -0.382 | $2.5 \times 10^{-14}$ | n.s. |
| RNA | 4257 | -0.613 | $1.2 \times 10^{-42}$ | 3469 | -0.429 | $4.7 \times 10^{-18}$ | $< 8 \times 10^{-15}$ |

**Supplemental Table 9.** – continued
| TE Class | Chromosome X |  |  |  |  |  |  |
| --- | --- | --- | --- | --- | --- | --- | --- |
|  | *D. yakuba* |  |  | *D. melanogaster* |  |  | Fisher's z |
|  | N | P | *P* | n | *p* | *P* | *P* |
| LTR | 376 | -0.136 | n.s. | 390 | -0.136 | n.s. | n.s. |
| TIR | 53 | -0.125 | n.s. | 20 | -0.325 | 0.0009 | n.s. |
| Line-like | 129 | -0.231 | 0.0125 | 49 | -0.085 | n.s. | n.s. |
| SINE | 341 | -0.379 | 2.7×10<sup>-05</sup> | 484 | -0.297 | 0.0021 | n.s. |
| non-LTR | 34 | -0.152 | n.s. | 91 | -0.301 | 0.0018 | n.s. |
| Foldback | 12 | -0.228 | 0.0147 | 15 | -0.130 | n.s. | n.s. |
| IR element | 219 | -0.246 | 0.0078 | 49 | -0.225 | 0.0210 | n.s. |
| MITE | 736 | -0.337 | 0.0002 | 1112 | -0.260 | 0.0073 | n.s. |
| DNA | 376 | -0.136 | n.s. | 390 | -0.136 | n.s. | n.s. |
| RNA | 53 | -0.125 | n.s. | 20 | -0.325 | 0.0009 | n.s. |

<sup>1</sup>TE classes except DNA and RNA are non-overlapping. Classes with less than 10 alignments in at least one species are not shown.

**Supplemental Table 10.**
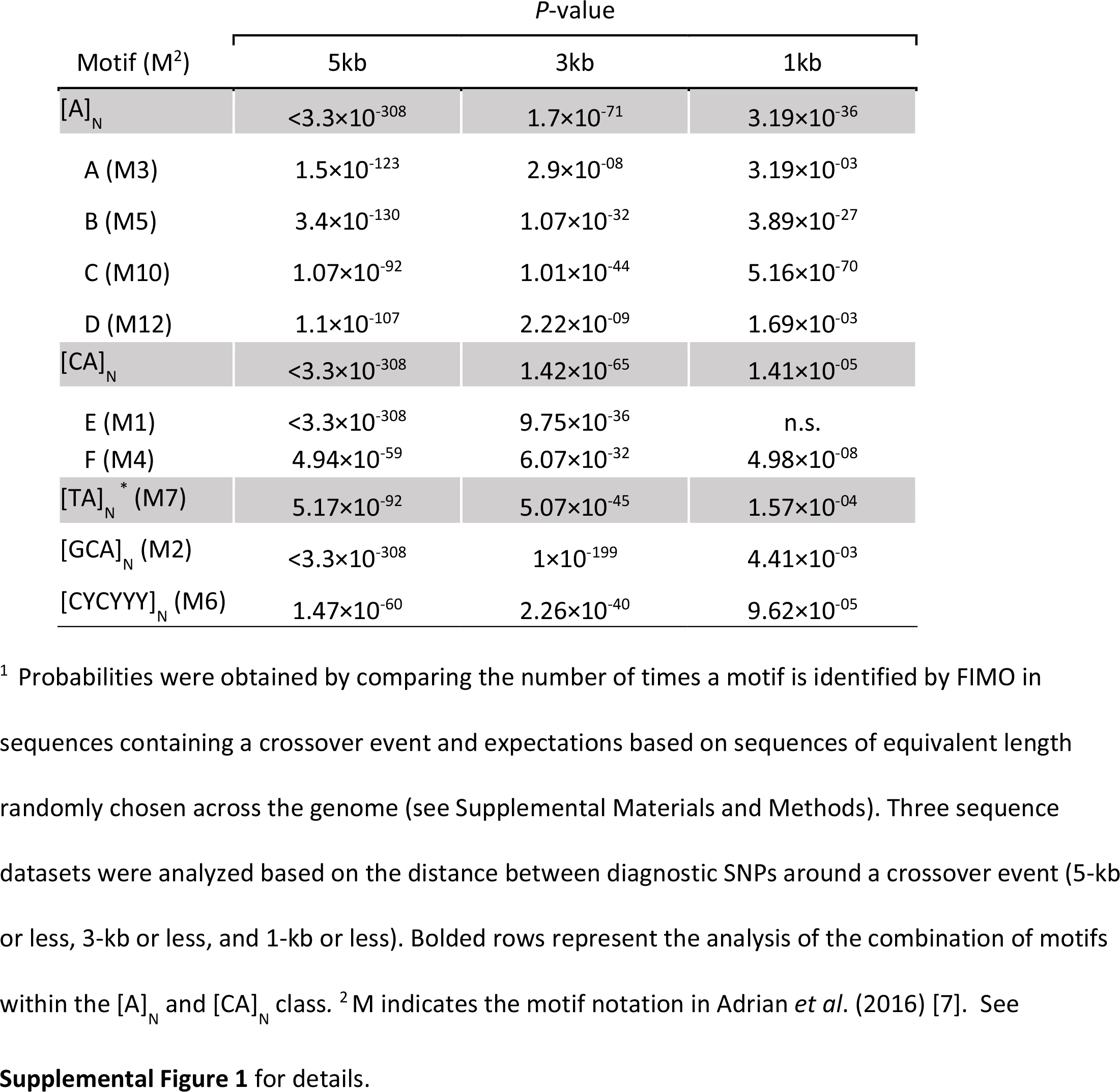
Enrichment analysis of individual motifs near crossover events in *D. yakuba*

| Motif (M<sup>2</sup>) | P-value |  |  |
| --- | --- | --- | --- |
|  | 5kb | 3kb | 1kb |
| **[A]<sub>N</sub>** | **<3.3×10<sup>-308</sup>** | **1.7×10<sup>-71</sup>** | **3.19×10<sup>-36</sup>** |
| A (M3) | 1.5×10<sup>-123</sup> | 2.9×10<sup>-08</sup> | 3.19×10<sup>-03</sup> |
| B (M5) | 3.4×10<sup>-130</sup> | 1.07×10<sup>-32</sup> | 3.89×10<sup>-27</sup> |
| C (M10) | 1.07×10<sup>-92</sup> | 1.01×10<sup>-44</sup> | 5.16×10<sup>-70</sup> |
| D (M12) | 1.1×10<sup>-107</sup> | 2.22×10<sup>-09</sup> | 1.69×10<sup>-03</sup> |
| **[CA]<sub>N</sub>** | **<3.3×10<sup>-308</sup>** | **1.42×10<sup>-65</sup>** | **1.41×10<sup>-05</sup>** |
| E (M1) | <3.3×10<sup>-308</sup> | 9.75×10<sup>-36</sup> | n.s. |
| F (M4) | 4.94×10<sup>-59</sup> | 6.07×10<sup>-32</sup> | 4.98×10<sup>-08</sup> |
| **[TA]<sub>N</sub><sup>\*</sup> (M7)** | **5.17×10<sup>-92</sup>** | **5.07×10<sup>-45</sup>** | **1.57×10<sup>-04</sup>** |
| [GCA]<sub>N</sub> (M2) | <3.3×10<sup>-308</sup> | 1×10<sup>-199</sup> | 4.41×10<sup>-03</sup> |
| [CYCYYY]<sub>N</sub> (M6) | 1.47×10<sup>-60</sup> | 2.26×10<sup>-40</sup> | 9.62×10<sup>-05</sup> |

<sup>1</sup> Probabilities were obtained by comparing the number of times a motif is identified by FIMO in sequences containing a crossover event and expectations based on sequences of equivalent length randomly chosen across the genome (see Supplemental Materials and Methods). Three sequence datasets were analyzed based on the distance between diagnostic SNPs around a crossover event (5-kb or less, 3-kb or less, and 1-kb or less). Bolded rows represent the analysis of the combination of motifs within the [A]<sub>N</sub> and [CA]<sub>N</sub> class. <sup>2</sup> M indicates the motif notation in Adrian *et al.* (2016) [7]. See

**Supplemental Table 11.** Data used to study tetrad frequencies in *D. melanogaster*

| CO Class | Chromosome |  |  |  |  |  |  |  |  |
| --- | --- | --- | --- | --- | --- | --- | --- | --- | --- |
| | $X^1$ | $2L^1$ | $2R^1$ | $3L^1$ | $3R^1$ | $X^2$ | $2L^3$ | $2L^2$ | $2R^4$ |
| NCO | 97 | 99 | 101 | 97 | 99 | 1,015 | 6,308 | 2,376 | 1,698 |
| 1CO | 86 | 88 | 90 | 84 | 86 | 964 | 4,909 | 1,758 | 1,150 |
| 2CO | 13 | 9 | 5 | 15 | 10 | 197 | 276 | 88 | 73 |
| 3CO | 0 | 0 | 0 | 0 | 1 | 3 | 2 | 0 | 0 |
| Total | 196 | 196 | 196 | 196 | 196 | 2,179 | 11,495 | 4,222 | 2,921 |

<sup>1</sup> Miller *et al.* (2016) [3], <sup>2</sup> Hatkevich *et al.* (2017) [4], <sup>3</sup> Baker and Carpenter (1972) [5], <sup>4</sup> Parry (1973) [6]. NCO, noncrossover; 1CO, single crossover; 2CO, double crossover; 3CO, triple crossover.

## Supplemental Figures

**Supplemental Figure 1.**
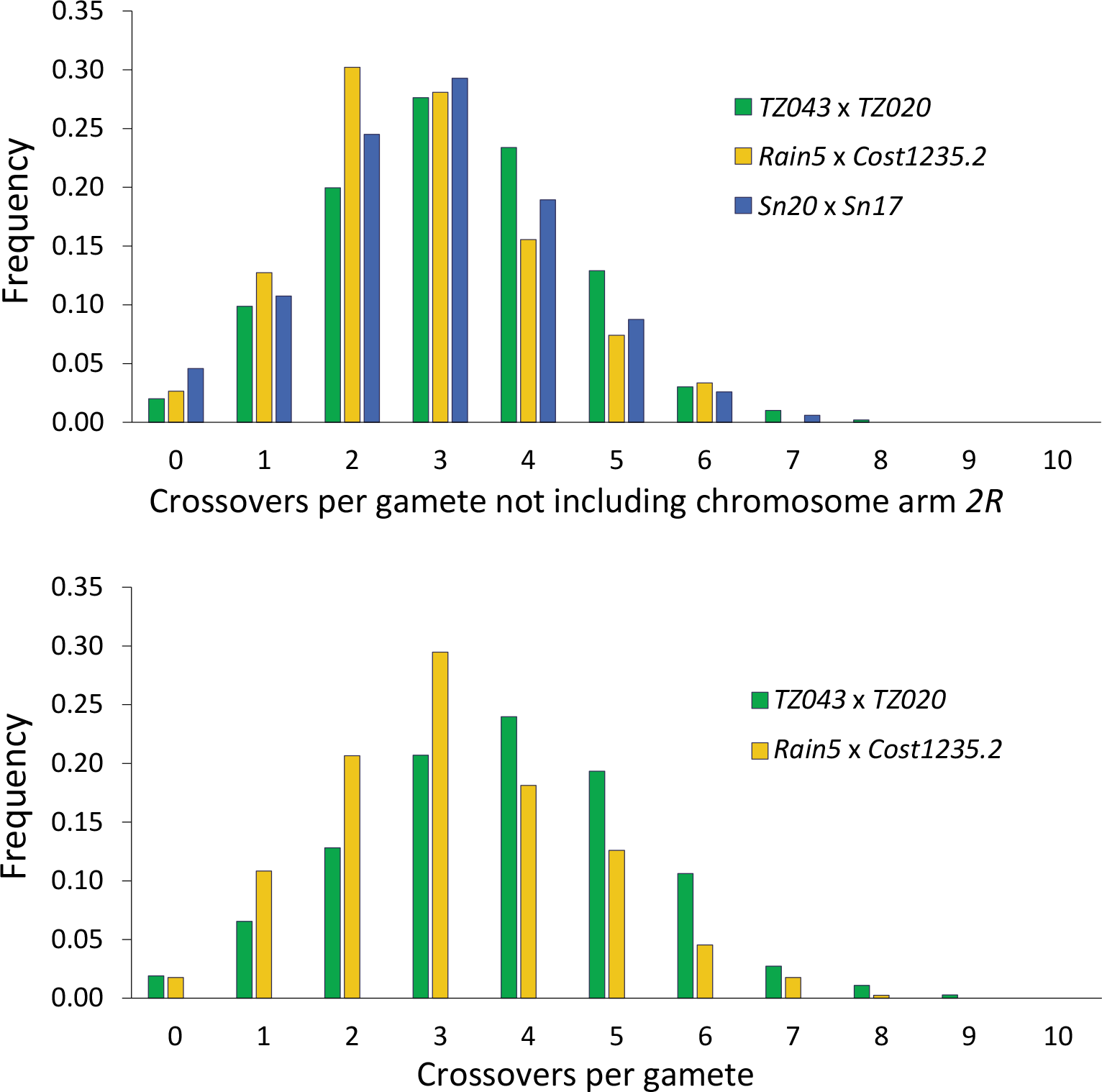
Frequency distribution of number of crossovers per gamete. A) Crossovers observed when not including data from chromosome arm *2R*, and B) crossovers for all chromosome arms for the two crosses not showing *2R* inversion heterozygosity (*TZ043* x *TZ020* and *Rain5* x *Cost1235.2*).

**Supplemental Figure 2.**
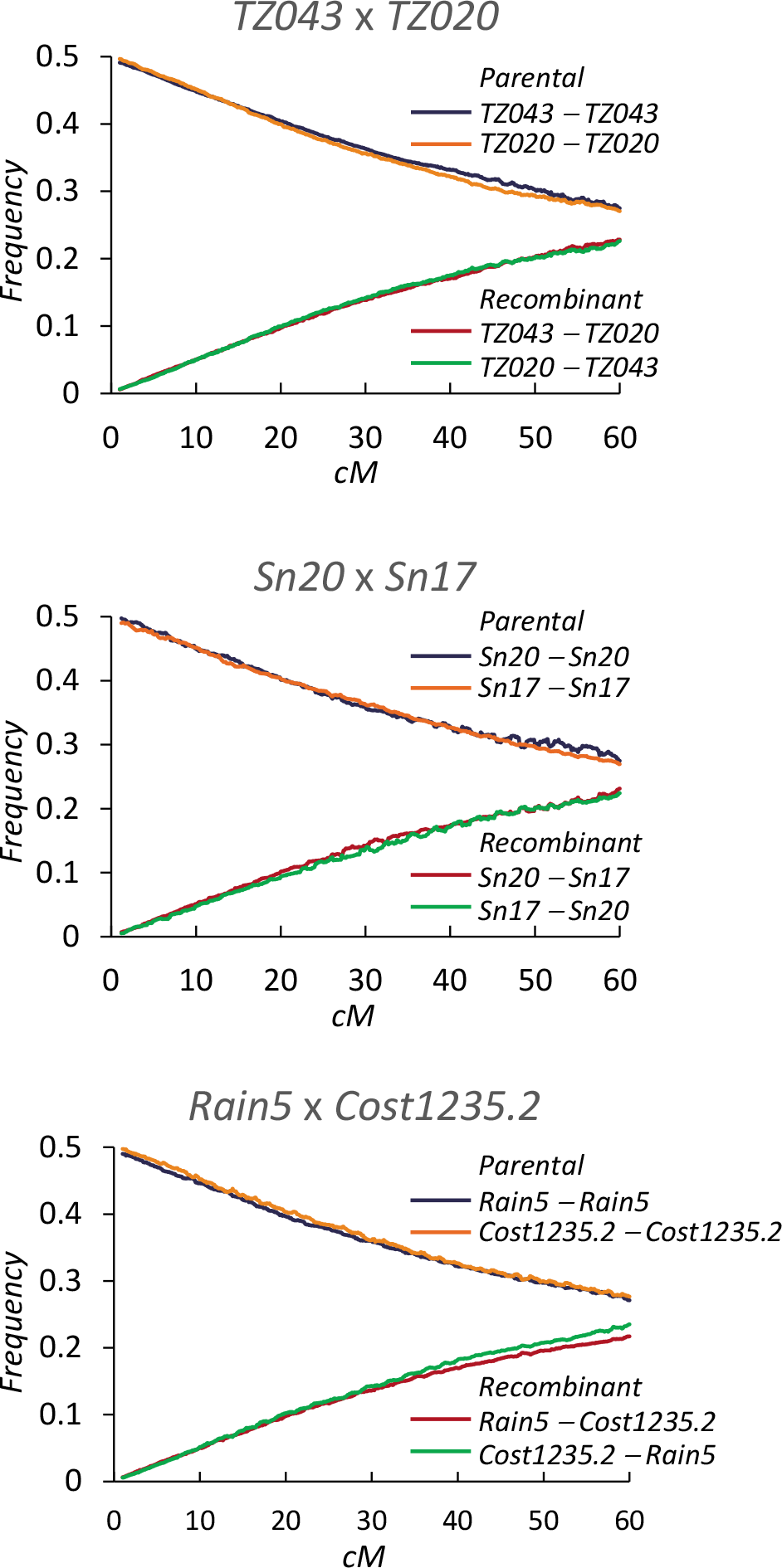
Frequency of parental and recombinant haplotypes as a function of genetic distance (cM) along the *X* chromosome. Frequencies shown as average within overlapping 1-cM windows with increments of 0.1 cM. For recombinant haplotypes, the order of the two parental genotypes is shown from telomere to centromere.

**Supplemental Figure 3.**
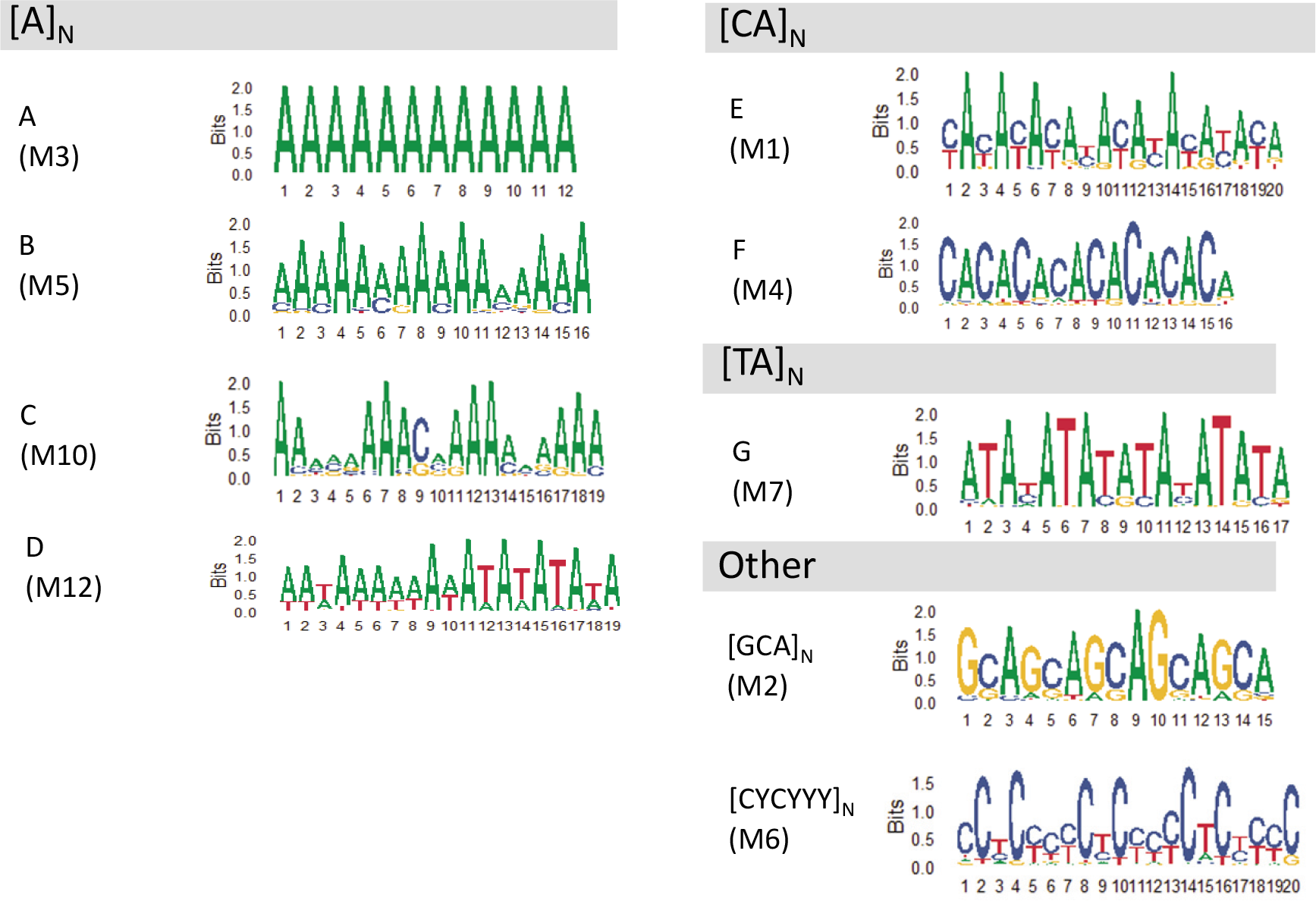
Motif logos for each of the short DNA motifs associated with crossovers in *D. yakuba*. Logos are presented with positions (y-axes) weighted by information content (x-axes) per site. Logos and notation in parenthesis are from Adrian *et al.* (2016).

**Supplemental Figure 4.**
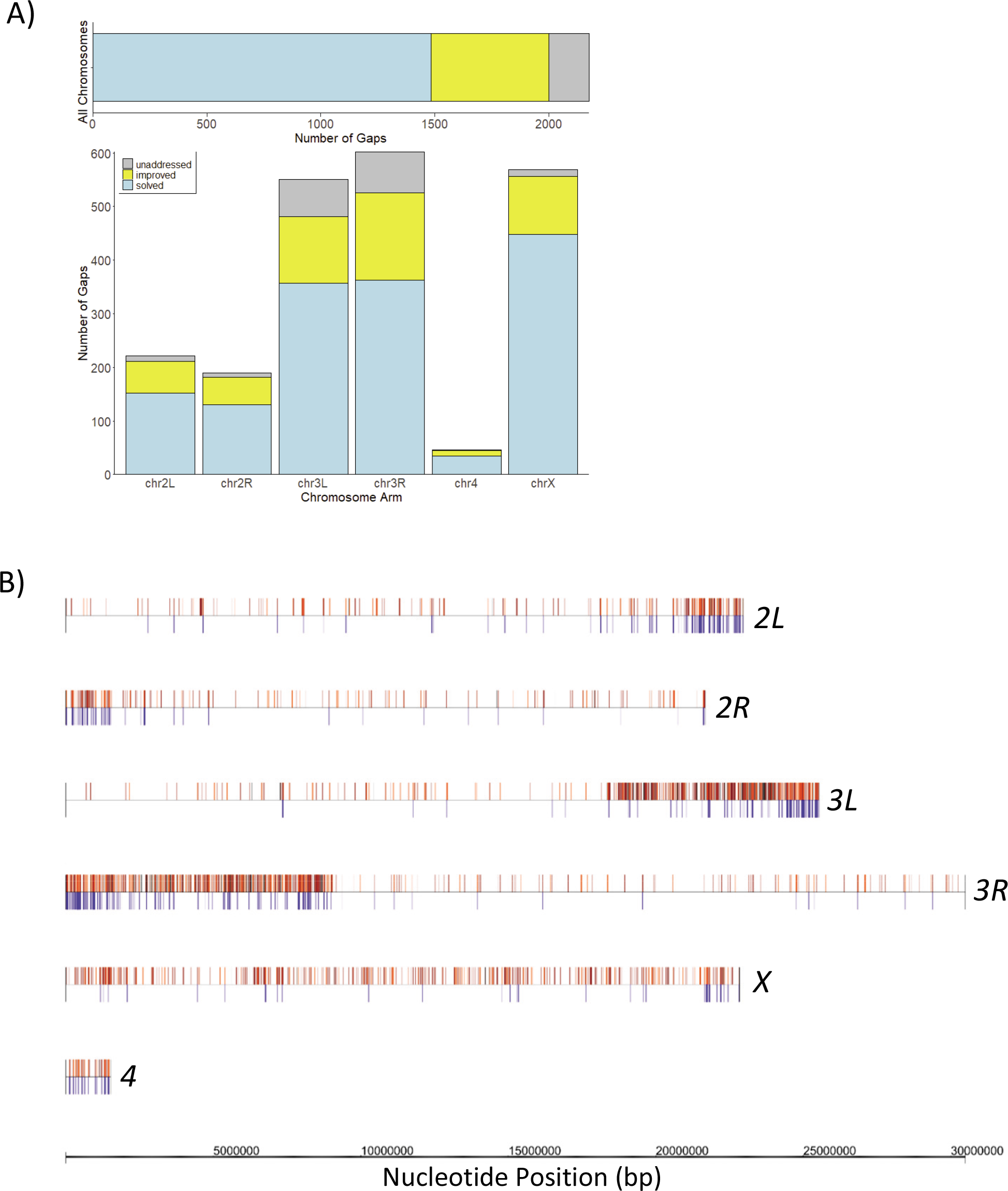
**A)** Description of gaps in the *D. yakuba* reference genome 2.0 that have been completely solved or improved in this study. **B)** Genomic locations of completely solved (dark red lines) or improved (light red lines) gaps. TE presence in added sequences are indicated as blue lines.

## Supplemental Materials and Methods

### Generation of an updated *D. yakuba* genome sequence

The *D. yakuba* reference genome release 2.0 [1] contains ∼5,400 gaps, 2,178 of them along the euchromatic genomic regions assigned to *2L*, *2R*, *3L* and *3R* chromosome arms, and the *X* and the dot chromosomes. This reference genome is based on the *D. yakuba* Tai18E2 line, which was established after multiple rounds of single pair (brother/sister) matings. In order to generate a more accurate genetic map for *D. yakuba,* and focusing on the euchromatic regions of the genome, we pursued filling gaps by PacBio (Pacific Biosciences) sequencing of the same Tai18E2 line maintained in our laboratory. High-molecular-weight DNA was extracted from adult females using standard protocols [2] and sent to the University of Washington PacBio Sequencing Services for library preparation and subsequent sequencing (https://pacbio.gs.washington.edu/). The sequences of PacBio reads used to fill gaps were corrected using illumina sequencing (see below).

### PacBio contigs and gap filling

Raw PacBio reads from two PacBio SMRT cells (4 lanes) were filtered using bash5tools version 0.8.0 with a minimum read length of 500 and a minimum read score of 0.75, generating approximately 10x coverage of the *D. yakuba* reference genome. Contigs were assembled using canu v.1.8 [3], and PBJelly2 version 15.8.24 [4] was used for gap filling. All stages in the PBJelly2 pipeline were done using the defaults, with the exception of the setup and assembly stages. In the setup stage, two different values were used for the minimum size of gap to take into account that gaps in the reference sequence can either represent unknown sequences between supercontigs or otherwise between contigs (which can be much smaller). In this regard, we also treated these two categories of gaps differently for the maximum number of nucleotides allowed to be added or subtracted, set to 5,000 or 1,000 for large and small gaps, respectively. Additionally, two different percent identity parameters were used for both gap categories, 70 (the program default) and 65 percent, resulting in four outputs from PBJelly2. A python program was written to merge outputs, sequentially checking for gaps filled or improved with 70 percent identity and, if not, considering whether the gap was filled or improved with the 65 percent identity parameter. If a gap was not addressed at either percent identity, the gap sequence in the *D. yakuba* Release 2.0 was mantained.

### Sample and Illumina library preparation

DNA was extracted from adult flies of Tai18E2 using a Tissue Lyser LT and the DNeasy Blood & Tissue Kit (Qiagen). After assessing concentration and quality on a NanoDrop One spectrophotometer (Thermo Scientific), the DNA was sheared using the Bioruptor UCD-200 (Diagenode) on the ‘High’ setting with 15 second pulse followed by 45 seconds of rest. This was repeated for a total of 24 cycles for each sample, and sheared samples were recovered using MinElute columns (Qiagen). Illumina libraries were prepared using the NEBNext DNA Library Prep Master Mix Set for Illumina (New England Biolabs). Samples were run on a 2% agarose gel for size selection (400-450 bp) and, after PCR enrichment, were purified with AMPure XP beads (Beckman-Coulter Life Sciences). Sequencing was carried out in an Illumina HiSeq 4000 at the Iowa Institute of Human Genetics (IIHG; University of Iowa) (https://medicine.uiowa.edu/humangenetics/genomics-division).

### Illumina alignment pipeline

After sequencing, reads were filtered using Trimmomatic-0.36 [5] with average quality 20, trailing quality 12 and minimum length 30. Filtered reads were mapped to the updated *D. yakuba* reference created with the PacBio reads using Bowtie2 version 2.1.0 [6], and unaligned reads were mapped again to the same reference using Stampy version 1.0.31 [7]. The aligned reads of both Bowtie2 and Stampy were then sorted and indexed using Samtools version 1.3.1 [8]. A Genome Analysis Toolkit pipeline (GATK version 3.5-0-g36282e4) was used to identify SNPs, indels and regions that needed realignment [9]. These regions were first identified by the RealignerTargetCreator and realigned with the IndelRealigner with the maximum number of reads allowed for alignment of 200,000 and all other default parameters. Finally, SNPs and indels were then called with the UnifiedGenotyper [9] with a minimum base quality score of 31; for indels we also required a minimum number of reads of 3 and a minimum fraction of reads supporting the indel of 0.51. Heterozygous sites and SNPs with allele frequency less than 0.75 and with a normalized Phred Score probability of heterozygote (het PL) < 50 were not considered at this time. FastaAlternateReferenceMaker [9] was used to create an intermediate reference genome that updated only PacBio sequences. BCFtools consensus version 1.3.1 [10] created a chain file that was used by CrossMap version 0.2.6 [11] to obtain a lift table with the new genome coordinates.

Following Lack *et al*. (2015) [12], we performed an additional round of mapping using the intermediate reference genome, which contained indels and SNPs. This second round was performed as described above using again Bowtie2 [6] and Stampy [7]. The output of CrossMap was used to filter out variants that were not located in PacBio sequences, and BCFtools consensus [10] was used to create the final chain file, which converted the coordinates of the intermediate reference to the coordinates of the new updated reference for *D. yakuba*. FastaAlternateReferenceMaker from GATK was used to create the final reference. As expected, sequence improvement with deep Illumina sequencing after two rounds of mapping allowed correcting more nucleotides of PacBio sequences (twice as many) than after one round of mapping, exemplifying the advantages of adding indels before a second mapping [12]. In all, more than 7,000 bases of the added PacBio sequences were corrected after both rounds of mapping. A final lift table utilizing pyliftover version 0.3 was constructed to allow direct comparison between the *D. yakuba* reference genome Release 2.0 and our updated (improved) *D. yakuba* genome sequence.

### Improved *D. yakuba* reference genome

Out of the original 2,178 existing gaps in the *D. yakuba* reference genome release 2.0 for *2L*, *2R*, *3L* and *3R* chromosome arms, and *X* and dot chromosomes, our PacBio and Illumina sequencing completely solved 1,483 gaps (approximately 68% of all gaps) and improved 517 (24%) additional gaps, for a total of more than 90% of gaps addressed (**Supplemental Figure 2A**). Most of the added sequence was in the centromere-proximal regions of chromosomes *2* and *3*, whereas the distribution of added sequences was more uniform in chromosomes *X* and *4* (**Supplemental Figure 2B**). We used this improved *D. yakuba* reference genome for all of our analyses.

We also explored the presence of transposable elements (TEs) in the sequences added to the reference using known *Drosophila* TEs [13] (see ’TE presence across the *D. yakuba* genome’ for details). As expected, we identified an enrichment in TEs along the added sequences relative to the reference genome release 2.0 [1] sequence (*P* < 2.2×10^-16^).

### Identification of chromosomal inversions

The presence of polymorphic inversions within *D. yakuba* is known [14, 15] and we first recognized inversions relative to the reference *D. yakuba* assembly visually as sharp peaks of crossover rates, representing inversion break points. We identified inversions relative to the reference assembly on both arms of chromosome *2* in our parental lines. The inversion on *2L* was present in all of our six parental genomes and therefore is homozygous in F_1_ females. A complex inversion within an inversion was identified on *2R*, consistent with the *2Rjk* inversion [15]. This complex *2R* inversion was homozygous in two of our crosses but heterozygous in F_1_ females for the cross *Sn20* x *Sn17*. Consistent with previous studies that show that heterozygous inversions suppress crossing over [16, 17], we did not detect crossovers within this complex inversion and data for chromosome arm *2R* from the cross *Sn20* x *Sn17* was not used in any of the analyses. For homozygous inversions we identified crossover events after inverting the location of diagnostic SNPs within the inversion, but show crossover locations and rates (**Figure 2** and **Figure 3**) based on the *D. yakuba* reference assembly order for simplicity.

### TE presence across the *D. yakuba* genome

To determine the distribution of TEs across the genome of *D. yakuba*, we used the sequences of known TEs from the Berkeley Drosophila Genome Project (BDGP; https://www.fruitfly.org/) [13] and aligned them to our improved *D. yakuba* reference using dc-megablast with default settings [18, 19]. The output from dc-megablast was then filtered to consider instances of different TEs aligned to the same genomic location due to sequence similarity, and the TE with the lowest Expected Value (*E*) was used. The BDGP TE database contains TEs mostly from *D. melanogaster* but also from other *Drosophila* species, including instances with the same TE characterized in different species. In these cases, we merged information to obtain a single identification per TE. Given previous studies indicating a significant presence of INE-1-like TEs in *D. yakuba* [20-24], we obtained a *D. yakuba* specific INE-1 sequence following Yang *et al*. (2006) [24]. Using the INE-1 sequences from the TE BDGP database, we identified the target sequences in the *D. yakuba* genome, aligned these target sequences to create a contig and obtained a consensus INE-1 sequence for *D. yakuba*.

To allow for a direct comparison between *D. yakuba* and *D. melanogaster,* we obtained data for TE presence in *D. melanogaster r5.3* and r*6* genome releases applying the same methodology as in *D. yakuba*. To identify differences in TE presence between *D. yakuba* and *D. melanogaster,* we applied a χ^2^ test to all TEs with more than 10 copies in at least one species. Unless noted, non-overlapping 200kb windows were used to obtain a distribution of TE presence across genomes.

### Lack of evidence of P-elements in *D. yakuba*

Within the *D. melanogaster* subgroup, P-elements have been identified in *D. melanogaster* and more recently in *D. simulans,* but not in the sister species *D. sechellia* or *D. mauritiana* [25-29]. Initial analyses of TE distribution across the *D. yakuba* genome recovered a few alignments to P-elements, possibly suggesting a recent invasion of P-elements in *D. yakuba*. We expanded this study by using Illumina reads from multiple lines of *D. yakuba* and Bowtie2 to recover alignments to the *D. melanogaster* P-element. The alignments to the P element were consistently limited to only small regions of exons 1 and 3. The absence of alignments to exons 0 and 2 suggests that there is no active P-element in *D. yakuba* because all four exons are required for transposition (reviewed in [30]). Furthermore, alignments to exon 1 cluster in an approximately 202bp portion in the center of the exon and all alignments to exon 3 cluster in a narrow region of ∼114 bp.

Notably, this same small region shows high sequence similarity with INE-1, and alignments of Illumina reads from an experimental profiling of *Drosophila* microRNAs [31-33] also show mapping to the regions of similarlity between INE-1 and P-elements. Combined, our analyses do not support the presence of P- elements in *D. yakuba*, either active or in recent past. Instead, the results indicate that recovered partial alignments are the consequence of sequence similarity with other TEs, including INE-1.

### Satellite repeats and TE presence in heterochromatic regions

The *D. yakuba* reference genome does not include heterochromatin and the sequences of centromere and telomere regions are unknown. Therefore, we used PacBio reads and assumed that reads that do not align to the updated *D. yakuba* reference genome (nuclear and mitochondrial) are mostly heterochromatic. To determine if satellite repeats enriched near centromeres in *D. melanogaster* were enriched also in *D. yakuba*, we investigated the presence of k-mers in PacBio reads of *D. yakuba* and *D. melanogaster*. Raw PacBio reads for *D. melanogaster* were obtained from Kim *et al.* (2014) [34] and analyzed and filtered using the same approach as that in *D. yakuba*. PacBio reads were aligned with BLASR [35] to our *D. yakuba* reference and to *D. melanogaster* r5.3 and r6 references. After alignment, mapped and unmapped reads were considered euchromatic and heterochromatic, respectively, and cut into 125bp sequences to find kmers with k-seek [36, 37]. For each repeat, a χ^2^ test was performed to compare the ratio of kmers aligned to heterochromatic PacBio reads in *D. yakuba* and *D. melanogaster* and the Benjamini-Hochberg method was applied to correct for multiple tests [38].

To examine the degree of TE enrichment in heterochromatic or euchromatic regions of *D. yakuba* and *D. melanogaster*, PacBio reads were used to create blast databases. The BDGP TE database was then aligned to these databases using dc-megablast. A goodness of fit test was performed to compare the number of alignments in the heterochromatic and euchromatic reads of each species, and the Benjamini-Hochberg correction was applied.

### Short DNA motifs and crossover localization

We investigated the presence of short DNA motifs previously reported to be predictive of changes in crossover rates across the *D. melanogaster* genome [39-41]. For each of these motifs, we used FIMO [42] and the position probability matrix (PPM) reported in [39] to annotate motif distribution across the *D. yakuba* genome. To investigate whether motif presence was enriched near crossover events, we took into account the variable distance between diagnostic SNPs flanking crossover events and the unknown precise location of crossovers. We, therefore, created three datasets of sequences containing crossovers defined by the different distance between diagnostic flanking SNPs (1kb or less, 3kb or less, and 5kb or less) and, unless noted, used the dataset of 1kb (or less) by default. For this analysis, chromosome arm *2R* was not included due to the smaller sample size relative to the other chromosome arms. To quantify enrichment, we applied FIMO to a set of randomly chosen genomic sequences of identical length to each of the sequences with crossovers, and this random selection was repeated 1,000 times to create a null expectation.

## REFERENCES

1. Baudat F, Imai Y, de Massy B. Meiotic recombination in mammals: localization and regulation. Nature reviews Genetics. 2013;14(11):794–806. doi: 10.1038/nrg3573. PubMed PMID: 24136506.

2. Lam I, Keeney S. Mechanism and regulation of meiotic recombination initiation. Cold Spring Harbor perspectives in biology. 2015;7(1):a016634. doi: 10.1101/cshperspect.a016634. PubMed PMID: 25324213; PubMed Central PMCID: PMC4292169.

3. Bergerat A, de Massy B, Gadelle D, Varoutas P-C, Nicolas A, Forterre P. An atypical topoisomerase II from archaea with implications for meiotic recombination. Nature. 1997;386(6623):414–7. doi: 10.1038/386414a0.

4. Keeney S, Giroux CN, Kleckner N. Meiosis-Specific DNA Double-Strand Breaks Are Catalyzed by Spo11, a Member of a Widely Conserved Protein Family. Cell. 1997;88(3):375–84. doi: https://doi.org/10.1016/S0092-8674(00)81876-0.

5. McKim KS, Jang JK, Manheim EA. Meiotic recombination and chromosome segregation in Drosophila females. Annu Rev Genet. 2002;36:205–32. Epub 2002/11/14. doi: 10.1146/annurev.genet.36.041102.113929. PubMed PMID: 12429692.

6. Hunter N. Meiotic Recombination: The Essence of Heredity. Cold Spring Harbor perspectives in biology. 2015;7(12). doi: 10.1101/cshperspect.a016618.

7. Oliver TR, Feingold E, Yu K, Cheung V, Tinker S, Yadav-Shah M, et al. New Insights into Human Nondisjunction of Chromosome 21 in Oocytes. PLOS Genetics. 2008;4(3):e1000033. doi: 10.1371/journal.pgen.1000033.

8. Nagaoka SI, Hassold TJ, Hunt PA. Human aneuploidy: mechanisms and new insights into an age-old problem. Nature reviews Genetics. 2012;13(7):493–504. Epub 2012/06/19. doi: 10.1038/nrg3245. PubMed PMID: 22705668; PubMed Central PMCID: PMCPMC3551553.

9. Hadany L, Comeron JM. Why are sex and recombination so common? Ann N Y Acad Sci. 2008;1133:26–43. Epub 2008/06/19. doi: 10.1196/annals.1438.011. PubMed PMID: 18559814.

10. Crow JF. An advantage of sexual reproduction in a rapidly changing environment. J Hered. 1992;83(3):169-73. Epub 1992/05/01. doi: 10.1093/oxfordjournals.jhered.a111187. PubMed PMID: 1624761.

11. Sharp NP, Otto SP. Evolution of sex: Using experimental genomics to select among competing theories. Bioessays. 2016;38(8):751–7. Epub 2016/06/18. doi: 10.1002/bies.201600074. PubMed PMID: 27315146.

12. Immler S, Otto SP. The evolution of sex chromosomes in organisms with separate haploid sexes. Evolution. 2015;69(3):694–708. Epub 2015/01/15. doi: 10.1111/evo.12602. PubMed PMID: 25582562.

13. Hodgson EE, Otto SP. The red queen coupled with directional selection favours the evolution of sex. J Evol Biol. 2012;25(4):797–802. Epub 2012/02/11. doi: 10.1111/j.1420-9101.2012.02468.x. PubMed PMID: 22320180.

14. Otto SP, Gerstein AC. Why have sex? The population genetics of sex and recombination. Biochem Soc Trans. 2006;34(Pt 4):519–22. Epub 2006/07/22. doi: 10.1042/BST0340519. PubMed PMID: 16856849.

15. Felsenstein J. The evolutionary advantage of recombination. Genetics. 1974;78(2):737–56. Epub 1974/10/01. PubMed PMID: 4448362; PubMed Central PMCID: PMCPMC1213231.

16. Wang S, Zickler D, Kleckner N, Zhang L. Meiotic crossover patterns: Obligatory crossover, interference and homeostasis in a single process. Cell Cycle. 2015;14(3):305–14. doi: 10.4161/15384101.2014.991185.

17. Hughes SE, Hawley RS. Meiosis: Location, Location, Location, How Crossovers Ensure Segregation. Curr Biol. 2020;30(7):R311–R3. Epub 2020/04/08. doi: 10.1016/j.cub.2020.02.020. PubMed PMID: 32259504.

18. Beadle GW. A Possible Influence of the Spindle Fibre on Crossing-Over in Drosophila. Proc Natl Acad Sci U S A. 1932;18(2):160–5. doi: 10.1073/pnas.18.2.160. PubMed PMID: 16577442.

19. Bridges CB. Correspondences Between Linkage Maps and Salivary Chromosome Structure, as Illustrated in the Tip of Chromosome 2R of *Drosophila melanogaster*. CYTOLOGIA. 1937;(2):745–55. doi: 10.1508/cytologia.FujiiJubilaei.745.

20. Mather K. Crossing Over and Heterochromatin in the X Chromosome of Drosophila melanogaster. Genetics. 1939;24(3):413–35.

21. Lindsley DL, Sandler L, Counce SJ, Chandley C, Lewis KR, Riley R, et al. The genetic analysis of meiosis in female Drosophila melanogaster. Philosophical Transactions of the Royal Society of London B, Biological Sciences. 1977;277(955):295–312. doi: 10.1098/rstb.1977.0019.

22. Offermann CA, Muller HJ. Regional differences in crossing over as a function of the chromosome structure. Proc Sixth Int Congress Genet. 1932;2:143–5.

23. Sturtevant AH, Beadle GW. The Relations of Inversions in the X Chromosome of Drosophila Melanogaster to Crossing over and Disjunction. Genetics. 1936;21(5):554–604. PubMed PMID: 17246812.

24. Hawley RS. Chromosomal sites necessary for normal levels of meiotic recombination in Drosophila melanogaster. I. Evidence for and mapping of the sites. Genetics. 1980;94(3):625–46. Epub 1980/03/01. PubMed PMID: 6772522; PubMed Central PMCID: PMCPMC1214164.

25. Lambie EJ, Shirleen Roeder G. Repression of meiotic crossing over by a centromere (CEN3) in Saccharomyces cerevisiae. Genetics. 1986;114(3):769–89.

26. Lambie EJ, Shirleen Roeder G. A yeast acts in (Cis) to inhibit meiotic gene conversion of adjacent sequences. Cell. 1988;52(6):863–73. doi: 10.1016/0092-8674(88)90428-X.

27. Haupt W, Fischer TC, Winderl S, Fransz P, Torres-Ruiz RA. The CENTROMERE1 (CEN1) region of Arabidopsis thaliana: architecture and functional impact of chromatin. The Plant Journal. 2001;27(4):285–96. doi: 10.1046/j.1365-313x.2001.01087.x.

28. Copenhaver GP, Browne WE, Preuss D. Assaying genome-wide recombination and centromere functions with *Arabidopsis* tetrads. Proceedings of the National Academy of Sciences. 1998;95(1):247–52. doi: 10.1073/pnas.95.1.247.

29. Mahtani MM, Willard HF. Physical and Genetic Mapping of the Human X Chromosome Centromere: Repression of Recombination. Genome Research. 1998;8(2):100–10. doi: 10.1101/gr.8.2.100.

30. Hartmann M, Umbanhowar J, Sekelsky J. Centromere-Proximal Meiotic Crossovers in Drosophila melanogaster Are Suppressed by Both Highly Repetitive Heterochromatin and Proximity to the Centromere. Genetics. 2019;213(1):113–25. Epub 2019/07/28. doi: 10.1534/genetics.119.302509. PubMed PMID: 31345993; PubMed Central PMCID: PMCPMC6727794.

31. True JR, Mercer JM, Laurie CC. Differences in crossover frequency and distribution among three sibling species of Drosophila. Genetics. 1996;142(2):507–23. PubMed PMID: 8852849.

32. McGaugh SE, Heil CS, Manzano-Winkler B, Loewe L, Goldstein S, Himmel TL, et al. Recombination modulates how selection affects linked sites in *Drosophila*. PLoS biology. 2012;10(11):e1001422. Epub 2012/11/16. doi: 10.1371/journal.pbio.1001422. PubMed PMID: 23152720; PubMed Central PMCID: PMC3496668.

33. Stevison LS, Noor MAF. Genetic and Evolutionary Correlates of Fine-Scale Recombination Rate Variation in Drosophila persimilis. Journal of Molecular Evolution. 2010;71(5):332–45. doi: 10.1007/s00239-010-9388-1.

34. Kulathinal RJ, Bennett SM, Fitzpatrick CL, Noor MAF. Fine-scale mapping of recombination rate in Drosophila refines its correlation to diversity and divergence. Proc Natl Acad Sci U S A. 2008;105(29):10051–6. Epub 07/11. doi: 10.1073/pnas.0801848105. PubMed PMID: 18621713.

35. Sturtevant AH. The behavior of the chromosomes as studied through linkage. Zeitschrift für induktive Abstammungs- und Vererbungslehre. 1915;13(1):234–87. doi: 10.1007/bf01792906.

36. Muller HJ. The Mechanism of Crossing-Over. The American Naturalist. 1916;50(592):193–221.

37. Hatkevich T, Kohl KP, McMahan S, Hartmann MA, Williams AM, Sekelsky J. Bloom Syndrome Helicase Promotes Meiotic Crossover Patterning and Homolog Disjunction. Current Biology. 2017;27(1):96–102. doi: https://doi.org/10.1016/j.cub.2016.10.055

38. Brady MM, McMahan S, Sekelsky J. Loss of *Drosophila* Mei-41/ATR Alters Meiotic Crossover Patterning. Genetics. 2018;208(2):579–88. doi: 10.1534/genetics.117.300634.

39. Crown KN, Miller DE, Sekelsky J, Hawley RS. Local Inversion Heterozygosity Alters Recombination throughout the Genome. Current Biology. 2018;28(18):2984–90.e3. doi: https://doi.org/10.1016/j.cub.2018.07.004.

40. Page SL, Hawley RS. c(3)G encodes a Drosophila synaptonemal complex protein. Genes & Development. 2001;15(23):3130–43. doi: 10.1101/gad.935001.

41. Miller DE, Smith CB, Kazemi NY, Cockrell AJ, Arvanitakas AV, Blumenstiel JP, et al. Whole-genome analysis of individual meiotic events in *Drosophila melanogaster* reveals that noncrossover gene conversions are insensitive to interference and the centromere effect. Genetics. 2016;203(1):159–71. Epub 2016/03/06. doi: 10.1534/genetics.115.186486. PubMed PMID: 26944917; PubMed Central PMCID: PMC4858771.

42. Housworth EA, Stahl FW. Crossover interference in humans. Am J Hum Genet. 2003;73(1):188–97. Epub 05/22. doi: 10.1086/376610. PubMed PMID: 12772089.

43. Broman KW, Rowe LB, Churchill GA, Paigen K. Crossover Interference in the Mouse. Genetics. 2002;160(3):1123–31.

44. Wang Z, Shen B, Jiang J, Li J, Ma L. Effect of sex, age and genetics on crossover interference in cattle. Scientific Reports. 2016;6:37698. doi: 10.1038/srep37698

45. Broman KW, Weber JL. Characterization of Human Crossover Interference. The American Journal of Human Genetics. 2000;66(6):1911-26. doi: https://doi.org/10.1086/302923.

46. Kwiatkowski DJ, Dib C, Slaugenhaupt SA, Povey S, Gusella JF, Haines JL. An index marker map of chromosome 9 provides strong evidence for positive interference. Am J Hum Genet. 1993;53(6):1279–88. PubMed PMID: 8250044.

47. Weeks DE, Ott J, Lathrop GM. Detection of genetic interference: simulation studies and mouse data. Genetics. 1994;136(3):1217–26.

48. Anderson LK, Reeves A, Webb LM, Ashley T. Distribution of Crossing Over on Mouse Synaptonemal Complexes Using Immunofluorescent Localization of MLH1 Protein. Genetics. 1999;151(4):1569–79.

49. Berchowitz LE, Copenhaver GP. Genetic interference: don’t stand so close to me. Curr Genomics. 2010;11(2):91–102. Epub 2010/10/05. doi: 10.2174/138920210790886835. PubMed PMID: 20885817; PubMed Central PMCID: PMCPMC2874225.

50. Kohli J, Bähler J. Homologous recombination in fission yeast: Absence of crossover interference and synaptonemal complex. Experientia. 1994;50(3):295–306. doi: 10.1007/bf01924013.

51. Snow R. Maximum Likelihood Estimation of Linkage and Interference from Tetrad Data. Genetics. 1979;92(1):231–45.

52. Egel-Mitani M, Olson LW, Egel R. Meiosis in Aspergillus nidulans: Another example for lacking synaptonemal complexes in the absence of crossover interference. Hereditas. 1982;97(2):179–87. doi: 10.1111/j.1601-5223.1982.tb00870.x.

53. Sym M, Roeder GS. Crossover interference is abolished in the absence of a synaptonemal complex protein. Cell. 1994;79(2):283–92. doi: https://doi.org/10.1016/0092-8674(94)90197-X.

54. Jones GH. Chiasmata. In: Moens PB, editor. Meiosis. Orlando, FL. : Academic Press,; 1987. p. 213–44.

55. Hillers KJ, Villeneuve AM. Chromosome-Wide Control of Meiotic Crossing over in C. elegans. Current Biology. 2003;13(18):1641–7. doi: https://doi.org/10.1016/j.cub.2003.08.026.

56. Morgan TH. Sex Limited Inheritance in Drosophila. Science. 1910;32(812):120-2. Epub 1910/07/22. doi: 10.1126/science.32.812.120. PubMed PMID: 17759620.

57. Gethmann RC. Crossing Over in Males of Higher Diptera (Brachycera). J Hered. 1988;79(5):344–50. Epub 1988/09/01. doi: 10.1093/oxfordjournals.jhered.a110526. PubMed PMID: 31581763.

58. Grell R. Distributive pairing. In: Ashburner M, Novitski E, editors. The genetics and biology of Drosophila: New York: Academic Press; 1976.

59. Hawley RS, McKim KS, Arbel T. Meiotic segregation in Drosophila melanogaster females: molecules, mechanisms, and myths. Annu Rev Genet. 1993;27:281–317. Epub 1993/01/01. doi: 10.1146/annurev.ge.27.120193.001433. PubMed PMID: 8122905.

60. Dernburg AF, Sedat JW, Hawley RS. Direct evidence of a role for heterochromatin in meiotic chromosome segregation. Cell. 1996;86(1):135–46. Epub 1996/07/12. doi: 10.1016/s0092-8674(00)80084-7. PubMed PMID: 8689681.

61. Hughes SE, Gilliland WD, Cotitta JL, Takeo S, Collins KA, Hawley RS. Heterochromatic threads connect oscillating chromosomes during prometaphase I in Drosophila oocytes. PLoS Genet. 2009;5(1):e1000348. Epub 2009/01/24. doi: 10.1371/journal.pgen.1000348. PubMed PMID: 19165317; PubMed Central PMCID: PMCPMC2615114.

62. Hawley RS, Irick H, Zitron AE, Haddox DA, Lohe A, New C, et al. There are two mechanisms of achiasmate segregation in Drosophila females, one of which requires heterochromatic homology. Dev Genet. 1992;13(6):440–67. Epub 1992/01/01. doi: 10.1002/dvg.1020130608. PubMed PMID: 1304424.

63. Puro J, Nokkala S. Meiotic segregation of chromosomes in Drosophila melanogaster oocytes. Chromosoma. 1977;63(3):273–86. doi: 10.1007/bf00327454.

64. Weinstein A. The theory of multiple-strand crossing over. Genetics. 1936;21(3):155–99.

65. Weinstein A. Coincidence of Crossing Over in *Drosophila melanogaster*. Genetics. 1918;3(2):135–72.

66. Zwick ME, Salstrom JL, Langley CH. Genetic variation in rates of nondisjunction: association of two naturally occurring polymorphisms in the chromokinesin nod with increased rates of nondisjunction in Drosophila melanogaster. Genetics. 1999;152(4):1605–14. Epub 1999/08/03. PubMed PMID: 10430586; PubMed Central PMCID: PMCPMC1460721.

67. Hemmer LW, Blumenstiel JP. The Recombination Landscape of *Drosophila virilis* is Robust to Transposon Activation in Hybrid Dysgenesis. bioRxiv. 2018:342824. doi: 10.1101/342824.

68. Chino M, Kikkawa H. Mutants and Crossing over in the Dot-like Chromosome of Drosophila virilis. Genetics. 1933;18(2):111–6. PubMed PMID: 17246680.

69. Maynard Smith J, Haigh J. The hitch-hiking effect of a favorable gene. Genet Res. 1974;23:23–35.

70. Charlesworth B. Background selection and patterns of genetic diversity in *Drosophila melanogaster*. Genet Res. 1996;68(2):131–49. PubMed PMID: 8940902.

71. Charlesworth B. Fundamental concepts in genetics: effective population size and patterns of molecular evolution and variation. Nature reviews Genetics. 2009;10(3):195–205. Epub 2009/02/11. doi: 10.1038/nrg2526. PubMed PMID: 19204717.

72. Cutter AD, Payseur BA. Genomic signatures of selection at linked sites: unifying the disparity among species. Nature reviews Genetics. 2013;14(4):262-74. Epub 2013/03/13. doi: 10.1038/nrg3425. PubMed PMID: 23478346; PubMed Central PMCID: PMC4066956.

73. Comeron JM. Background selection as null hypothesis in population genomics: insights and challenges from Drosophila studies. Philos Trans R Soc Lond B Biol Sci. 2017;372(1736). Epub 2017/11/08. doi: 10.1098/rstb.2016.0471. PubMed PMID: 29109230; PubMed Central PMCID: PMCPMC5698629.

74. Gillespie JH. Genetic drift in an infinite population. The pseudohitchhiking model. Genetics. 2000;155(2):909–19. Epub 2000/06/03. PubMed PMID: 10835409; PubMed Central PMCID: PMC1461093.

75. Fisher RA. The genetical theory of natural selection. New York: Oxford University Press; 1930.

76. Muller HJ. Some genetic aspects of sex. American Naturalist. 1932;66:118–38.

77. Wiehe TH, Stephan W. Analysis of a genetic hitchhiking model, and its application to DNA polymorphism data from *Drosophila melanogaster*. Mol Biol Evol. 1993;10(4):842–54. PubMed PMID: 8355603.

78. Kim Y, Stephan W. Detecting a local signature of genetic hitchhiking along a recombining chromosome. Genetics. 2002;160(2):765–77. PubMed PMID: 11861577; PubMed Central PMCID: PMCPMC1461968.

79. Elyashiv E, Bullaughey K, Sattath S, Rinott Y, Przeworski M, Sella G. Shifts in the intensity of purifying selection: an analysis of genome-wide polymorphism data from two closely related yeast species. Genome Res. 2010;20(11):1558–73. doi: 10.1101/gr.108993.110. PubMed PMID: 20817943; PubMed Central PMCID: PMCPMC2963819.

80. Elyashiv E, Sattath S, Hu TT, Strutsovsky A, McVicker G, Andolfatto P, et al. A genomic map of the effects of linked selection in *Drosophila*. PLoS Genet. 2016;12(8):e1006130. Epub 2016/08/19. doi: 10.1371/journal.pgen.1006130. PubMed PMID: 27536991; PubMed Central PMCID: PMC4990265.

81. Nordborg M, Charlesworth B, Charlesworth D. The effect of recombination on background selection. Genet Res. 1996;67(2):159–74. Epub 1996/04/01. PubMed PMID: 8801188.

82. Charlesworth B, Morgan MT, Charlesworth D. The effect of deleterious mutations on neutral molecular variation. Genetics. 1993;134(4):1289–303. Epub 1993/08/01. PubMed PMID: 8375663; PubMed Central PMCID: PMC1205596.

83. Comeron JM. Background selection as baseline for nucleotide variation across the *Drosophila* genome. PLoS Genet. 2014;10(6):e1004434. Epub 2014/06/27. doi: 10.1371/journal.pgen.1004434. PubMed PMID: 24968283; PubMed Central PMCID: PMC4072542.

84. Sella G, Petrov DA, Przeworski M, Andolfatto P. Pervasive natural selection in the *Drosophila* genome? PLoS Genet. 2009;5(6):e1000495. Epub 2009/06/09. doi: 10.1371/journal.pgen.1000495. PubMed PMID: 19503600; PubMed Central PMCID: PMC2684638.

85. Langley CH, Stevens K, Cardeno C, Lee YC, Schrider DR, Pool JE, et al. Genomic variation in natural populations of Drosophila melanogaster. Genetics. 2012;192(2):533–98. Epub 2012/06/08. doi: 10.1534/genetics.112.142018. PubMed PMID: 22673804; PubMed Central PMCID: PMCPMC3454882.

86. Johri P, Charlesworth B, Jensen JD. Toward an Evolutionarily Appropriate Null Model: Jointly Inferring Demography and Purifying Selection. Genetics. 2020;215(1):173–92. Epub 2020/03/11. doi: 10.1534/genetics.119.303002. PubMed PMID: 32152045; PubMed Central PMCID: PMCPMC7198275.

87. Webster MT, Hurst LD. Direct and indirect consequences of meiotic recombination: implications for genome evolution. Trends Genet. 2012;28(3):101–9. Epub 2011/12/14. doi: 10.1016/j.tig.2011.11.002. PubMed PMID: 22154475.

88. Charlesworth B, Campos JL. The relations between recombination rate and patterns of molecular variation and evolution in *Drosophila*. Annu Rev Genet. 2014;48:383–403. doi: 10.1146/annurev-genet-120213-092525. PubMed PMID: 25251853.

89. Hershberg R, Petrov DA. Selection on codon bias. Annu Rev Genet. 2008;42:287–99. Epub 2008/11/06. doi: 10.1146/annurev.genet.42.110807.091442. PubMed PMID: 18983258.

90. Akashi H, Eyre-Walker A. Translational selection and molecular evolution. Curr Opin Genet Dev. 1998;8(6):688–93. Epub 1999/01/23. doi: 10.1016/s0959-437x(98)80038-5. PubMed PMID: 9914211.

91. Bulmer M. The selection-mutation-drift theory of synonymous codon usage. Genetics. 1991;129(3):897–907. Epub 1991/11/01. doi: 10.1093/genetics/129.3.897. PubMed PMID: 1752426; PubMed Central PMCID: PMCPMC1204756.

92. Duret L. Evolution of synonymous codon usage in metazoans. Curr Opin Genet Dev. 2002;12(6):640–9. Epub 2002/11/16. doi: 10.1016/s0959-437x(02)00353-2. PubMed PMID: 12433576.

93. Kliman RM, Hey J. Reduced natural selection associated with low recombination in Drosophila melanogaster. Mol Biol Evol. 1993;10(6):1239–58. Epub 1993/11/01. doi: 10.1093/oxfordjournals.molbev.a040074. PubMed PMID: 8277853.

94. Comeron JM, Kreitman M, Aguade M. Natural selection on synonymous sites is correlated with gene length and recombination in Drosophila. Genetics. 1999;151(1):239–49. Epub 1999/01/05. PubMed PMID: 9872963; PubMed Central PMCID: PMCPMC1460462.

95. Campos JL, Halligan DL, Haddrill PR, Charlesworth B. The relation between recombination rate and patterns of molecular evolution and variation in *Drosophila melanogaster*. Mol Biol Evol. 2014;31(4):1010–28. doi: 10.1093/molbev/msu056. PubMed PMID: 24489114; PubMed Central PMCID: PMCPMC3969569.

96. Hey J, Kliman RM. Interactions between natural selection, recombination and gene density in the genes of Drosophila. Genetics. 2002;160(2):595–608. Epub 2002/02/28. PubMed PMID: 11861564; PubMed Central PMCID: PMCPMC1461979.

97. Comeron JM, Kreitman M. Population, evolutionary and genomic consequences of interference selection. Genetics. 2002;161(1):389–410. Epub 2002/05/23. PubMed PMID: 12019253; PubMed Central PMCID: PMCPMC1462104.

98. Singh ND, Davis JC, Petrov DA. Codon bias and noncoding GC content correlate negatively with recombination rate on the Drosophila X chromosome. J Mol Evol. 2005;61(3):315–24. Epub 2005/07/27. doi: 10.1007/s00239-004-0287-1. PubMed PMID: 16044248.

99. Machado HE, Lawrie DS, Petrov DA. Pervasive Strong Selection at the Level of Codon Usage Bias in Drosophila melanogaster. Genetics. 2020;214(2):511–28. Epub 2019/12/25. doi: 10.1534/genetics.119.302542. PubMed PMID: 31871131; PubMed Central PMCID: PMCPMC7017021.

100. Lawrie DS, Messer PW, Hershberg R, Petrov DA. Strong purifying selection at synonymous sites in D. melanogaster. PLoS Genet. 2013;9(5):e1003527. Epub 2013/06/06. doi: 10.1371/journal.pgen.1003527. PubMed PMID: 23737754; PubMed Central PMCID: PMCPMC3667748.

101. Betancourt AJ, Presgraves DC. Linkage limits the power of natural selection in Drosophila. Proc Natl Acad Sci U S A. 2002;99(21):13616–20. Epub 2002/10/09. doi: 10.1073/pnas.212277199. PubMed PMID: 12370444; PubMed Central PMCID: PMCPMC129723.

102. Presgraves DC. Recombination enhances protein adaptation in Drosophila melanogaster. Curr Biol. 2005;15(18):1651–6. Epub 2005/09/20. doi: 10.1016/j.cub.2005.07.065. PubMed PMID: 16169487.

103. Singh ND, Larracuente AM, Clark AG. Contrasting the efficacy of selection on the X and autosomes in Drosophila. Mol Biol Evol. 2008;25(2):454–67. Epub 2007/12/18. doi: 10.1093/molbev/msm275. PubMed PMID: 18083702.

104. Larracuente AM, Sackton TB, Greenberg AJ, Wong A, Singh ND, Sturgill D, et al. Evolution of protein-coding genes in Drosophila. Trends Genet. 2008;24(3):114–23. Epub 2008/02/06. doi: 10.1016/j.tig.2007.12.001. PubMed PMID: 18249460.

105. Zhang Z, Parsch J. Positive correlation between evolutionary rate and recombination rate in Drosophila genes with male-biased expression. Mol Biol Evol. 2005;22(10):1945–7. Epub 2005/06/10. doi: 10.1093/molbev/msi189. PubMed PMID: 15944440.

106. Ewing GB, Jensen JD. The consequences of not accounting for background selection in demographic inference. Molecular ecology. 2016;25(1):135–41. Epub 2015/09/24. doi: 10.1111/mec.13390. PubMed PMID: 26394805.

107. Torres R, Stetter MG, Hernandez RD, Ross-Ibarra J. The Temporal Dynamics of Background Selection in Nonequilibrium Populations. Genetics. 2020;214(4):1019–30. Epub 2020/02/20. doi: 10.1534/genetics.119.302892. PubMed PMID: 32071195; PubMed Central PMCID: PMCPMC7153942.

108. Pavlidis P, Jensen JD, Stephan W. Searching for footprints of positive selection in whole-genome SNP data from nonequilibrium populations. Genetics. 2010;185(3):907–22. Epub 2010/04/22. doi: 10.1534/genetics.110.116459. PubMed PMID: 20407129; PubMed Central PMCID: PMCPMC2907208.

109. Kliman RM. Evidence that natural selection on codon usage in Drosophila pseudoobscura varies across codons. G3 (Bethesda). 2014;4(4):681–92. Epub 2014/02/18. doi: 10.1534/g3.114.010488. PubMed PMID: 24531731; PubMed Central PMCID: PMCPMC4059240.

110. Bartolomé C, Maside X, Charlesworth B. On the Abundance and Distribution of Transposable Elements in the Genome of Drosophila melanogaster. Molecular Biology and Evolution. 2002;19(6):926–37. doi: 10.1093/oxfordjournals.molbev.a004150.

111. Kofler R, Betancourt AJ, Schlötterer C. Sequencing of Pooled DNA Samples (Pool-Seq) Uncovers Complex Dynamics of Transposable Element Insertions in Drosophila melanogaster. PLOS Genetics. 2012;8(1):e1002487. doi: 10.1371/journal.pgen.1002487.

112. Rizzon C, Marais G, Gouy M, Biémont C. Recombination rate and the distribution of transposable elements in the Drosophila melanogaster genome. Genome research. 2002;12(3):400–7. doi: 10.1101/gr.210802. PubMed PMID: 11875027.

113. Comeron JM, Ratnappan R, Bailin S. The Many Landscapes of Recombination in *Drosophila melanogaster*. PLOS Genetics. 2012;8(10):e1002905. doi: 10.1371/journal.pgen.1002905.

114. Wang L, Bai B, Liu P, Huang SQ, Wan ZY, Chua E, et al. Construction of high-resolution recombination maps in Asian seabass. BMC Genomics. 2017;18(1):63. doi: 10.1186/s12864-016-3462-z.

115. Littrell J, Tsaih S-W, Baud A, Rastas P, Solberg-Woods L, Flister MJ. A High-Resolution Genetic Map for the Laboratory Rat. G3: Genes|Genomes|Genetics. 2018;8(7):2241–8. doi: 10.1534/g3.118.200187.

116. Wallberg A, Glémin S, Webster MT. Extreme Recombination Frequencies Shape Genome Variation and Evolution in the Honeybee, Apis mellifera. PLOS Genetics. 2015;11(4):e1005189. doi: 10.1371/journal.pgen.1005189.

117. Hemmer LW, Dias GB, Smith B, Van Vaerenberghe K, Howard A, Bergman CM, et al. Hybrid dysgenesis in Drosophila virilis results in clusters of mitotic recombination and loss-of-heterozygosity but leaves meiotic recombination unaltered. Mob DNA. 2020;11:10. Epub 2020/02/23. doi: 10.1186/s13100-020-0205-0. PubMed PMID: 32082426; PubMed Central PMCID: PMCPMC7023781.

118. Samuk K, Manzano-Winkler B, Ritz KR, Noor MAF. Natural Selection Shapes Variation in Genome-wide Recombination Rate in Drosophila pseudoobscura. Curr Biol. 2020;30(8):1517–28 e6. Epub 2020/04/11. doi: 10.1016/j.cub.2020.03.053. PubMed PMID: 32275873.

119. Mancera E, Bourgon R, Brozzi A, Huber W, Steinmetz LM. High-resolution mapping of meiotic crossovers and non-crossovers in yeast. Nature. 2008;454(7203):479–85. Epub 2008/07/11. doi: 10.1038/nature07135. PubMed PMID: 18615017; PubMed Central PMCID: PMCPMC2780006.

120. Smukowski CS, Noor MAF. Recombination rate variation in closely related species. Heredity (Edinb). 2011;107(6):496–508. Epub 06/15. doi: 10.1038/hdy.2011.44. PubMed PMID: 21673743.

121. Cheung VG, Burdick JT, Hirschmann D, Morley M. Polymorphic variation in human meiotic recombination. Am J Hum Genet. 2007;80(3):526-30. Epub 01/23. doi: 10.1086/512131. PubMed PMID: 17273974.

122. Bridges CB. A linkage variation in Drosophila. Journal of Experimental Zoology. 1915;19(1):1–21. doi: 10.1002/jez.1400190102.

123. Wang RJ, Gray MM, Parmenter MD, Broman KW, Payseur BA. Recombination rate variation in mice from an isolated island. Molecular ecology. 2017;26(2):457–70. Epub 12/21. doi: 10.1111/mec.13932. PubMed PMID: 27864900.

124. Jensen-Seaman MI, Furey TS, Payseur BA, Lu Y, Roskin KM, Chen C-F, et al. Comparative recombination rates in the rat, mouse, and human genomes. Genome research. 2004;14(4):528–38. doi: 10.1101/gr.1970304. PubMed PMID: 15059993.

125. Parvanov ED, Petkov PM, Paigen K. Prdm9 controls activation of mammalian recombination hotspots. Science. 2010;327(5967):835-. Epub 12/31. doi: 10.1126/science.1181495. PubMed PMID: 20044538.

126. Brand CL, Cattani MV, Kingan SB, Landeen EL, Presgraves DC. Molecular Evolution at a Meiosis Gene Mediates Species Differences in the Rate and Patterning of Recombination. Current Biology. 2018;28(8):1289-95.e4. doi: https://doi.org/10.1016/j.cub.2018.02.056.

127. Dumont BL, Payseur BA. Evolution of the genomic rate of recombination in mammals. Evolution. 2008;62(2):276–94. Epub 2007/12/11. doi: 10.1111/j.1558-5646.2007.00278.x. PubMed PMID: 18067567.

128. Coop G, Przeworski M. An evolutionary view of human recombination. Nature reviews Genetics. 2007;8(1):23–34. Epub 2006/12/06. doi: 10.1038/nrg1947. PubMed PMID: 17146469.

129. Ritz KR, Noor MAF, Singh ND. Variation in Recombination Rate: Adaptive or Not? Trends Genet. 2017;33(5):364–74. Epub 2017/04/01. doi: 10.1016/j.tig.2017.03.003. PubMed PMID: 28359582.

130. Stevison LS, Woerner AE, Kidd JM, Kelley JL, Veeramah KR, McManus KF, et al. The Time Scale of Recombination Rate Evolution in Great Apes. Mol Biol Evol. 2016;33(4):928–45. Epub 2015/12/17. doi: 10.1093/molbev/msv331. PubMed PMID: 26671457; PubMed Central PMCID: PMCPMC5870646.

131. Pink CJ, Swaminathan SK, Dunham I, Rogers J, Ward A, Hurst LD. Evidence that replication-associated mutation alone does not explain between-chromosome differences in substitution rates. Genome Biol Evol. 2009;1:13–22. Epub 2009/01/01. doi: 10.1093/gbe/evp001. PubMed PMID: 20333173; PubMed Central PMCID: PMCPMC2817397.

132. Patterson JT, Muller HJ. Are “Progressive” Mutations Produced by X-Rays? Genetics. 1930;15(6):495–577.

133. Hawley RS, Theurkauf WE. Requiem for distributive segregation: achiasmate segregation in Drosophila females. Trends Genet. 1993;9(9):310-7. Epub 1993/09/01. doi: 10.1016/0168-9525(93)90249-h. PubMed PMID: 8236460.

134. Lucchesi JC, Suzuki DT. The interchromosomal control of recombination. Annual Review of Genetics. 1968;2(1):53–86. doi: 10.1146/annurev.ge.02.120168.000413.

135. Stevison LS, Hoehn KB, Noor MA. Effects of inversions on within-and between-species recombination and divergence. Genome Biol Evol. 2011;3:830–41. Epub 2011/08/11. doi: 10.1093/gbe/evr081. PubMed PMID: 21828374; PubMed Central PMCID: PMCPMC3171675.

136. Lindsley DL, Zimm GG. The genome of Drosophila melanogaster. San Diego, CA: Academic Press; 1992.

137. Yamamoto M, Miklos GLG. Genetic studies on heterochromatin in Drosophila melanogaster and their implications for the functions of satellite DNA. Chromosoma. 1978;66(1):71–98. doi: 10.1007/bf00285817.

138. Yamamoto M, Gabor Miklos GL. Genetic dissection of heterochromatin in *Drosophila*: The role of basal X heterochromatin in meiotic sex chromosome behaviour. Chromosoma. 1977;60(3):283–96. doi: 10.1007/bf00329776.

139. Talbert PB, Kasinathan S, Henikoff S. Simple and Complex Centromeric Satellites in Drosophila Sibling Species. Genetics. 2018;208(3):977–90. Epub 01/05. doi: 10.1534/genetics.117.300620. PubMed PMID: 29305387.

140. Wei KH, Lower SE, Caldas IV, Sless TJS, Barbash DA, Clark AG. Variable Rates of Simple Satellite Gains across the Drosophila Phylogeny. Mol Biol Evol. 2018;35(4):925–41. Epub 2018/01/24. doi: 10.1093/molbev/msy005. PubMed PMID: 29361128; PubMed Central PMCID: PMCPMC5888958.

141. Chang C-H, Chavan A, Palladino J, Wei X, Martins NMC, Santinello B, et al. Islands of retroelements are major components of Drosophila centromeres. PLoS biology. 2019;17(5):e3000241. doi: 10.1371/journal.pbio.3000241.

142. Wei KH-C, Grenier JK, Barbash DA, Clark AG. Correlated variation and population differentiation in satellite DNA abundance among lines of *Drosophila melanogaster*. Proceedings of the National Academy of Sciences. 2014;111(52):18793–8. doi: 10.1073/pnas.1421951112.

143. Hoskins RA, Carlson JW, Wan KH, Park S, Mendez I, Galle SE, et al. The Release 6 reference sequence of the Drosophila melanogaster genome. Genome Research. 2015;25(3):445–58. doi: 10.1101/gr.185579.114.

144. McPeek MS, Speed TP. Modeling interference in genetic recombination. Genetics. 1995;139(2):1031–44. Epub 1995/02/01. doi: 10.1093/genetics/139.2.1031. PubMed PMID: 7713406; PubMed Central PMCID: PMCPMC1206354.

145. Zhao H, McPeek MS, Speed TP. Statistical analysis of chromatid interference. Genetics. 1995;139(2):1057–65. Epub 1995/02/01. PubMed PMID: 7713408; PubMed Central PMCID: PMCPMC1206356.

146. Schultz J, Redfield H. Interchromosomal effects on crossing over in *Drosophila*. Cold Spring Harb Symp Quant Biol. 1951;16:175–97. PubMed PMID: 14942738.

147. Clarke JM, Maynard Smith J. The genetics and cytology of Drosophila subobscura. XI. Hybrid vigor and longevity. J Genetics. 1955;53:172–80.

148. Maynard Smith J. Sex-limited inheritance of longevity in Drosophila subobscura. J Genetics. 1959;56:1-9.

149. Hughes KA. The inbreeding decline and average dominance of genes affecting male life-history characters in Drosophila melanogaster. Genet Res. 1995;65(1):41–52. Epub 1995/02/01. doi: 10.1017/s0016672300032997. PubMed PMID: 7750745.

150. Vermeulen CJ, Bijlsma R. Changes in mortality patterns and temperature dependence of lifespan in Drosophila melanogaster caused by inbreeding. Heredity (Edinb). 2004;92(4):275–81. Epub 2003/12/18. doi: 10.1038/sj.hdy.6800412. PubMed PMID: 14679396.

151. Charlesworth D, Charlesworth B. Inbreeding Depression and Its Evolutionary Consequences. Annu Rev Ecol Syst. 1987;18:237–68. doi: DOI 10.1146/annurev.ecolsys.18.1.237. PubMed PMID: WOS:A1987K958800011.

152. Mallet MA, Chippindale AK. Inbreeding reveals stronger net selection on Drosophila melanogaster males: implications for mutation load and the fitness of sexual females. Heredity (Edinb). 2011;106(6):994–1002. Epub 2010/12/02. doi: 10.1038/hdy.2010.148. PubMed PMID: 21119701; PubMed Central PMCID: PMCPMC3186252.

153. Perez-Pereira N, Pouso R, Rus A, Vilas A, Lopez-Cortegano E, Garcia-Dorado A, et al. Long-term exhaustion of the inbreeding load in Drosophila melanogaster. Heredity (Edinb). 2021;127(4):373–83. Epub 2021/08/18. doi: 10.1038/s41437-021-00464-3. PubMed PMID: 34400819; PubMed Central PMCID: PMCPMC8478893.

154. Sandler L, Novitski E. Meiotic Drive as an Evolutionary Force. The American Naturalist. 1957;91(857):105–10. doi: https://www.jstor.org/stable/2458433.

155. Zimmering S, Sandler L, Nicoletti B. Mechanisms of meiotic drive. Annu Rev Genet. 1970;4:409–36. Epub 1970/01/01. doi: 10.1146/annurev.ge.04.120170.002205. PubMed PMID: 4950062.

156. Courret C, Chang CH, Wei KH, Montchamp-Moreau C, Larracuente AM. Meiotic drive mechanisms: lessons from Drosophila. Proc Biol Sci. 2019;286(1913):20191430. Epub 2019/10/24. doi: 10.1098/rspb.2019.1430. PubMed PMID: 31640520; PubMed Central PMCID: PMCPMC6834043.

157. Lyttle TW. Segregation distorters. Annu Rev Genet. 1991;25:511-57. Epub 1991/01/01. doi: 10.1146/annurev.ge.25.120191.002455. PubMed PMID: 1812815.

158. Lindholm AK, Dyer KA, Firman RC, Fishman L, Forstmeier W, Holman L, et al. The Ecology and Evolutionary Dynamics of Meiotic Drive. Trends Ecol Evol. 2016;31(4):315–26. Epub 2016/02/28. doi: 10.1016/j.tree.2016.02.001. PubMed PMID: 26920473.

159. Zanders SE, Malik HS. Chromosome segregation: human female meiosis breaks all the rules. Curr Biol. 2015;25(15):R654–6. Epub 2015/08/05. doi: 10.1016/j.cub.2015.06.054. PubMed PMID: 26241139.

160. Kursel LE, Malik HS. The cellular mechanisms and consequences of centromere drive. Curr Opin Cell Biol. 2018;52:58–65. Epub 2018/02/18. doi: 10.1016/j.ceb.2018.01.011. PubMed PMID: 29454259; PubMed Central PMCID: PMCPMC5988936.

161. Brandvain Y, Coop G. Scrambling eggs: meiotic drive and the evolution of female recombination rates. Genetics. 2012;190(2):709–23. Epub 2011/12/07. doi: 10.1534/genetics.111.136721. PubMed PMID: 22143919; PubMed Central PMCID: PMCPMC3276612.

162. Bravo Nunez MA, Nuckolls NL, Zanders SE. Genetic Villains: Killer Meiotic Drivers. Trends Genet. 2018;34(6):424–33. Epub 2018/03/04. doi: 10.1016/j.tig.2018.02.003. PubMed PMID: 29499907; PubMed Central PMCID: PMCPMC5959745.

163. Larracuente AM, Presgraves DC. The selfish Segregation Distorter gene complex of Drosophila melanogaster. Genetics. 2012;192(1):33–53. Epub 2012/09/12. doi: 10.1534/genetics.112.141390. PubMed PMID: 22964836; PubMed Central PMCID: PMCPMC3430544.

164. Fishman L, Willis JH. A novel meiotic drive locus almost completely distorts segregation in mimulus (monkeyflower) hybrids. Genetics. 2005;169(1):347–53. Epub 2004/10/07. doi: 10.1534/genetics.104.032789. PubMed PMID: 15466426; PubMed Central PMCID: PMCPMC1448871.

165. Teuscher F, Brockmann GA, Rudolph PE, Swalve HH, Guiard V. Models for chromatid interference with applications to recombination data. Genetics. 2000;156(3):1449–60. Epub 2000/11/07. PubMed PMID: 11063716; PubMed Central PMCID: PMCPMC1461339.

166. Kent TV, Uzunovic J, Wright SI. Coevolution between transposable elements and recombination. Philos Trans R Soc Lond B Biol Sci. 2017;372(1736). Epub 2017/11/08. doi: 10.1098/rstb.2016.0458. PubMed PMID: 29109221; PubMed Central PMCID: PMCPMC5698620.

167. Dolgin ES, Charlesworth B. The effects of recombination rate on the distribution and abundance of transposable elements. Genetics. 2008;178(4):2169–77. Epub 2008/04/24. doi: 10.1534/genetics.107.082743. PubMed PMID: 18430942; PubMed Central PMCID: PMCPMC2323806.

168. Bergman CM, Quesneville H, Anxolabéhère D, Ashburner M. Recurrent insertion and duplication generate networks of transposable element sequences in the Drosophila melanogaster genome. Genome Biology. 2006;7(11):R112. doi: 10.1186/gb-2006-7-11-r112.

169. Kaminker JS, Bergman CM, Kronmiller B, Carlson J, Svirskas R, Patel S, et al. The transposable elements of the Drosophila melanogaster euchromatin: a genomics perspective. Genome Biology. 2002;3(12):research0084.1. doi: 10.1186/gb-2002-3-12-research0084.

170. Casals F, González J, Ruiz A. Abundance and chromosomal distribution of six Drosophila buzzatii transposons: BuT1, BuT2, BuT3, BuT4, BuT5, and BuT6. Chromosoma. 2006;115(5):403. doi: 10.1007/s00412-006-0071-7.

171. Buckley RM, Adelson DL. Mammalian genome evolution as a result of epigenetic regulation of transposable elements. Biomol Concepts. 2014;5(3):183–94. Epub 2014/11/06. doi: 10.1515/bmc-2014-0013. PubMed PMID: 25372752.

172. Slotkin RK, Martienssen R. Transposable elements and the epigenetic regulation of the genome. Nature reviews Genetics. 2007;8(4):272–85. Epub 2007/03/17. doi: 10.1038/nrg2072. PubMed PMID: 17363976.

173. Stuart T, Eichten SR, Cahn J, Karpievitch YV, Borevitz JO, Lister R. Population scale mapping of transposable element diversity reveals links to gene regulation and epigenomic variation. Elife. 2016;5. Epub 2016/12/03. doi: 10.7554/eLife.20777. PubMed PMID: 27911260; PubMed Central PMCID: PMCPMC5167521.

174. Cridland JM, Thornton KR, Long AD. Gene expression variation in Drosophila melanogaster due to rare transposable element insertion alleles of large effect. Genetics. 2015;199(1):85–93. Epub 2014/10/23. doi: 10.1534/genetics.114.170837. PubMed PMID: 25335504; PubMed Central PMCID: PMCPMC4286695.

175. Hollister JD, Gaut BS. Epigenetic silencing of transposable elements: a trade-off between reduced transposition and deleterious effects on neighboring gene expression. Genome Res. 2009;19(8):1419–28. Epub 2009/05/30. doi: 10.1101/gr.091678.109. PubMed PMID: 19478138; PubMed Central PMCID: PMCPMC2720190.

176. Lee YC. The Role of piRNA-Mediated Epigenetic Silencing in the Population Dynamics of Transposable Elements in Drosophila melanogaster. PLoS Genet. 2015;11(6):e1005269. Epub 2015/06/05. doi: 10.1371/journal.pgen.1005269. PubMed PMID: 26042931; PubMed Central PMCID: PMCPMC4456100.

177. Sentmanat MF, Elgin SC. Ectopic assembly of heterochromatin in Drosophila melanogaster triggered by transposable elements. Proc Natl Acad Sci U S A. 2012;109(35):14104–9. Epub 2012/08/15. doi: 10.1073/pnas.1207036109. PubMed PMID: 22891327; PubMed Central PMCID: PMCPMC3435190.

178. Drosophila 12 Genomes Consortium. Evolution of genes and genomes on the Drosophila phylogeny. Nature. 2007;450:203. doi: 10.1038/nature06341.

179. Locke J, Howard LT, Aippersbach N, Podemski L, Hodgetts RB. The characterization of DINE-1, a short, interspersed repetitive element present on chromosome and in the centric heterochromatin of Drosophila melanogaster. Chromosoma. 1999;108(6):356–66. Epub 1999/12/11. doi: 10.1007/s004120050387. PubMed PMID: 10591995.

180. Kapitonov VV, Jurka J. Molecular paleontology of transposable elements in the *Drosophila melanogaster* genome. Proceedings of the National Academy of Sciences. 2003;100(11):6569–74. doi: 10.1073/pnas.0732024100.

181. Petrov DA, Fiston-Lavier A-S, Lipatov M, Lenkov K, González J. Population genomics of transposable elements in Drosophila melanogaster. Molecular biology and evolution. 2011;28(5):1633–44. Epub 12/16. doi: 10.1093/molbev/msq337. PubMed PMID: 21172826.

182. Yang H-P, Barbash DA. Abundant and species-specific DINE-1 transposable elements in 12 Drosophila genomes. Genome Biology. 2008;9(2):R39. doi: 10.1186/gb-2008-9-2-r39.

183. Lu J, Clark AG. Population dynamics of PIWI-interacting RNAs (piRNAs) and their targets in Drosophila. Genome Research. 2010;20(2):212–27. doi: 10.1101/gr.095406.109.

184. Yang H-P, Hung T-L, You T-L, Yang T-H. Genomewide Comparative Analysis of the Highly Abundant Transposable Element *DINE-1* Suggests a Recent Transpositional Burst in *Drosophila yakuba*. Genetics. 2006;173(1):189–96. doi: 10.1534/genetics.105.051714.

185. Adrian AB, Corchado JC, Comeron JM. Predictive Models of Recombination Rate Variation across the *Drosophila melanogaster* Genome. Genome Biology and Evolution. 2016;8(8):2597–612. doi: 10.1093/gbe/evw181.

186. Howie JM, Mazzucco R, Taus T, Nolte V, Schlötterer C. DNA Motifs Are Not General Predictors of Recombination in Two Drosophila Sister Species. Genome Biology and Evolution. 2019;11(4):1345–57. doi: 10.1093/gbe/evz082.

187. Tamura K, Subramanian S, Kumar S. Temporal patterns of fruit fly (Drosophila) evolution revealed by mutation clocks. Mol Biol Evol. 2004;21(1):36–44. Epub 2003/09/02. doi: 10.1093/molbev/msg236. PubMed PMID: 12949132.

188. Duret L, Galtier N. Biased gene conversion and the evolution of mammalian genomic landscapes. Annu Rev Genomics Hum Genet. 2009;10:285–311. Epub 2009/07/28. doi: 10.1146/annurev-genom-082908-150001. PubMed PMID: 19630562.

189. Marais G. Biased gene conversion: implications for genome and sex evolution. Trends Genet. 2003;19(6):330–8. Epub 2003/06/13. doi: 10.1016/S0168-9525(03)00116-1. PubMed PMID: 12801726.

190. Nagylaki T. Evolution of a finite population under gene conversion. Proc Natl Acad Sci U S A. 1983;80(20):6278–81. Epub 1983/10/01. doi: 10.1073/pnas.80.20.6278. PubMed PMID: 6578508; PubMed Central PMCID: PMCPMC394279.

191. Yang Z. PAML 4: phylogenetic analysis by maximum likelihood. Mol Biol Evol. 2007;24(8):1586–91. Epub 2007/05/08. doi: 10.1093/molbev/msm088. PubMed PMID: 17483113.

192. Yang Z. PAML: a program package for phylogenetic analysis by maximum likelihood. Comput Appl Biosci. 1997;13(5):555–6. Epub 1997/11/21. doi: 10.1093/bioinformatics/13.5.555. PubMed PMID: 9367129.

193. Ortiz-Barrientos D, Chang AS, Noor MAF. A recombinational portrait of the Drosophila pseudoobscura genome. Genetical Research. 2006;87(1):23–31. Epub 03/14. doi: 10.1017/S0016672306007932.

194. Anderson WW. Linkage map of *Drosophila pseudoobscura*. In: O’Brien SJ, editor. Genetic Maps: Locus Maps of Complex Genomes. Plainview, NY: Cold Spring Harbor Laboratory Press; 1993. p. 3252–3.

195. Stapley J, Feulner PGD, Johnston SE, Santure AW, Smadja CM. Variation in recombination frequency and distribution across eukaryotes: patterns and processes. Philos Trans R Soc Lond B Biol Sci. 2017;372(1736). Epub 2017/11/08. doi: 10.1098/rstb.2016.0455. PubMed PMID: 29109219; PubMed Central PMCID: PMCPMC5698618.

196. Ottolini CS, Newnham L, Capalbo A, Natesan SA, Joshi HA, Cimadomo D, et al. Genome-wide maps of recombination and chromosome segregation in human oocytes and embryos show selection for maternal recombination rates. Nat Genet. 2015;47(7):727–35. Epub 2015/05/20. doi: 10.1038/ng.3306. PubMed PMID: 25985139; PubMed Central PMCID: PMCPMC4770575.

197. Singh ND, Criscoe DR, Skolfield S, Kohl KP, Keebaugh ES, Schlenke TA. Fruit flies diversify their offspring in response to parasite infection. Science. 2015;349(6249):747–50. Epub 2015/08/15. doi: 10.1126/science.aab1768. PubMed PMID: 26273057.

198. Dumont BL, Broman KW, Payseur BA. Variation in genomic recombination rates among heterogeneous stock mice. Genetics. 2009;182(4):1345–9. Epub 2009/06/19. doi: 10.1534/genetics.109.105114. PubMed PMID: 19535547; PubMed Central PMCID: PMCPMC2728871.

199. Kong A, Thorleifsson G, Frigge ML, Masson G, Gudbjartsson DF, Villemoes R, et al. Common and low-frequency variants associated with genome-wide recombination rate. Nat Genet. 2014;46(1):11–6. Epub 2013/11/26. doi: 10.1038/ng.2833. PubMed PMID: 24270358.

200. Hunter CM, Huang W, Mackay TF, Singh ND. The Genetic Architecture of Natural Variation in Recombination Rate in Drosophila melanogaster. PLoS Genet. 2016;12(4):e1005951. Epub 2016/04/02. doi: 10.1371/journal.pgen.1005951. PubMed PMID: 27035832; PubMed Central PMCID: PMCPMC4817973.

201. Johnston SE, Berenos C, Slate J, Pemberton JM. Conserved Genetic Architecture Underlying Individual Recombination Rate Variation in a Wild Population of Soay Sheep (Ovis aries). Genetics. 2016;203(1):583–98. Epub 2016/04/01. doi: 10.1534/genetics.115.185553. PubMed PMID: 27029733; PubMed Central PMCID: PMCPMC4858801.

202. Otto SP. The evolutionary enigma of sex. Am Nat. 2009;174 Suppl 1:S1–S14. Epub 2009/05/16. doi: 10.1086/599084. PubMed PMID: 19441962.

203. Andolfatto P, Wong KM, Bachtrog D. Effective population size and the efficacy of selection on the X chromosomes of two closely related Drosophila species. Genome Biol Evol. 2011;3:114–28. Epub 2010/12/22. doi: 10.1093/gbe/evq086. PubMed PMID: 21173424; PubMed Central PMCID: PMCPMC3038356.

204. Baines JF, Sawyer SA, Hartl DL, Parsch J. Effects of X-linkage and sex-biased gene expression on the rate of adaptive protein evolution in Drosophila. Mol Biol Evol. 2008;25(8):1639–50. Epub 2008/05/15. doi: 10.1093/molbev/msn111. PubMed PMID: 18477586; PubMed Central PMCID: PMCPMC2727381.

205. Llopart A, Brud E, Pettie N, Comeron JM. Support for the Dominance Theory in Drosophila Transcriptomes. Genetics. 2018;210(2):703–18. Epub 08/21. doi: 10.1534/genetics.118.301229. PubMed PMID: 30131345.

206. Llopart A. The rapid evolution of X-linked male-biased gene expression and the large-X effect in Drosophila yakuba, D. santomea, and their hybrids. Mol Biol Evol. 2012;29(12):3873–86. Epub 2012/07/31. doi: 10.1093/molbev/mss190. PubMed PMID: 22844069.

207. Meisel RP, Malone JH, Clark AG. Faster-X evolution of gene expression in Drosophila. PLoS Genet. 2012;8(10):e1003013. Epub 2012/10/17. doi: 10.1371/journal.pgen.1003013. PubMed PMID: 23071459; PubMed Central PMCID: PMCPMC3469423.

208. Llopart A. Parallel faster-X evolution of gene expression and protein sequences in Drosophila: beyond differences in expression properties and protein interactions. PLoS One. 2015;10(3):e0116829. Epub 2015/03/20. doi: 10.1371/journal.pone.0116829. PubMed PMID: 25789611; PubMed Central PMCID: PMCPMC4366066.

209. Llopart A. Faster-X evolution of gene expression is driven by recessive adaptive cis-regulatory variation in Drosophila. Molecular ecology. 2018;27(19):3811–21. Epub 2018/05/03. doi: 10.1111/mec.14708. PubMed PMID: 29717553.

210. Charlesworth B, Campos JL, Jackson BC. Faster-X evolution: Theory and evidence from Drosophila. Molecular ecology. 2018;27(19):3753–71. Epub 2018/02/13. doi: 10.1111/mec.14534. PubMed PMID: 29431881.

211. Kayserili MA, Gerrard DT, Tomancak P, Kalinka AT. An excess of gene expression divergence on the X chromosome in Drosophila embryos: implications for the faster-X hypothesis. PLoS Genet. 2012;8(12):e1003200. Epub 2013/01/10. doi: 10.1371/journal.pgen.1003200. PubMed PMID: 23300473; PubMed Central PMCID: PMCPMC3531489.

212. Merriam JR, Frost JN. Exchange and Nondisjunction of the X Chromosomes in Female Drosophila Melanogaster. Genetics. 1964;49:109–22. Epub 1964/01/01. PubMed PMID: 14105104; PubMed Central PMCID: PMCPMC1210549.

213. Montgomery EA, Huang SM, Langley CH, Judd BH. Chromosome rearrangement by ectopic recombination in Drosophila melanogaster: genome structure and evolution. Genetics. 1991;129(4):1085–98. Epub 1991/12/01. PubMed PMID: 1783293; PubMed Central PMCID: PMCPMC1204773.

214. Lohe AR, Roberts PA. Evolution of DNA in heterochromatin: the Drosophila melanogaster sibling species subgroup as a resource. Genetica. 2000;109(1-2):125–30. Epub 2001/04/11. doi: 10.1023/a:1026588217432. PubMed PMID: 11293787.

215. Petes TD. Meiotic recombination hot spots and cold spots. Nature Reviews Genetics. 2001;2(5):360–9. doi: 10.1038/35072078.

216. Adrian AB, Comeron JM. The *Drosophila* early ovarian transcriptome provides insight to the molecular causes of recombination rate variation across genomes. BMC Genomics. 2013;14:794. doi: 10.1186/1471-2164-14-794. PubMed PMID: 24228734; PubMed Central PMCID: PMCPMC3840681.

217. Aguilera A, Gaillard H. Transcription and recombination: when RNA meets DNA. Cold Spring Harbor perspectives in biology. 2014;6(8). Epub 2014/08/03. doi: 10.1101/cshperspect.a016543. PubMed PMID: 25085910; PubMed Central PMCID: PMCPMC4107990.

218. Aguilera A, Garcia-Muse T. R loops: from transcription byproducts to threats to genome stability. Mol Cell. 2012;46(2):115–24. Epub 2012/05/01. doi: 10.1016/j.molcel.2012.04.009. PubMed PMID: 22541554.

219. Li X, Manley JL. Cotranscriptional processes and their influence on genome stability. Genes Dev. 2006;20(14):1838–47. Epub 2006/07/19. doi: 10.1101/gad.1438306. PubMed PMID: 16847344.

220. Skourti-Stathaki K, Proudfoot NJ. A double-edged sword: R loops as threats to genome integrity and powerful regulators of gene expression. Genes Dev. 2014;28(13):1384–96. Epub 2014/07/06. doi: 10.1101/gad.242990.114. PubMed PMID: 24990962; PubMed Central PMCID: PMCPMC4083084.

221. Treco D, Arnheim N. The evolutionarily conserved repetitive sequence d(TG.AC)n promotes reciprocal exchange and generates unusual recombinant tetrads during yeast meiosis. Mol Cell Biol. 1986;6(11):3934–47. Epub 1986/11/01. doi: 10.1128/mcb.6.11.3934. PubMed PMID: 3540602; PubMed Central PMCID: PMCPMC367157.

222. Fowler KR, Sasaki M, Milman N, Keeney S, Smith GR. Evolutionarily diverse determinants of meiotic DNA break and recombination landscapes across the genome. Genome research. 2014;24(10):1650–64. doi: 10.1101/gr.172122.114. PubMed PMID: 25024163.

223. Wahba L, Costantino L, Tan FJ, Zimmer A, Koshland D. S1-DRIP-seq identifies high expression and polyA tracts as major contributors to R-loop formation. Genes Dev. 2016;30(11):1327–38. Epub 2016/06/15. doi: 10.1101/gad.280834.116. PubMed PMID: 27298336; PubMed Central PMCID: PMCPMC4911931.

224. Campos JL, Zhao L, Charlesworth B. Estimating the parameters of background selection and selective sweeps in *Drosophila* in the presence of gene conversion. Proc Natl Acad Sci U S A. 2017;114(24):E4762–E71. doi: 10.1073/pnas.1619434114. PubMed PMID: 28559322; PubMed Central PMCID: PMCPMC5474792.

225. Beissinger TM, Wang L, Crosby K, Durvasula A, Hufford MB, Ross-Ibarra J. Recent demography drives changes in linked selection across the maize genome. Nat Plants. 2016;2:16084. doi: 10.1038/nplants.2016.84. PubMed PMID: 27294617.

226. Pfeifer SP, Jensen JD. The impact of linked selection in chimpanzees: A comparative study. Genome Biol Evol. 2016;8(10):3202–8. Epub 2016/11/01. doi: 10.1093/gbe/evw240. PubMed PMID: 27678122; PubMed Central PMCID: PMC5174744.

227. Schrider DR, Kern AD. Supervised Machine Learning for Population Genetics: A New Paradigm. Trends Genet. 2018;34(4):301–12. Epub 2018/01/15. doi: 10.1016/j.tig.2017.12.005. PubMed PMID: 29331490; PubMed Central PMCID: PMCPMC5905713.

228. Schrider DR, Kern AD. S/HIC: Robust identification of soft and hard sweeps using machine learning. PLoS Genet. 2016;12(3):e1005928. Epub 2016/03/16. doi: 10.1371/journal.pgen.1005928. PubMed PMID: 26977894; PubMed Central PMCID: PMC4792382.

229. Charlesworth B. How Good Are Predictions of the Effects of Selective Sweeps on Levels of Neutral Diversity? Genetics. 2020;216(4):1217–38. Epub 2020/10/28. doi: 10.1534/genetics.120.303734. PubMed PMID: 33106248.

230. Becher H, Jackson BC, Charlesworth B. Patterns of Genetic Variability in Genomic Regions with Low Rates of Recombination. Curr Biol. 2020;30(1):94–100 e3. Epub 2019/12/24. doi: 10.1016/j.cub.2019.10.047. PubMed PMID: 31866366.

231. Comeron JM, Guthrie TB. Intragenic Hill-Robertson interference influences selection intensity on synonymous mutations in Drosophila. Mol Biol Evol. 2005;22(12):2519–30. Epub 2005/08/27. doi: 10.1093/molbev/msi246. PubMed PMID: 16120803.

232. Takano-Shimizu T. Local changes in GC/AT substitution biases and in crossover frequencies on Drosophila chromosomes. Mol Biol Evol. 2001;18(4):606–19. Epub 2001/03/27. doi: 10.1093/oxfordjournals.molbev.a003841. PubMed PMID: 11264413.

233. Takano-Shimizu T. Local recombination and mutation effects on molecular evolution in Drosophila. Genetics. 1999;153(3):1285–96. Epub 1999/11/05. PubMed PMID: 10545459; PubMed Central PMCID: PMCPMC1460815.

234. Langmead B, Salzberg SL. Fast gapped-read alignment with Bowtie 2. Nature Methods. 2012;9:357. doi: 10.1038/nmeth.1923.

235. Lunter G, Goodson M. Stampy: A statistical algorithm for sensitive and fast mapping of Illumina sequence reads. Genome Res. 2011;21(6):936–9. doi: 10.1101/gr.111120.110. PubMed PMID: PMC3106326.

236. Li H, Handsaker B, Wysoker A, Fennell T, Ruan J, Homer N, et al. The Sequence Alignment/Map format and SAMtools. Bioinformatics. 2009;25(16):2078–9. doi: 10.1093/bioinformatics/btp352. PubMed PMID: PMC2723002.

237. Thurmond J, Goodman JL, Strelets VB, Attrill H, Gramates L S, Marygold SJ, et al. FlyBase 2.0: the next generation. Nucleic Acids Research. 2018;47(D1):D759–D65. doi: 10.1093/nar/gky1003.

238. Otto SP, Payseur BA. Crossover Interference: Shedding Light on the Evolution of Recombination. Annu Rev Genet. 2019;53:19–44. Epub 2019/08/21. doi: 10.1146/annurev-genet-040119-093957. PubMed PMID: 31430178.

239. Baker BS, Carpenter ATC. Genetic analysis of sex chromosomal meiotic mutants in *Drosophila melanogaster*. Genetics. 1972;71(2):255–86.

240. Parry DM. A meiotic mutant affecting recombination in female *Drosophila melanogaster*. Genetics. 1973;73(3):465–86.

241. Stanley CE, Jr., Kulathinal RJ. flyDIVaS: A Comparative Genomics Resource for Drosophila Divergence and Selection. G3 (Bethesda). 2016;6(8):2355–63. Epub 2016/05/27. doi: 10.1534/g3.116.031138. PubMed PMID: 27226167; PubMed Central PMCID: PMCPMC4978890.

242. Benjamini Y, Hochberg Y. Controlling the false discovery rate: a practical and powerful approach to multiple testing. Journal of the Royal Statistical Society, Series B 1995;57(1):289–300. doi: 10.1016/s0166-4328(01)00297-2 PubMed Central PMCID: PMC11682119.

243. Wright F. The ’effective number of codons’ used in a gene. Gene. 1990;87(1):23–9. Epub 1990/03/01. doi: 10.1016/0378-1119(90)90491-9. PubMed PMID: 2110097.

244. Comeron JM, Aguade M. An evaluation of measures of synonymous codon usage bias. J Mol Evol. 1998;47(3):268–74. Epub 1998/09/11. doi: 10.1007/pl00006384. PubMed PMID: 9732453.

245. Tajima F. Statistical method for testing the neutral mutation hypothesis by DNA polymorphism. Genetics. 1989;123(3):585–95. Epub 1989/11/01. PubMed PMID: 2513255; PubMed Central PMCID: PMC1203831.

246. Lack JB, Cardeno CM, Crepeau MW, Taylor W, Corbett-Detig RB, Stevens KA, et al. The *Drosophila* Genome Nexus: A Population Genomic Resource of 623 *Drosophila melanogaster* Genomes, Including 197 From a Single Ancestral Range Population. Genetics. 2015;199(4):1229–41. Epub 2015/01/30. doi: 10.1534/genetics.115.174664. PubMed PMID: 25631317; PubMed Central PMCID: PMCPMC4391556.

247. Pool JE, Corbett-Detig RB, Sugino RP, Stevens KA, Cardeno CM, Crepeau MW, et al. Population genomics of sub-saharan *Drosophila melanogaster*: African diversity and non-African admixture. PLoS Genet. 2012;8(12):e1003080. Epub 2013/01/04. doi: 10.1371/journal.pgen.1003080. PubMed PMID: 23284287; PubMed Central PMCID: PMC3527209.

## References

1. Chang C-H, Chavan A, Palladino J, Wei X, Martins NMC, Santinello B, et al. Islands of retroelements are major components of Drosophila centromeres. PLoS biology. 2019;17(5):e3000241. doi: 10.1371/journal.pbio.3000241.

2. Weinstein A. The theory of multiple-strand crossing over. Genetics. 1936;21(3):155-99.

3. Miller DE, Smith CB, Kazemi NY, Cockrell AJ, Arvanitakas AV, Blumenstiel JP, et al. Whole-genome analysis of individual meiotic events in *Drosophila melanogaster* reveals that noncrossover gene conversions are insensitive to interference and the centromere effect. Genetics. 2016;203(1):159–71. Epub 2016/03/06. doi: 10.1534/genetics.115.186486. PubMed PMID: 26944917; PubMed Central PMCID: PMC4858771.

4. Hatkevich T, Kohl KP, McMahan S, Hartmann MA, Williams AM, Sekelsky J. Bloom Syndrome Helicase Promotes Meiotic Crossover Patterning and Homolog Disjunction. Current Biology. 2017;27(1):96–102. doi: https://doi.org/10.1016/j.cub.2016.10.055.

5. Baker BS, Carpenter ATC. Genetic analysis of sex chromosomal meiotic mutants in *Drosophila melanogaster*. Genetics. 1972;71(2):255–86.

6. Parry DM. A meiotic mutant affecting recombination in female *Drosophila melanogaster*. Genetics. 1973;73(3):465–86.

7. Adrian AB, Corchado JC, Comeron JM. Predictive Models of Recombination Rate Variation across the *Drosophila melanogaster* Genome. Genome Biology and Evolution. 2016;8(8):2597–612. doi: 10.1093/gbe/evw181.

## references

1. Drosophila 12 Genomes Consortium. Evolution of genes and genomes on the Drosophila phylogeny. Nature. 2007;450:203. doi: 10.1038/nature06341.

2. Ashburner M. Drosophila: A Laboratory Manual. New York: Cold Spring Harbor Laboratory Press; 1989.

3. Koren S, Walenz BP, Berlin K, Miller JR, Bergman NH, Phillippy AM. Canu: scalable and accurate long-read assembly via adaptive k-mer weighting and repeat separation. Genome Res. 2017;27(5):722–36. Epub 2017/03/17. doi: 10.1101/gr.215087.116. PubMed PMID: 28298431; PubMed Central PMCID: PMCPMC5411767.

4. English AC, Richards S, Han Y, Wang M, Vee V, Qu J, et al. Mind the Gap: Upgrading Genomes with Pacific Biosciences RS Long-Read Sequencing Technology. PLOS ONE. 2012;7(11):e47768. doi: 10.1371/journal.pone.0047768.

5. Bolger AM, Lohse M, Usadel B. Trimmomatic: a flexible trimmer for Illumina sequence data. Bioinformatics. 2014;30(15):2114–20. doi: 10.1093/bioinformatics/btu170. PubMed PMID: PMC4103590.

6. Langmead B, Salzberg SL. Fast gapped-read alignment with Bowtie 2. Nature Methods. 2012;9:357. doi: 10.1038/nmeth.1923.

7. Lunter G, Goodson M. Stampy: A statistical algorithm for sensitive and fast mapping of Illumina sequence reads. Genome Res. 2011;21(6):936–9. doi: 10.1101/gr.111120.110. PubMed PMID: PMC3106326.

8. Li H. A statistical framework for SNP calling, mutation discovery, association mapping and population genetical parameter estimation from sequencing data. Bioinformatics. 2011;27(21):2987–93. doi: 10.1093/bioinformatics/btr509. PubMed PMID: PMC3198575.

9. DePristo MA, Banks E, Poplin R, Garimella KV, Maguire JR, Hartl C, et al. A framework for variation discovery and genotyping using next-generation DNA sequencing data. Nat Genet. 2011;43(5):491–8. Epub 2011/04/12. doi: 10.1038/ng.806. PubMed PMID: 21478889; PubMed Central PMCID: PMCPMC3083463.

10. Li H, Handsaker B, Wysoker A, Fennell T, Ruan J, Homer N, et al. The Sequence Alignment/Map format and SAMtools. Bioinformatics. 2009;25(16):2078–9. doi: 10.1093/bioinformatics/btp352. PubMed PMID: PMC2723002.

11. Zhao H, Sun Z, Wang J, Huang H, Kocher JP, Wang L. CrossMap: a versatile tool for coordinate conversion between genome assemblies. Bioinformatics. 2014;30(7):1006–7. Epub 2013/12/20. doi: 10.1093/bioinformatics/btt730. PubMed PMID: 24351709; PubMed Central PMCID: PMCPMC3967108.

12. Lack JB, Cardeno CM, Crepeau MW, Taylor W, Corbett-Detig RB, Stevens KA, et al. The *Drosophila* Genome Nexus: A Population Genomic Resource of 623 *Drosophila melanogaster* Genomes, Including 197 From a Single Ancestral Range Population. Genetics. 2015;199(4):1229–41. Epub 2015/01/30. doi: 10.1534/genetics.115.174664. PubMed PMID: 25631317; PubMed Central PMCID: PMCPMC4391556.

13. Kaminker JS, Bergman CM, Kronmiller B, Carlson J, Svirskas R, Patel S, et al. The transposable elements of the Drosophila melanogaster euchromatin: a genomics perspective. Genome Biology. 2002;3(12):research0084.1. doi: 10.1186/gb-2002-3-12-research0084.

14. Ranz JM, Maurin D, Chan YS, von Grotthuss M, Hillier LW, Roote J, et al. Principles of Genome Evolution in the Drosophila melanogaster Species Group. PLoS biology. 2007;5(6):e152. doi: 10.1371/journal.pbio.0050152.

15. Lemeunier F, Ashburner M, Thoday JM. Relationships within the melanogaster species subgroup of the genus Drosophila (Sophophora) - II. Phylogenetic relationships between six species based upon polytene chromosome banding sequences. Proceedings of the Royal Society of London Series B Biological Sciences. 1976;193(1112):275–94. doi: 10.1098/rspb.1976.0046.

16. Sturtevant AH, Beadle GW. The Relations of Inversions in the X Chromosome of Drosophila Melanogaster to Crossing over and Disjunction. Genetics. 1936;21(5):554–604. PubMed PMID: 17246812.

17. Sturtevant AH. A Case of Rearrangement of Genes in Drosophila. Proc Natl Acad Sci U S A. 1921;7(8):235–7. doi: 10.1073/pnas.7.8.235. PubMed PMID: 16576597.

18. Altschul SF, Gish W, Miller W, Myers EW, Lipman DJ. Basic local alignment search tool. Journal of Molecular Biology. 1990;215(3):403–10. doi: https://doi.org/10.1016/S0022-2836(05)80360-2.

19. Morgulis A, Coulouris G, Raytselis Y, Madden TL, Agarwala R, Schäffer AA. Database indexing for production MegaBLAST searches. Bioinformatics. 2008;24(16):1757–64. doi: 10.1093/bioinformatics/btn322.

20. Kapitonov VV, Jurka J. Molecular paleontology of transposable elements in the *Drosophila melanogaster* genome. Proceedings of the National Academy of Sciences. 2003;100(11):6569–74. doi: 10.1073/pnas.0732024100.

21. Petrov DA, Fiston-Lavier A-S, Lipatov M, Lenkov K, González J. Population genomics of transposable elements in Drosophila melanogaster. Molecular biology and evolution. 2011;28(5):1633–44. Epub 12/16. doi: 10.1093/molbev/msq337. PubMed PMID: 21172826.

22. Yang H-P, Barbash DA. Abundant and species-specific DINE-1 transposable elements in 12 Drosophila genomes. Genome Biology. 2008;9(2):R39. doi: 10.1186/gb-2008-9-2-r39.

23. Lu J, Clark AG. Population dynamics of PIWI-interacting RNAs (piRNAs) and their targets in Drosophila. Genome Research. 2010;20(2):212–27. doi: 10.1101/gr.095406.109.

24. Yang H-P, Hung T-L, You T-L, Yang T-H. Genomewide Comparative Analysis of the Highly Abundant Transposable Element *DINE-1* Suggests a Recent Transpositional Burst in *Drosophila yakuba*. Genetics. 2006;173(1):189–96. doi: 10.1534/genetics.105.051714.

25. Kofler R, Hill T, Nolte V, Betancourt AJ, Schlotterer C. The recent invasion of natural Drosophila simulans populations by the P-element. Proc Natl Acad Sci U S A. 2015;112(21):6659–63. Epub 2015/05/13. doi: 10.1073/pnas.1500758112. PubMed PMID: 25964349; PubMed Central PMCID: PMCPMC4450375.

26. Kidwell MG. Evolution of hybrid dysgenesis determinants in Drosophila melanogaster. Proc Natl Acad Sci U S A. 1983;80(6):1655–9. Epub 1983/03/01. doi: 10.1073/pnas.80.6.1655. PubMed PMID: 6300863; PubMed Central PMCID: PMCPMC393661.

27. Anxolabehere D, Kidwell MG, Periquet G. Molecular characteristics of diverse populations are consistent with the hypothesis of a recent invasion of Drosophila melanogaster by mobile P elements. Mol Biol Evol. 1988;5(3):252–69. Epub 1988/05/01. doi: 10.1093/oxfordjournals.molbev.a040491. PubMed PMID: 2838720.

28. Engels WR. The origin of P elements in Drosophila melanogaster. Bioessays. 1992;14(10):681–6. Epub 1992/10/01. doi: 10.1002/bies.950141007. PubMed PMID: 1285420.

29. Daniels SB, Peterson KR, Strausbaugh LD, Kidwell MG, Chovnick A. Evidence for horizontal transmission of the P transposable element between Drosophila species. Genetics. 1990;124(2):339–55. Epub 1990/02/01. PubMed PMID: 2155157; PubMed Central PMCID: PMCPMC1203926.

30. Kelleher ES. Reexamining the *P*-Element Invasion of *Drosophila melanogaster* Through the Lens of piRNA Silencing. Genetics. 2016;203(4):1513–31. doi: 10.1534/genetics.115.184119.

31. Mohammed J, Flynt AS, Panzarino AM, Mondal MMH, DeCruz M, Siepel A, et al. Deep experimental profiling of microRNA diversity, deployment, and evolution across the Drosophila genus. Genome Res. 2018;28(1):52–65. doi: 10.1101/gr.226068.117.

32. Mohammed J, Bortolamiol-Becet D, Flynt AS, Gronau I, Siepel A, Lai EC. Adaptive evolution of testis-specific, recently evolved, clustered miRNAs in Drosophila. RNA. 2014;20(8):1195–209. doi: 10.1261/rna.044644.114.

33. Shpiz S, Ryazansky S, Olovnikov I, Abramov Y, Kalmykova A. Euchromatic Transposon Insertions Trigger Production of Novel Pi- and Endo-siRNAs at the Target Sites in the Drosophila Germline. PLOS Genetics. 2014;10(2):e1004138. doi: 10.1371/journal.pgen.1004138.

34. Kim KE, Peluso P, Babayan P, Yeadon PJ, Yu C, Fisher WW, et al. Long-read, whole-genome shotgun sequence data for five model organisms. Sci Data. 2014;1:140045-. doi: 10.1038/sdata.2014.45. PubMed PMID: 25977796.

35. Chaisson MJ, Tesler G. Mapping single molecule sequencing reads using basic local alignment with successive refinement (BLASR): application and theory. BMC Bioinformatics. 2012;13(1):238. doi: 10.1186/1471-2105-13-238.

36. Wei KH-C, Grenier JK, Barbash DA, Clark AG. Correlated variation and population differentiation in satellite DNA abundance among lines of *Drosophila melanogaster*. Proceedings of the National Academy of Sciences. 2014;111(52):18793–8. doi: 10.1073/pnas.1421951112.

37. Wei KH, Lower SE, Caldas IV, Sless TJS, Barbash DA, Clark AG. Variable Rates of Simple Satellite Gains across the Drosophila Phylogeny. Mol Biol Evol. 2018;35(4):925–41. Epub 2018/01/24. doi: 10.1093/molbev/msy005. PubMed PMID: 29361128; PubMed Central PMCID: PMCPMC5888958.

38. Benjamini Y, Hochberg Y. Controlling the false discovery rate: a practical and powerful approach to multiple testing. Journal of the Royal Statistical Society, Series B 1995;57(1):289–300. doi: 10.1016/s0166-4328(01)00297-2 PubMed Central PMCID: PMC11682119.

39. Adrian AB, Corchado JC, Comeron JM. Predictive Models of Recombination Rate Variation across the *Drosophila melanogaster* Genome. Genome Biology and Evolution. 2016;8(8):2597–612. doi: 10.1093/gbe/evw181.

40. Howie JM, Mazzucco R, Taus T, Nolte V, Schlötterer C. DNA Motifs Are Not General Predictors of Recombination in Two Drosophila Sister Species. Genome Biology and Evolution. 2019;11(4):1345–57. doi: 10.1093/gbe/evz082.

41. Comeron JM, Ratnappan R, Bailin S. The Many Landscapes of Recombination in *Drosophila melanogaster*. PLOS Genetics. 2012;8(10):e1002905. doi: 10.1371/journal.pgen.1002905.

42. Bailey TL, Boden M, Buske FA, Frith M, Grant CE, Clementi L, et al. MEME Suite: tools for motif discovery and searching. Nucleic Acids Research. 2009;37(suppl_2):W202–W8. doi: 10.1093/nar/gkp335.

